# Measuring genome sizes using read-depth, k-mers, and flow cytometry: methodological comparisons in beetles (Coleoptera)

**DOI:** 10.1101/761304

**Authors:** James M. Pflug, Valerie Renee Holmes, Crystal Burrus, J. Spencer Johnston, David R. Maddison

## Abstract

Measuring genome size across different species can yield important insights into evolution of the genome and allow for more informed decisions when designing next-generation genomic sequencing projects. New techniques for estimating genome size using shallow genomic sequence data have emerged which have the potential to augment our knowledge of genome sizes, yet these methods have only been used in a limited number of empirical studies. In this project, we compare estimation methods using next-generation sequencing (k-mer methods and average read depth of single-copy genes) to measurements from flow cytometry, the gold standard for genome size measures, using ground beetles (Carabidae) and other members of the beetle suborder Adephaga as our test system. We also present a new protocol for using read-depth of single-copy genes to estimate genome size. Additionally, we report flow cytometry measurements for five previously unmeasured carabid species, as well as 21 new draft genomes and six new draft transcriptomes across eight species of adephagan beetles. No single sequence-based method performed well on all species, and all tended to underestimate the genome sizes, although only slightly in most samples. For one species, *Bembidion haplogonum*, most sequence-based methods yielded estimates half the size suggested by flow cytometry. This discrepancy for k-mer methods can be explained by a large number of repetitive sequences, but we have no explanation for why read-depth methods yielded results that were also strikingly low.

## INTRODUCTION

The advent of modern genomics and the resulting deluge of data from next generation sequencing (NGS) has been a tremendous boon to the biological sciences. In spite of this, many foundational questions about genomes have remained largely unanswered. One such question is why genomes vary so much in size: there is an over 3,000-fold difference between the smallest and largest genomes in animals (Gregory, 2001). Revealing the myriad evolutionary causes behind this variation has proven to be a difficult and enduring challenge (Cavalier-Smith 1978, Elliott and Gregory 2015). One limitation to understanding genome size evolution is the relative lack of knowledge of genome sizes in some of the larger clades of life, such as the arthropods (Hanrahan and Johnston 2011).

Knowledge of genome size is desirable for multiple reasons. It can be important to understand phenomena such as whole genome duplication and polyploidy (Allen 1983, Némorin et al. 2013), genome reduction driven by changes in selective pressure (Johnston et al. 2004), or proliferation of non-coding DNA sequence (Gregory 2005). Genome size has also been observed to correlate with a variety of developmental factors, such as egg size and cell division rate (Gregory 2001). Knowledge about genome size can also be valuable in species delimitation. Differences sufficient to reproductively isolate a population into separate species may be difficult to distinguish using traditional morphological or DNA sequence data; however, such differences may be more apparent once genome sizes are taken into consideration alongside other evidence (Gregory 2005, Leong-Škorničková 2007).

Traditional methods for determining genome size, such as Feulgen densitometry and flow cytometry, involve staining cells with a DNA-specific dye and comparing the results to stained cells from a standard reference of a known genome size. These methods are well-tested and generally reliable (Chen et al. 2015, Hanrahan and Johnston 2011). Flow cytometry in particular is considered the “gold standard” for estimating genome size (Mounsey et al. 2012). However, these techniques rely on live, fixed, or frozen tissues with largely intact cells, effectively limiting study to organisms that can be raised in the lab or easily found in nature and transported to the lab (Hanrahan & Johnston, 2011). This can be an insurmountable problem if the specimens are difficult to collect, endangered or extinct, or only known from museum collections. NGS can potentially provide a relatively simple alternative for bioinformatically estimating genome size; however, the accuracy of these methods has not been extensively studied.

The first of these methods uses k-mer distributions. K-mers are unique subsequences of a particular length, k, from a larger DNA sequence. For example, the DNA sequence AACCTG can be decomposed into four unique k-mers that are three bases long (referred to as 3-mers): AAC, ACC, CCT, and CTG. Any set of DNA sequences, including unassembled short reads produced by NGS, can be broken down into its constituent k-mers. Each unique k-mer can be assigned a value for coverage based on the number of times it occurs in a sequence (e.g., if the 3-mer CTG is found a total of 20 times, it would have a coverage of 20). The distribution of coverages for all k-mers from a sequence can be plotted to produce a k-mer frequency distribution. For k-mers generated from genomic sequencing reads with negligible levels of sequence artifacts (sequencing errors, repeats, or coverage bias), the distribution of k-mer frequencies will approximate a Poisson distribution, with the peak centered on the average sequencing depth for the genome (Li & Waterman, 2003). The value of k varies among analyses, though values ranging between 17 and 35 are typical (Liu et al., 2013; Chen et al., 2015).

Techniques for estimating genome size using k-mer distributions generally work best when the average coverage is greater than 10X (Williams et al. 2013), but newer methods with more comprehensive models for addressing sequencing errors and repetitive sequence are showing promise at coverages as low as 0.5X (Hozza, Vinař, & Brejová, 2015). Examples of accurate k-mer based genome size estimates exist for a variety of organisms, including giant pandas (Li et al. 2010), cultivated potatoes (Potato Genome Sequencing Consortium, 2011), the agricultural pest *Bemisia tabaci* (Chen et al. 2015), and oyster (Zhang et al. 2012). However, these methods can also produce ambiguous or incorrect estimates. K-mer analysis of genomic reads from a male milkweed bug produced estimates that were 60Mb to 1110Mb higher than the approximately 930Mb flow cytometry genome estimate (Panfilio et al, 2018), with the magnitude of this overestimation increased at larger values of k. A separate study on the *Bemisia tabaci* genome (Guo et al. 2015) found that k-mer estimates of one particular biotype were about 60Mb larger than those given by flow cytometry. An alternative approach to inferring genome size from sequence data is to map NGS reads onto a set of putative single-copy genes using a reference-based assembler to determine the average coverage for the set of genes as a whole, and use that average as an estimate of coverage for the entire genome (Desvillechabrol 2016, Kanda et al. 2015).

Despite the potential value of sequence-based genome size estimation methods, little empirical research to verify them has been conducted. In most instances, these methods are incidental to the overall project and are only applied to a single individual or species. In this study, we perform three of these sequence-based genome size estimation techniques, focusing on beetles in the suborder Adephaga, and compare the results to genome size estimates derived from flow cytometry.

## MATERIALS AND METHODS

### Taxon sampling and specimen processing

Specimens were collected in Oregon and California (Tables S1, S2). Specimens assessed with flow cytometry were collected live and chilled, and their heads removed and stored at −80°C. The remaining portions of the beetles were stored in 95%-100% ethanol and retained as vouchers. Eight of these specimens were also sequenced (Table S2). No flow cytometry was performed on *Amphizoa insolens*, *Omoglymmius hamatus*, and *Trachypachus gibbsii* as sufficient numbers of specimens could not be collected at the time of the study. These three species were only assessed using sequence-based methods.

To insure sufficient low-copy-number reference sequences would be available for read mapping, six transcriptomes were also sequenced (Table S3). These transcriptomes were derived from whole-body RNA extractions from individual beetles conspecific to those used for genomic sequencing with the exception of *Lionepha casta* DNA4602, which is a close relative to *Lionepha* “Waterfalls.” Specimens used for transcriptome sequencing were either stored in RNAlater (Thermo Fisher Scientific, Waltham, MA) or kept alive until RNA extraction (Table S4). The reference transcriptome for *Amphizoa insolens* was obtained from the NCBI SRA (accession number: SRR5930489).

### DNA extraction and shallow genome sequencing

DNA was extracted from all specimens using a Qiagen DNeasy Blood and Tissue Kit (Qiagen, Hilden, Germany). DNA was extracted from muscle tissue in most cases; however, reproductive tissue (one *Lionepha* “Waterfalls” and three *Bembidion haplogonum* specimens) and, in one case (*Bembidion lividulum* DNA4146), the entire body was extracted instead. The extractions for *Lionepha* “Waterfalls” DNA5435 and *Lionepha* “Waterfalls” DNA5436 were derived from different tissues of the same specimen (reproductive and muscle, respectively) to determine if any obvious qualitative differences resulted from use of different tissue types.

The DNA was quantified using a Qubit Fluorometer (Life Technologies, Carlsbad, CA) with a Quant-iT dsDNA HS Assay Kit, and DNA fragment length distributions with a 2100 Bioanalyzer (Agilent Technologies) using the High Sensitivity DNA Analysis Kit. The samples were sheared for 10 minutes (30 seconds on, 30 seconds off) to a length of approximately 300bp using a Bioruptor Pico Sonication System (Diagenode, Denville, NJ).

Libraries were prepared using either a NEBNext DNA Ultra II kit (New England BioLabs) or an Illumina TruSeq DNA Sample Prep Kit (Table S5). Illumina sequencing was performed at the Oregon State University Center for Genomic Research and Biocomputing (OSU CGRB) on either a Hiseq 2500 or HiSeq 3000 (Table S5).

### RNA extraction and transcriptome sequencing

All transcriptomic specimens were extracted using Trizol Reagent (Thermo Fisher Scientific) and a Qiagen RNeasy Mini Kit. Whole beetles were homogenized in liquid nitrogen, with the genitalia as well as a single antenna, leg, and elytron being retained as a voucher for each individual. RNA extractions were quantified using a Qubit Fluorometer (Life Technologies) with a Qubit RNA BR Assay Kit (Thermo Fisher Scientific).

The RNA library for *Bembidion haplogonum* DNA3229 was prepared using an Illumina TruSeq RNA Sample Prep Kit at HudsonAlpha Institute for Biotechnology (Huntsville, AL). For all other specimens, mRNA was isolated using NEBNext Poly(A) mRNA Magnetic Isolation Module (New England Biolabs, Ipswich, MA), and libraries were constructed with NEBNext Ultra RNA Library Prep Kit for Illumina (New England Biolabs). The fragment size distribution of each library was characterized with a 2100 Bioanalyzer (Agilent Technologies, Santa Clara, CA) using the High Sensitivity DNA Analysis Kit and 1μl of sample.

*Bembidion haplogonum* DNA3229 was sequenced on an Illumina HiSeq 2000 at the HudsonAlpha Institute for Biotechnology. The remaining transcriptome libraries were run on either an Illumina HiSeq 2500 maintained by the OSU CGRB, or on an Illumina HiSeq 2500 at the Oregon Health and Science University’s Massively Parallel Sequencing Shared Resource (Table S4).

### Read processing and de novo assembly

Slightly different protocols were used to assemble genomes and transcriptomes. The reads of one representative of each of the eight species were imported into CLC Genomics Workbench (GW) v9.5.3 (CLC Bio-Qiagen, Aarhus, Denmark), and low quality and Illumina adapter contaminated reads were removed using the “Trim Sequences” tool with a quality limit parameter of 0.05 and an ambiguity limit of 2. *De novo* genome assembly was performed in GW using an automatic word and bubble size. Transcriptome reads were quality and adapter trimmed using the Agalma workflow (Dunn, Howison, and Zapata 2013), and then assembled using Trinity (Grabherr et al. 2011).

For genome size estimation using k-mer methods or read mapping, reads were preprocessed. To insure consistent read length for read mapping and k-mer genome size estimation, the raw reads were reprocessed using BBduk v37.62 from the BBTools package (Bushnell 2017). Reads containing Illumina adapters or other NGS artifact sequences, or with an average quality score below 10, were discarded.

The relative quality of assemblies was assessed by identifying single-copy orthologs with BUSCO v3 (Waterhouse et al. 2017) using the Endopterygota odb9 reference data set. In order to identify and quantify repetitive elements, a random sample of 500,000 read pairs was generated for each of the eight assemblies and analyzed with RepeatExplorer V2 (Novàk et al. 2013) with default parameters and using the Metazoa V3 database.

Given that mitochondrial DNA can make up a substantial portion of the DNA present in a cell (Moraes 2001), all libraries were screened for mitochondrial sequence. Mitochondrial DNA sequences were assembled for each genome using NOVOPlasty v2.7.1 (Dierckxsens 2016). NOVOPlasty produced complete circularized mitochondrial genomes for five of the eight libraries (*Amphizoa insolens* DNA3784, *Chlaenius sericeus* DNA4821, *Omoglymmius hamatus* DNA3783, *Pterostichus melanarius* DNA3787), while the remaining two libraries (*Bembidion haplogonum* DNA2544 and *Lionepha* “Waterfalls” DNA3782) produced a single large (>16,000 bp) and several small (<2,500 bp) contigs. The six complete mitochondrial genomes and the two largest contigs from DNA2544 and DNA3782 were combined in a single FASTA file. BBmap was used to map reads from each library to these mitochondrial reference sequences (minid=0.7), and the unmapped reads were used for subsequent “No Mito” read mapping analyses, which are the primary focus of this paper unless otherwise stated. Although this will also remove from consideration nuclear copies of mitochondrial DNA (“numts”) that are similar enough to the current mitochondrial genome, the fraction of reads that match mtDNA is low enough (0.14– 2.81%; see below) that removing numts can have at most a minimal effect on the estimate of genome size, especially as most of the reads that match mtDNA are presumably from the mitochondria.

### Flow cytometry

Genome size was determined following methods in Johnston, Bernardini, and Hjelmen (2019). One half of the head of each frozen adult sample was placed in ice-cold Galbraith buffer, along with the head of a female *Drosophila virilis* strain maintained in the laboratory of JSJ (1C = 328 Mb). All specimens of *Bembidion haplogonum* also contained the head of a *Periplaneta americana* (1C = 3324 Mb) to act as a second internal standard. Combined heads of the sample and standard were ground using 15 strokes of the “A” pestle in a 2 ml Kontes Dounce tissue grinder and filtered through a 20μm nylon mesh. The DNA in the nuclei released by grinding was stained for 2 hours under dark refrigeration with 25 μg/ml propidium iodide. The mean red PI fluorescence of stained nuclei was quantified using a Beckman-Coulter (Brea, CA) CytoFlex flow cytometer with a solid-state laser emitting at 488 nm. The total quantity of DNA in the sample was calculated as the ratio of red fluorescence of sample/standard (mean red fluorescence of the 2C peak of the sample divided by the mean fluorescence of the 2C peak of the standard) times the 1C amount of DNA in the standard. To increase precision, the genome size for each sample was estimated as the average from 2 technical replicates from the two halves of the head of each individual. Up to 5 individuals of each sex were scored to produce biological replicates, allowing calculation of the standard error of the mean genome size estimate. The genome size is reported as 1C, the mean amount of DNA in Mb in a haploid nucleus.

### K-mer distribution

The optimal k-mer length for genome size estimation has not been extensively tested, and the choice of k may impact the accuracy of estimates. At least one study observed that k lengths above 16 resulted in an overestimation of genome size (Panfilio et al. 2017), while another observed little variation at different lengths of k (Sun et al. 2018). The authors of GenomeScope recommend k=21 as a good tradeoff between computation speed and accuracy (Vurture *et al*. 2017), though values between 17 and 27 have been used in other studies (Zhang et al. 2015, Chen et al. 2015, Sun et al. 2018). To ensure that the length of k was not affecting the estimates, all k-mer analyses were performed with values of k ranging from 13 to 31, at steps of two, for the four species represented by multiple specimens (*Bembidion haplogonum, Chlaenius sericeus, Lionepha “*Waterfalls*”, and Pterostichus melanarius*). One-way ANOVA F-tests were performed for each combination of species and k-mer method (Figures S1–S4, Sup File 1).

The input for all k-mer based analyses in this study were plain text histograms depicting the number of k-mers at a given frequency. These were generated using the filtered Illumina reads with Jellyfish v2.0 (Marçais & Kingsford 2011), and genome sizes were estimated using two programs: GenomeScope (Vurture *et al*. 2017) and CovEST (Hozza et al. 2015). Histograms were analyzed using GenomeScope with no maximum k-mer coverage (maximum k-mer coverage = −1). GenomeScope uses an analytical model to estimate properties of a genome, including genome size, average coverage, and heterozygosity, from the distribution of k-mer frequencies. The k-mer histograms were also analyzed with CovEST using the “basic” and “repeats” models. CovEST employs a model that accounts for error rate and repeat structure, and has been demonstrated to perform well on low-coverage datasets (Hozza et al. 2015).

### Read mapping

Two separate sets of loci were selected for use as reference sequences, both containing putatively low-copy-number nuclear protein-coding genes. All loci consisted solely of protein-coding, exonic sequence to exclude introns or other non-coding sequence unexpectedly affecting read mapping. The first set contained the 74 low-copy-number nuclear genes (Supplemental File 2). This set of loci (here referred to as “Regier”) was initially selected for studying arthropod evolution (Regier *et al*. 2008), and has been used for validating NGS results in several previous studies of carabid beetles (Kanda *et al*. 2015; Sproul & Maddison 2017). The reference sequences of these genes for each species were obtained by performing a pairwise search, using Exonerate v2.4.0 (Slater & Birney 2005), of the corresponding transcriptome and the *Bembidion haplogonum* references from Kanda et al. (2015). Because no transcriptome was available for *Omoglymmius* or a near relative, reference sequences were generated directly from the assembled *Omoglymmius hamatus* DNA3783 genome.

Read mapping was repeated with a second non-overlapping set of putatively single-copy genes from OrthoDB v9.1 (Zdobnov et al. 2016); this set is here referred to as “ODB” (Supplemental File 3). Loci from six coleopteran genomes were obtained (from *Agrilus planipennis*, GCF_000699045.1; *Anoplophora glabripennis*, GCF_000390285.2; *Dendroctonus ponderosae*, GCF_000355655.1; *Leptinotarsa decemlineata*, GCF_000500325.1; *Onthophagus taurus*, GCF_000648695.1; *Tribolium castaneum*, GCF_000002335.3) by filtering for genes present in >80% of species at the Endopterygota level, resulting in a total of 135 loci (Supplemental File 4). Next, the loci were used as reference sequences for an orthologous gene search of the eight adephagan transcriptomes created here using the Orthograph software package (Petersen et al. 2017). Orthograph uses profile hidden Markov models derived from the amino acid sequences of predetermined ortholog groups from related species to identify orthologous genes in novel NGS data. These putative orthologs are then validated by a reciprocal BLAST using the same set of orthologous gene groups.

Read mapping for both data sets was performed using BBmap v37.62 with the minid parameter set to 0.7. Preprocessed reads from each of the genomic libraries were mapped to the each of the gene sets. Sequencing coverage for each library was estimated by calculating the mean coverage across all loci. The raw coverage data was generated using BBmap’s per-scaffold coverage output (using the *basecov* option). We generated plots showing coverage at each nucleotide position of the reference loci (Figures 1, S5) and observed that the coverage profile varied across each locus, with the first and last 60 to 70 bases having substantially lower coverage than the central region of the locus. Some variation in coverage is inevitable due to the stochastic nature of shotgun sequencing (Linder et al. 2013, Desvillechabrol et al. 2016); however, the decreased coverage towards the flanks is likely an artifact of the read mapping procedure, since BBMap does not map a read if only a small portion of the read overlaps with the reference sequence. This causes the average coverage of the locus to be underestimated to some degree, resulting in the average coverage estimate for all loci to be underestimated.

**Figure 1.**
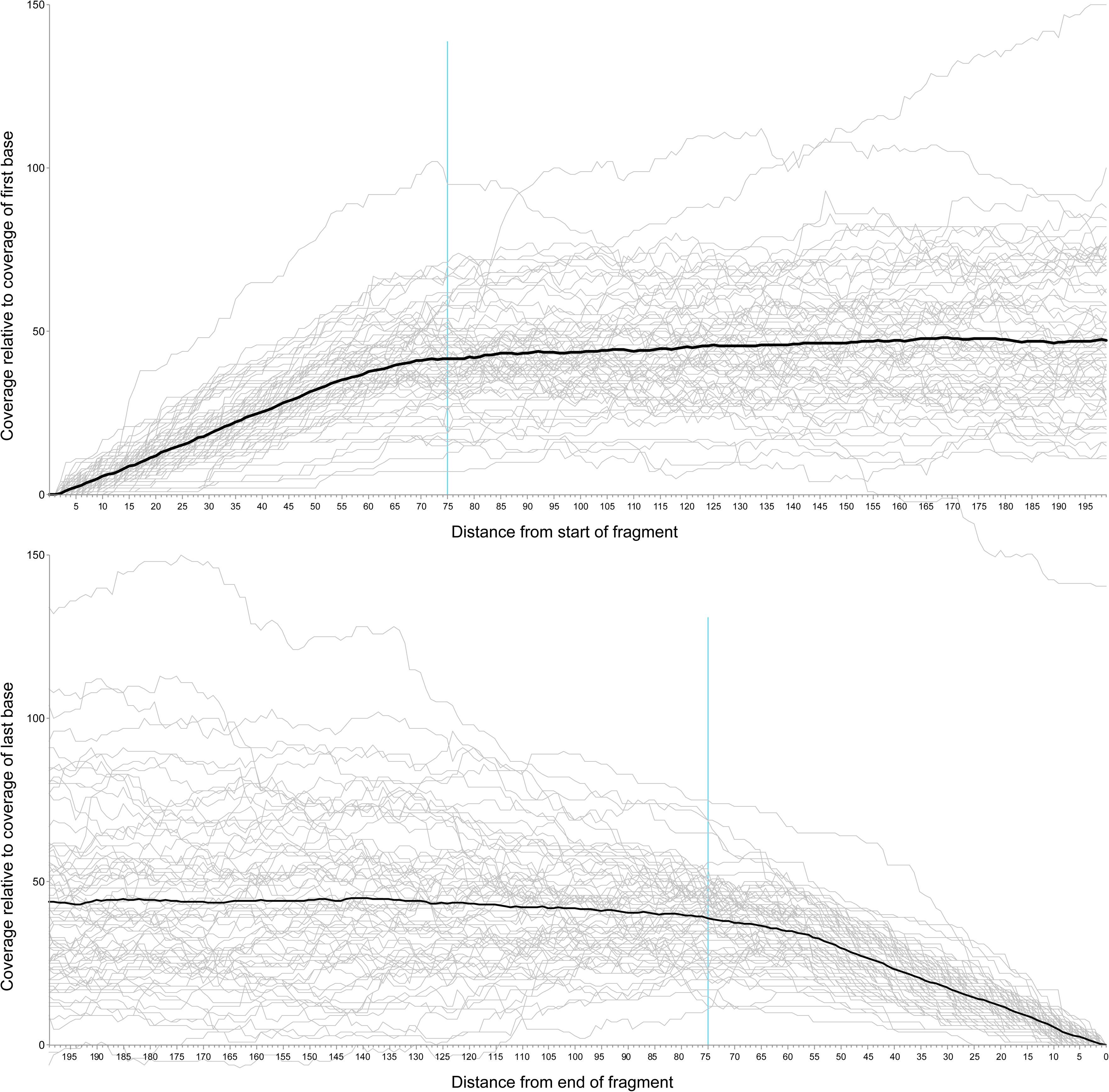
Read mapping coverage at the beginning and end of each of the Regier set loci using reads from *Bembidion haplogonum* DNA2544. The black line indicates the average relative coverage along the length of the locus, and the blue line shows 75 base positions from either end of the locus.

To compensate, a custom python script (https://github.com/JMPflug/gsec) was made to trim low coverage, flanking regions of the read mapping and produce corrected coverage estimates. The script begins with one “per-base” coverage file, containing the coverage at each base position of all loci in the reference set, for each sequenced library. This type of file can be generated by several read mapping utilities, including BBMap, Samtools, and Bedtools. The script then excludes a user-selected number of base positions at the beginning and end of each locus. The coverage profiles of the Regier set mappings for several specimens (Figures 1, S5) indicated that the drop-off in relative coverage begins at roughly 75 bases from the ends of each locus; we chose this value as the number of bases excluded by the script from each end. To prevent outlier loci from skewing the calculated means, we included loci with coverage and length values within three interquartile ranges of the median; loci with more extreme coverage or length values were removed. The script then recalculates the average per-base coverage of each locus in the reference set and uses these values to calculate the average genome coverage for the library across all loci.

These average coverage values are then used as an estimate of the average coverage of the genome, and genome size is calculated using the Lander-Waterman equation (Lander and Waterman 1988),

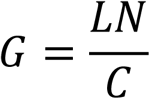

where C is the coverage estimated by read mapping, L is read length, N is the total number of reads, and G is the haploid genome length.

### Cytogenetic methods

A total of 14 males of *Bembidion haplogonum* and three males of *Lionepha* “Waterfalls” were examined for chromosome number and size; specimens came from the same localities studied for genome size. In brief, Feulgen staining followed by squashing was used; more details are given in Maddison (1985; 2008). First and second meiotic metaphase and anaphase were studied to determine chromosome number and sex chromosome system.

### Data Availability

Raw genomic reads were deposited in the Short Read Archive of NCBI’s Genbank database under accession numbers SRR8518612 to SRR8518632 (Table 1), and transcriptome reads were deposited to the NCBI SRA under accession numbers SRR8801541 to SRR8801545 (Table S3). The custom python script used is available at: https://github.com/JMPflug/gsec. All supplemental figures and tables are available on Figshare: https://TBD. The supplemental figures are:

Figures S1-S4. Genome size estimates (in Mb) from GenomeScope and CovEST at different values of K for S1. *Bembidion haplogonum*, S2. *Chlaenius sericeus*, S3. *Lionepha* “Waterfalls”, and S4. *Pterostichus melanarius*.

Figure S5. Average per-base read coverage for the first 200 bases in loci of the Regier set for four specimens.

Figure S6. Relative red fluorescence and the number of nuclei counted at each level fluorescence level of representative *Chlaenius sericeus, Lionepha “*Waterfalls*”,* and *Pterostichus melanarius* compared to a *Drosophila virilis* standard.

Figure S7. Percent of reads inferred to contain repetitive elements as inferred by RepeatExplorer from a sample of 500,000 read pairs for each of eight beetle species, with reads classified to major group of repetitive elements.

The supplemental tables are:

Table S1. Information on genomic specimens sequenced.

Table S2. Flow Cytometry values for specimens examined.

Table S3. Information on transcriptomic specimens sequenced.

Table S4. Additional details for methods used on transcriptomic sequencing specimens.

Table S5. Additional details for methods used on genomic sequencing specimens.

Table S6. Results of BUSCO analysis on eight genome assemblies using the 2442 gene Endopterygota odb9 reference set.

Table S7. Results of BUSCO analysis on six transcriptome assemblies using the 2442 gene Endopterygota odb9 reference set.

Table S8. GenomeScope results.

Table S9. Summary of read mapping genome size estimates for the Regier and OrthoDB gene sets using three different read filtering methods.

Table S10. CovEST genome size (in Mb) and coverage estimates for two models, Basic and Repeat, performed using a k value of 21.

Table S11. Regier read mapping mean coverages before and after removing outliers using the 3*IQR rule.

Table S12. OrthoDB read mapping mean coverages before (“Untrimmed”) and after (“IQR Trim”) removing outliers more than three interquartiles from the median.

Table S13. Comparison of total repetitive DNA sequence (in Mb) estimated by RepeatExplorer and GenomeScope.

Table S14. GenomeScope genome size estimates using *Drosophila melanogaster* embryo, salivary gland, and ovarian cell data set from Yarosh and Spradling (2014).

## RESULTS

### Flow cytometry

The genome sizes of the five species of carabid beetles studied (Tables 1, S2) vary over a five-fold range. *Bembidion haplogonum* possessed the largest genome, with estimates of 1C = 2,193.4♂ / 2,118.1♀Mb (Figure 2), while the genome of *Chlaenius sericeus* was the smallest with 1C = 408.4♂ / 391.5♀Mb (Figure S6). The genome of the former is over twice the size of the largest previously measured carabid beetle, *Calosoma scrutator,* which was estimated to be 1C = 1,017.1Mb (Hanrahan and Johnston 2011), and ranks as the 8th largest beetle genome out of the nearly 300 analyzed Coleoptera in the Animal Genome Size Database (Gregory 2001).

**Table 1.** Average flow cytometry genome size measurements. Values given in Mb. SD indicates standard deviation; SE indicates standard error.

|  | Female |  |  |  | Male |  |  |  |
| --- | --- | --- | --- | --- | --- | --- | --- | --- |
|  | Genome Size | N | SD | SE | Genome Size | N | SD | SE |
| *Bembidion haplogonum* | 2193.41 | 9 | 26.26 | 8.75 | 2118.05 | 9 | 27.77 | 9.26 |
| *Bembidion lividulum* | 837.39 | 1 | - | - | 831.70 | 5 | 6.57 | 2.94 |
| *Chlaenius sericeus* | 408.37 | 6 | 12.43 | 5.08 | 391.50 | 4 | 4.01 | 2.00 |
| *Lionepha "Waterfall"* | 597.60 | 1 | - | - | 585.70 | 5 | 5.41 | 2.42 |
| *Pterostichus melanarius* | 1040.65 | 4 | 18.42 | 9.21 | 1000.93 | 4 | 19.16 | 9.58 |

**Figure 2.**
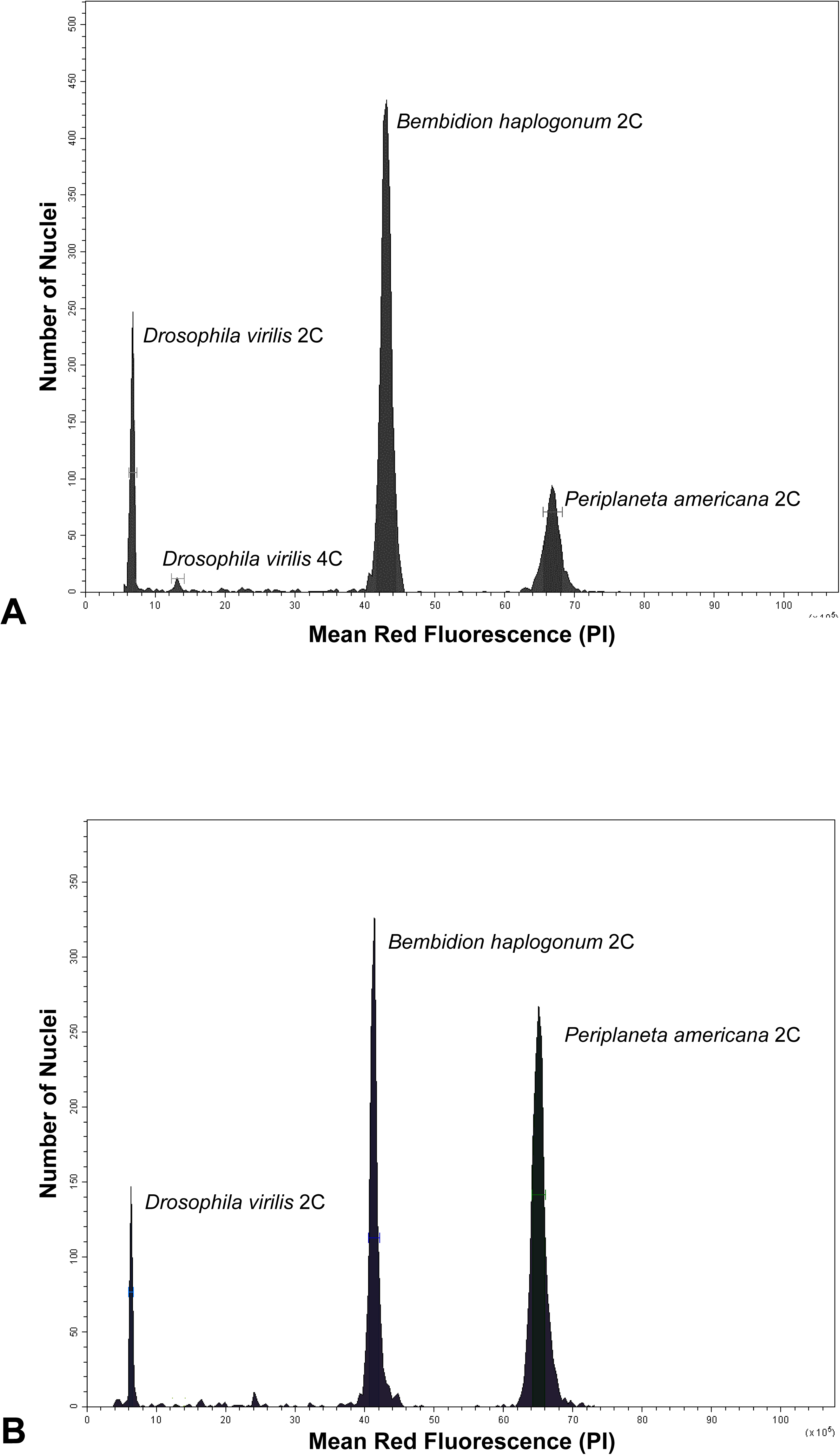
Relative red fluorescence and the number of nuclei counted at each fluorescence level of representative male (A) and female (B) *Bembidion haplogonum*. Bars around each peak represent statistical gates that provide the total nuclei in that peak, average channel number of nuclei in the peak, and the coefficient of variation (CV). *D. virilis* standard 1C=328 Mb, *P. americana* standard 1C=3,338 Mb.

### Sequencing and Assembly

Approximately 3.5 billion reads were generated across all 21 genomes, resulting in a total 473.6 billion bases. These were used to create draft assemblies of the eight species of adephagan beetles studied. The total assembled scaffold lengths varied between 153.1Mb for *Omoglymmius hamatus* DNA3783, and 665.1Mb for *Bembidion haplogonum* DNA2544 (Table 2). The assembly lengths are substantially smaller than the genome sizes inferred by both sequence-based and flow cytometric methods, indicating that large portions of the genomes could not be assembled. The number of genes in each assembly identified by BUSCO as “complete” (i.e., a putative orthologous gene found to similar to one of the 2442 BUSCO gene groups, and whose length is within two standard deviations of the mean length of the genes in that BUSCO group) varied greatly across the genome assemblies (Table S6). 1,923 (78.7%) of the 2442 genes were completely found in *Chlaenius sericeus* DNA4821, while *Omoglymmius hamatus* DNA3783 contained 178 (7.3%) complete genes.

**Table 2.** Summary of results for genomes assembled with CLC Genomics Workbench.

|  | Read Length | Total Reads | Reads After Trim | N50 | Average Contig (bp) | Maximum Contig (bp) | Contig Count | Total Assembled Bases |
| --- | --- | --- | --- | --- | --- | --- | --- | --- |
| *Amphizoa insolens* DNA3784 | 101 | 59,590,984 | 59,182,421 | 355 | 299 | 79,486 | 905,365 | 270,465,134 |
| *Bembidion haplogonum* DNA2544 | 151 | 717,036,094 | 713,035,405 | 525 | 460 | 140,578 | 1,438,922 | 661,385,315 |
| *Bembidion lividulum* DNA4161 | 101 | 55,816,204 | 51,442,737 | 673 | 543 | 16,894 | 280,998 | 152,578,104 |
| *Chlaenius sericeus* DNA4821 | 151 | 77,809,004 | 75,284,196 | 1,411 | 893 | 68,744 | 340,400 | 304,113,552 |
| *Lionepha* "Waterfalls" DNA3782 | 101 | 80,214,318 | 79,962,073 | 4,114 | 1,481 | 47,683 | 124,458 | 184,268,766 |
| *Omoglymmius hamatus* DNA3783 | 101 | 80,209,476 | 79,995,111 | 366 | 306 | 13,857 | 1,298,649 | 397,294,330 |
| *Pterostichus melanarius* DNA3787 | 101 | 69,486,602 | 69,476,201 | 341 | 289 | 17,107 | 1,160,880 | 335,949,421 |
| *Trachypachus gibbsii* DNA3786 | 101 | 88,000,000 | 86,431,416 | 1,258 | 588 | 65,825 | 472,429 | 277,787,110 |

A total of over 570 million reads were generated from the six transcriptomes (Table 3). The number of transcripts assembled ranged from 22,330 with *Bembidion lividulum* DNA4279 to 57,119 with *Bembidion haplogonum* DNA3229. As anticipated, BUSCO was able to locate many more genes in the transcriptomic assemblies (Table S7).

**Table 3.** Summary of results for transcriptomes assembled with Trinity.

|  | Read Length | Reads Examined | Transcripts | N50 | Average contig length | Total assembled bases |
| --- | --- | --- | --- | --- | --- | --- |
| *Bembidion haplogonum* DNA3229 | 50 | 425,577,514 | 57,119 | 1274 | 898.66 | 35,767,428 |
| *Bembidion lividulum* DNA4279 | 101 | 22,400,532 | 22,330 | 1727 | 1177.05 | 20,375,906 |
| *Chlaenius sericeus* JMPR007 | 101 | 22,103,790 | 24,327 | 1764 | 1146.54 | 22,566,122 |
| *Lionepha casta* DNA4602 | 100 | 30,791,800 | 26,994 | 1963 | 1338.76 | 23,724,243 |
| *Pterostichus melanarius* DNA4765 | 100 | 38,061,712 | 34,153 | 2184 | 1338.91 | 30,509,665 |
| *Trachypachus gibbsii* DNA4436 | 101 | 31,571,972 | 29,159 | 1823 | 1209.21 | 26,260,335 |

RepeatExplorer classified between 23.9% and 66.6% of the sampled reads as originating from repetitive elements (Figure S7). Three samples, all from the carabid tribe Bembidiini, consisted of over 50% repetitive DNA: *Bembidion haplogonum* DNA2544, *Lionepha* “Waterfalls” DNA3782, and *Bembidion lividulum* DNA4161. Although a variety of repeat families were identified, including Ty3/Gypsy, Ty1/Copia, Penelope, and LINEs, RepeatExplorer was unable to classify the overwhelming majority of these repetitive sequences (70.1% to 99.3%). 10.3% of the *Bembidion lividulum* DNA4161 reads were classified as ribosomal DNA, a finding consistent with other studies on this species (Sproul and Maddison 2017). The genomic libraries consisted of between 0.14% to 2.81% mitochondrial DNA.

### K-mer distribution

The estimated genome size changed with k-mer length for both CovEST models, but not for GenomeScope. The length of k had a significant effect on estimated genome size in all four species (one-way ANOVA, p-value < 0.001) for both “repeat” and “basic” models. A non-significant difference in GenomeScope genome size estimates was observed at varying sizes of k for all species (one-way ANOVA, p-value > 0.40).

Post-hoc Tukey-Kramer tests of the CovEST results revealed that estimates using a k of less than 17 differed significantly (p-value < 0.05) from those using k values above 21 for all species (Figures S1–S4, Sup. File 1). Values of k less than 17 always yielded much lower genome size estimates than suggested by both flow cytometry and read mapping, while larger k values (19 to 31) produced estimates that were more consistent with the results of other estimation methods. Given these observations, and as the computational expense of generating k-mer graphs increases as k becomes larger (Marçais & Kingsford 2011), a k value of 21 was selected as the standard k value for this study. All subsequent discussion about k-mer based estimates in this study, including tables and figures, refer to analyses performed using this value unless otherwise stated.

GenomeScope converged and produced genome size estimates for 12 of the 21 specimens (Tables S6, S8). GenomeScope failed to converge for *Amphizoa insolens* DNA3784, *Bembidion lividulum* DNA4161, and four of the five *Pterostichus melanarius* samples (JMP059, JMP060, JMP061, and JMP062). *Omoglymmius hamatus* DNA3783 and *Pterostichus melanarius* DNA3787 did converge but yielded implausibly low genome size estimates (74Mb and 74.6MB, respectively). The specimens that failed to converge lacked obvious coverage peaks in their k-mer histograms, suggesting they did not have sufficient coverage for the GenomeScope model. Coverage estimates of these specimens using read mapping provide support for this idea, as all specimens that did not converge had an estimated coverage less than 26X, while only one specimen whose coverage was less than 26X, *Lionepha* “Waterfalls” DNA3782, converged (Tables S9, S10). A similar situation was observed for both CovEST models. All non-converging specimens were estimated to have a coverage less than 13X for the “basic” model and less than 22X for the “repeat” model, with *Lionepha* “Waterfalls” DNA3782 again being the only specimen to converge with a coverage below this level. This result is consistent the Vurture et al. (2017)’s recommended minimum coverage of at least 25X coverage for an accurate estimate. The successful analyses produced genome size estimates ranging from 1,113.7Mb for *Bembidion haplogonum* DNA2544 to 264.0Mb for *Trachypachus gibbsii* DNA3786.

CovEST produced estimates for all 21 samples (Tables S6, S10), though the two models behaved differently. The “repeats” model yielded estimates approximately double the size of the “basic” model estimates, and the “repeat” model always produced the largest estimate among all the sequence-based methods.

### Read Mapping

There was no statistically significant difference between the estimates using the Regier and ODB gene sets (two-sample t-test, p=0.1611). Removing mitochondrial reads resulted, on average, in a modest increase in genome size estimates for both Regier (1.31%, s = 0.75%) and ODB (1.2%, s = 0.66%) gene sets (Table S9).

In most samples, we observed that a small number of loci with notably high coverage was removed by the gsec script. In all but one instance, excluding these loci had a minimal impact on the average coverage (Tables S11, S12). The single exception, *Omoglymmius hamatus* DNA3783, contained three Regier set loci with substantially higher coverage (>160X) than the rest of the set (Figure 3), which ranged between 2.10X to 9.27X. The inclusion of these three loci nearly doubled the estimated genome size. The high-coverage of these loci suggests that all or part of these genes are multi-copy in this species, or their coverage is being inflated by extraneous reads from an undetected source such as a pseudogene or contaminant DNA.

**Figure 3.**
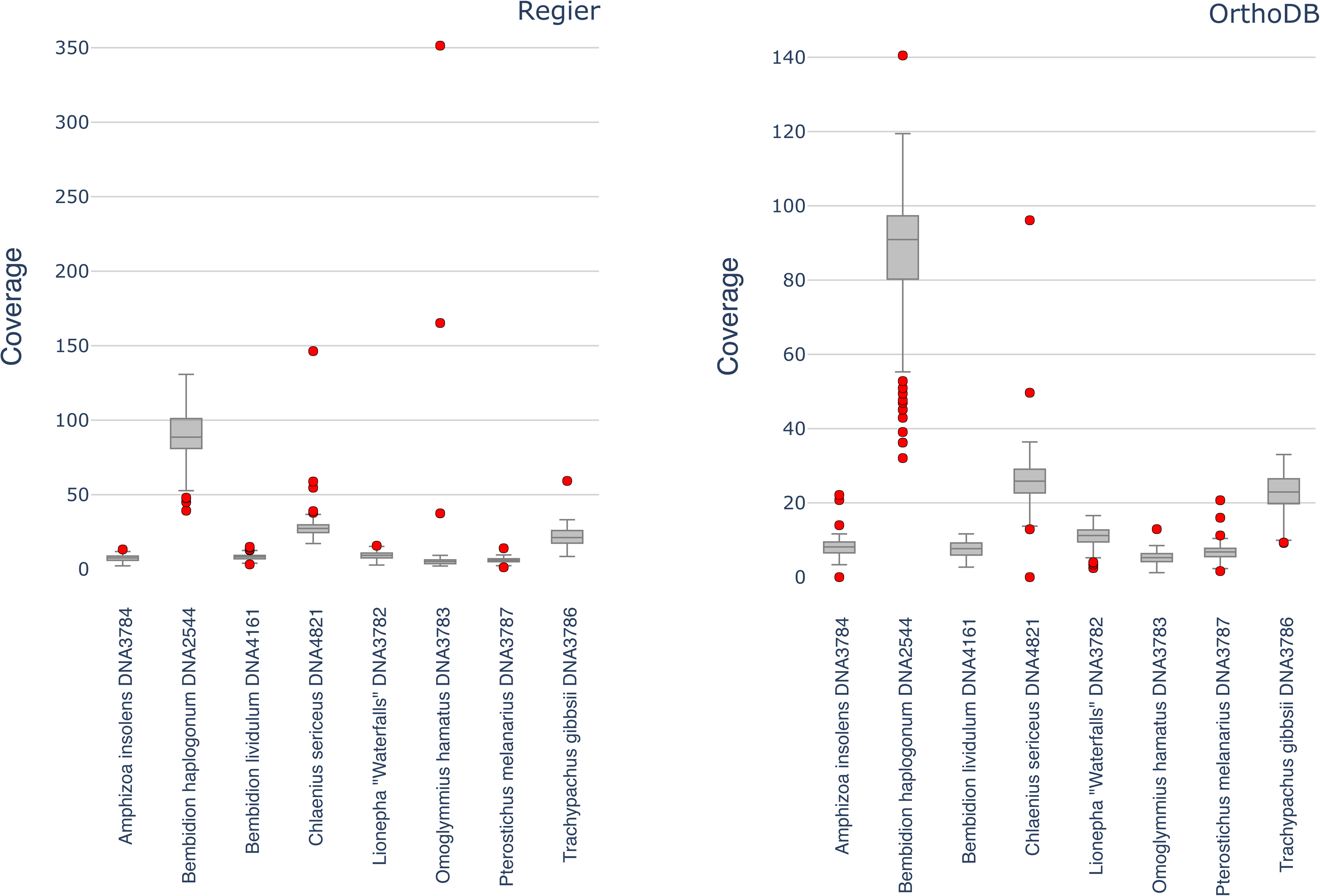
Boxplot of average genomic read mapping coverages for each of the Regier (A) and OrthoDB (B) genes for eight representative specimens. Red dots indicate outlier genes with coverage outside three interquartiles from the median.

### Comparison of Flow Cytometry and Sequence-based Genome Size Estimates

Accuracy of the sequence-based genome size estimation methods depended greatly on the species being analyzed (Figures 4, 5, Table 4, S6). In most cases, sequence-based methods underestimated size of the genome as measured by flow cytometry. In addition, estimates produced by the various sequence-based methods sometimes differed from each other. The largest discrepancy among methods was observed with the large genome of *Bembidion haplogonum*. The average flow cytometry measurement of male *B. haplogonum* was 2,118.1Mb; however, read mapping, GenomeScope, and the “basic” CovEST model estimated the genome to be approximately half that size (932.0Mb to 1134.1Mb). A similar, though less pronounced, pattern was observed with the next largest genome, of *Pterostichus melanarius*. In contrast, the CovEST “repeats” estimate was within 10% of the flow cytometric value in three of the four *B. haplogonum* samples (Table S6). For *Chlaenius sericeus*: read mapping, GenomeScope, and CovEST “basic” underestimated genome size by a modest 4.50% on average, while the CovEST “repeats” model produced the only overestimate, inflating the genome by an average of 78.0%. Sequence-based estimates of *Lionepha* “Waterfalls” varied depending on the method but were generally lower than the flow cytometric value.

**Figure 4.**
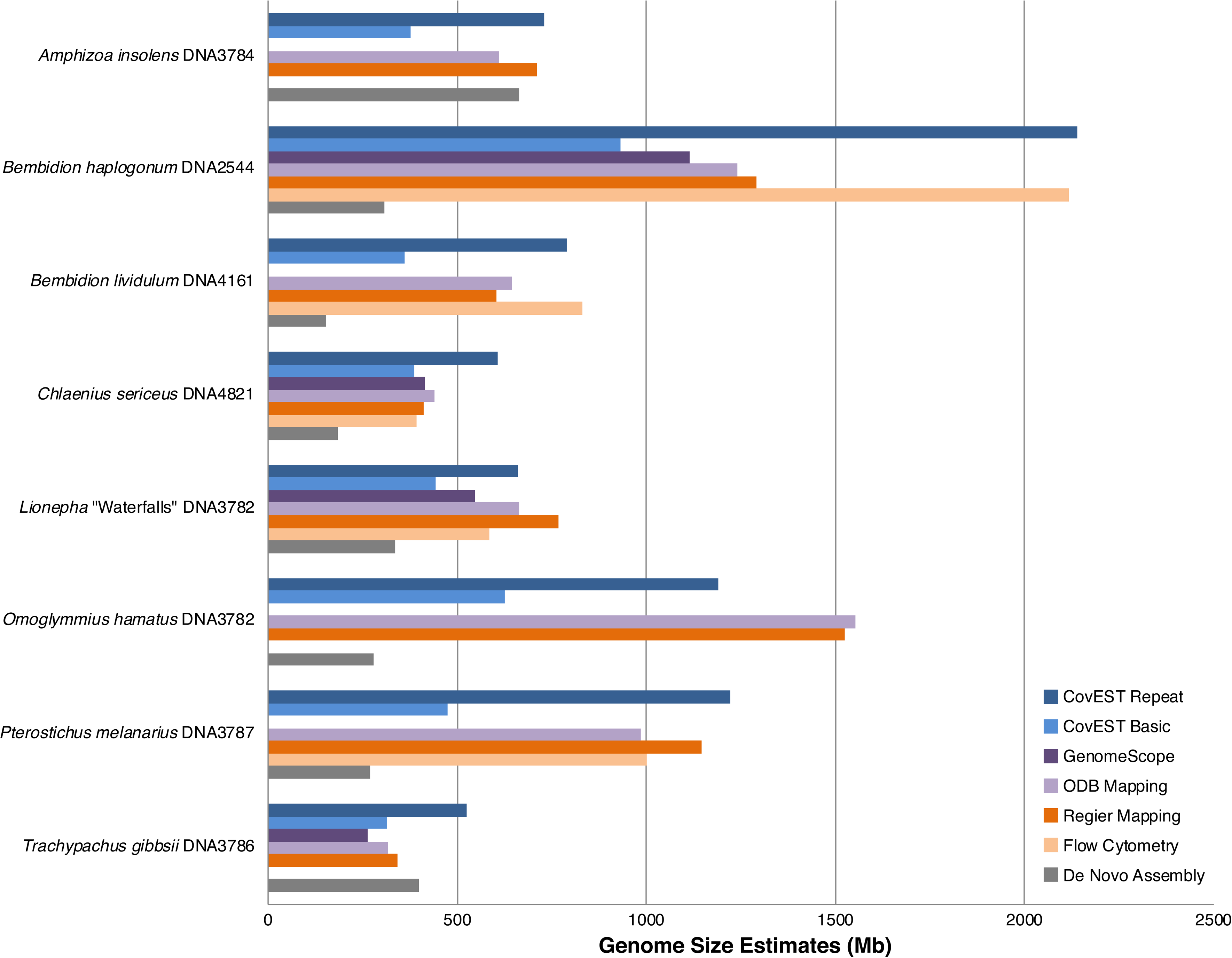
Summary of genome size estimates using flow cytometry and sequence-based methods for the eight adephagan species. Flow cytometry measurements are averages of multiple individuals (see Tables TX02 and S3). CovEST Basic, CovEST Repeat, and GenomeScope analyses were conducted using a k value of 21. Sequence-based estimates were obtained from different individual specimens (Table TX01) than those analyzed with flow cytometry.

**Figure 5.**
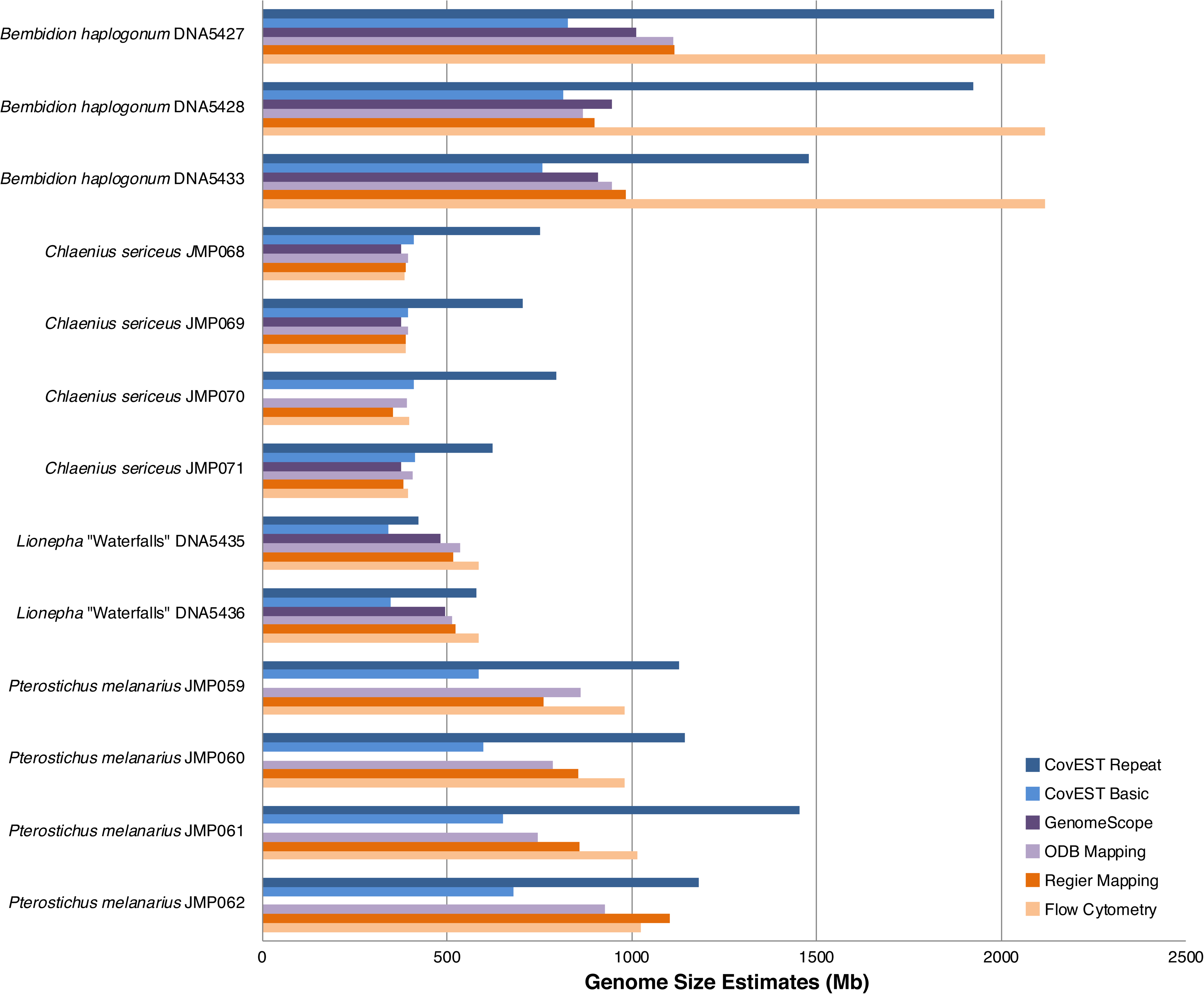
Summary of genome size estimates using flow cytometry and sequence-based methods for the 13 samples. CovEST Basic, CovEST Repeat, and GenomeScope analyses were conducted using a k value of 21. a. Samples made with DNA extracted from different tissues of the same individual.

**Table 4.** Summary of genome size estimates using flow cytometry and sequence-based methods. Values given in Mb. Flow cytometry Flow cytometry was not performed on *Trachypachus gibbsii* DNA3786, *Amphizoa insolens* DNA3784, and *Omoglymmius hamatus* DNA3782. CovEST Basic, CovEST Repeat, and GenomeScope analyses were conducted using a k value of 21. Cells in the GenomeScope column containing dashes indicate that sample failed to converge.

|  | Flow Cytometry | Regier Mapping | ODB Mapping | GenomeScope | CovEST Basic | CovEST Repeat |
| --- | --- | --- | --- | --- | --- | --- |
| *Amphizoa insolens* DNA3784 | - | 710.1 | 610.7 | - | 376.78 | 728.47 |
| *Bembidion haplogonum* DNA2544 | 2118.1<sup>b</sup> | 1,291.1 | 1,241.0 | 1113.67 | 932.04 | 2140.04 |
| *Bembidion haplogonum* DNA5427 | 2118.1<sup>b</sup> | 1,114.9 | 1,113.4 | 1010.67 | 827.88 | 1980.24 |
| *Bembidion haplogonum* DNA5428 | 2118.1<sup>b</sup> | 897.9 | 866.4 | 945.51 | 813.44 | 1924.27 |
| *Bembidion haplogonum* DNA5433 | 2118.1<sup>b</sup> | 983.3 | 946.4 | 908.41 | 758.09 | 1480.15 |
| *Bembidion lividulum* DNA4161 | 831.7<sup>b</sup> | 603.1 | 645.8 | - | 359.94 | 790.81 |
| *Chlaenius sericeus* DNA4821 | 391.5 | 411.5 | 438.5 | 414.46 | 386.92 | 608.44 |
| *Chlaenius sericeus* JMP068 | 385.8 | 390.1 | 395.7 | 374.46 | 410.91 | 751.89 |
| *Chlaenius sericeus* JMP069 | 390.1 | 389.5 | 393.4 | 374.67 | 395.86 | 704.51 |
| *Chlaenius sericeus* JMP070 | 396.5 | 354.5 | 393.1 | - | 409.81 | 796.39 |
| *Chlaenius sericeus* JMP071 | 393.6 | 383.1 | 406.7 | 377.09 | 412.78 | 624.53 |
| *Lionepha* "Waterfalls" DNA3782 | 585.7<sup>b</sup> | 769.1 | 663.1 | 545.72 | 442.32 | 659.97 |
| *Lionepha* "Waterfalls" DNA5435<sup>a</sup> | 585.7<sup>b</sup> | 518.1 | 535.9 | 483.39 | 340.38 | 423.66 |
| *Lionepha* "Waterfalls" DNA5436<sup>a</sup> | 585.7<sup>b</sup> | 522.1 | 513.9 | 493.63 | 346.52 | 578.96 |
| *Omoglymmius hamatus* DNA3783 | - | 1,525.7 | 1,552.1 | - | 627.13 | 1188.94 |
| *Pterostichus melanarius* DNA3787 | 1000.9 | 1,144.9 | 984.6 | - | 475.11 | 1221.83 |
| *Pterostichus melanarius* JMP059 | 1045.1 | 760.3 | 860.5 | - | 586.67 | 1127.86 |
| *Pterostichus melanarius* JMP060 | 1018.3 | 855.5 | 787.4 | - | 597.29 | 1145.04 |
| *Pterostichus melanarius* JMP061 | 1068.0 | 859.9 | 746.4 | - | 650.22 | 1454.54 |
| *Pterostichus melanarius* JMP062 | 1031.2 | 1,103.7 | 927.6 | - | 678.61 | 1181.38 |
| *Trachypachus gibbsii* DNA3786 | - | 342.0 | 315.5 | 264.08 | 314.81 | 525.13 |

a. Samples made with DNA extracted from different tissues of the same individual.
b. Flow cytometry measurements for sample are species averages of multiple individuals (see Tables 2 and S3).

### Cytogenetic results

The chromosomes of *Bembidion haplogonum* are visibly larger than those of *Lionepha* “Waterfalls” (Figure 6), which agrees with both flow cytometry and sequence-based estimates. Males of *Bembidion haplogonum* have 11 pairs of autosomes and an XY pair of sex chromosomes; males of *Lionepha* “Waterfalls” have 12 pairs of autosomes and have a single X chromosome with no Y chromosome.

**Figure 6.**
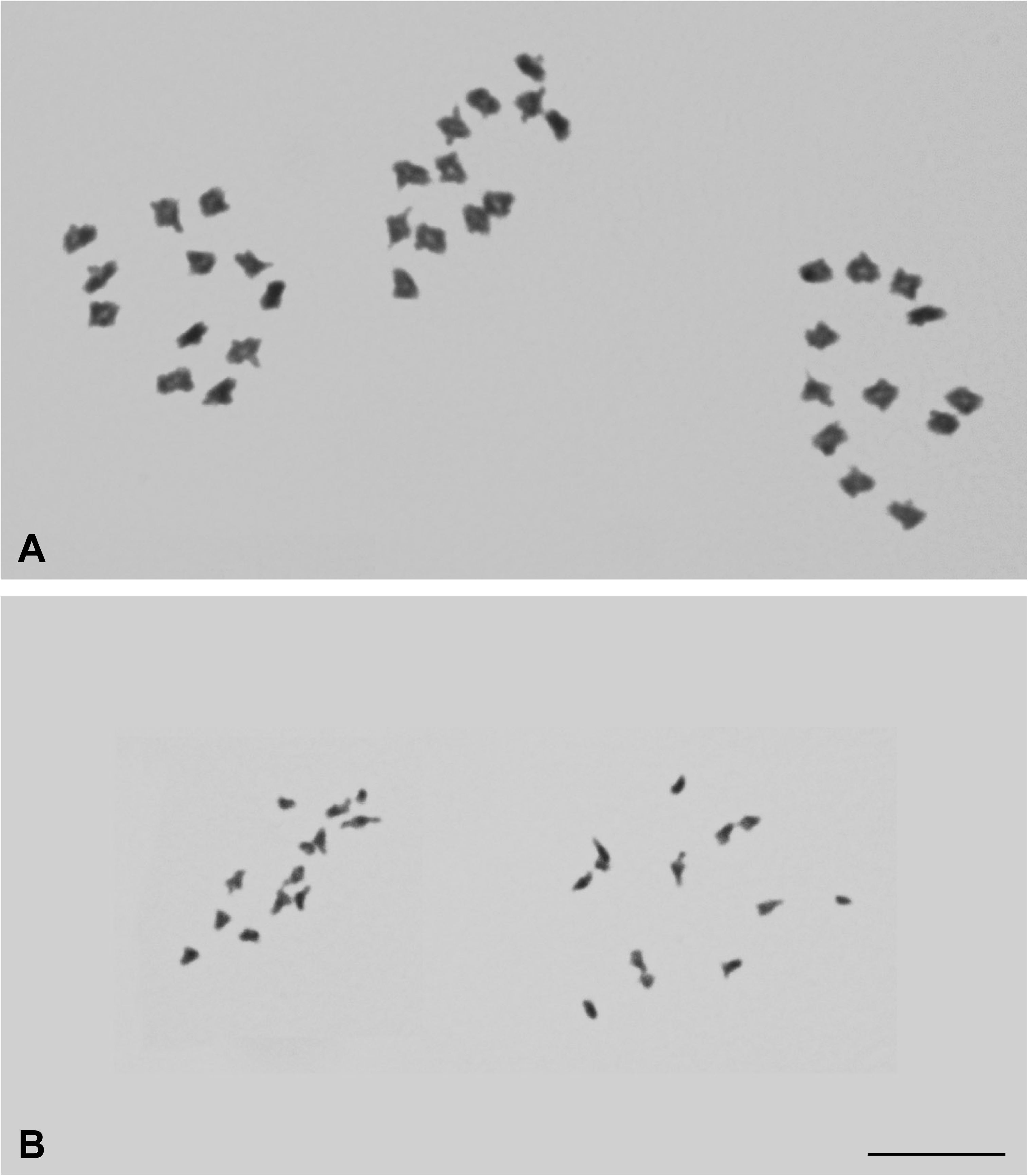
Meiotic first metaphase cells. (A) Three cells of male *Bembidion haplogonum* (B) Two cells of male *Lionepha* “Waterfalls”. Photographs are at the same scale. Scale bar 10µm.

## DISCUSSION

### K-mer Analysis

GenomeScope was unable to converge on nine of the 21 samples in this study, which is likely attributable to the relatively low coverage of those samples. All five specimens of *Pterostichus melanarius* had a coverage of <15X, and all five failed to converge. Of the samples that did converge, the estimates for four of the five *Chlaenius sericeus* were within 6% of the flow cytometric value. The estimates of the three *Lionepha* “Waterfalls” samples were not as close, ranging between 6.8% and 17.5% lower than the true value. Curiously, the estimates for all four *Bembidion haplogonum* were gross underestimates, around half the size of the flow cytometry measurements. The coverage estimated by GenomeScope was double the value expected for a species with a 2,118Mb haploid genome, leading to the underestimated genome size. The k-mer histograms of *Bembidion haplogonum* DNA2544, the specimen with by far the most reads, showed a distinct bimodal profile, with a large peak at 40X coverage and a shorter peak at 80X. This is potentially indicative of a highly heterozygous genome (Vurture *et al*. 2017). Interestingly, GenomeScope also estimated that all four *Bembidion haplogonum* genomes consisted of upwards of 75% repetitive sequence, which is similar to the RepeatExplorer estimate (Table S13). A previous study has found that repeats, as well as high heterozygosity and sequencing errors, can decrease the accuracy of genome size estimation using k-mer frequency (Liu et al. 2013). This suggests the large number of repetitive sequences in the *Bembidion haplogonum* genome may be at least partially responsible for the observed underestimates.

The striking difference between the estimates of the two CovEST models, with “repeat” consistently giving estimates twice the size of “basic,” suggests that neither model is well suited to all genomes. Genome size estimates of the five *Chlaenius sericeus* specimens were closer to the true value using the “basic” model, while the opposite was true of the five *Pterostichus melanarius* specimens. The CovEST “repeat” model was the sequence-based method which came closest to correctly estimating the 2,100Mb genome of *Bembidion haplogonum* with estimates ranging between 1,480Mb to 2,140Mb. This is an interesting result given that *Bembidion haplogonum* was shown by RepeatExplorer and GenomeScope to have the most repeat-rich genome of the eight species analyzed, suggesting that *a priori* knowledge of the amount of repetitive content of a genome may be necessary in order to select an appropriate sequence-based genome size estimation approach.

### Read Mapping

The method we used to infer coverage from read mapping data (averaging the coverage across many single-copy loci) is relatively simple. Despite its simplicity, this approach managed to perform well in some species, particularly those with smaller genomes such as *Chlaenius sericeus* and *Lionepha* “Waterfalls.” Read mapping proved to be inconsistent when estimating genome size in *Pterostichus melanarius*. As with GenomeScope, it consistently underestimated *Bembidion haplogonum* by approximately half.

The reason for the underestimation of *B. haplogonum* genome size based on single-copy read coverage is unclear. Unlike k-mer based methods, which can struggle to assess highly repetitive genomes, the read mapping approach used in this study only infers coverage from single-copy exonic regions. In principle, as long as the selected loci are truly single-copy and the resulting sequence data exhibit no biases in regions sequenced, the estimated coverage should approach the true coverage. It follows that the presence of repetitive sequences elsewhere in the genome should have no effect on this estimate. In actual genomes, evolutionary processes responsible for increasing the size of the genome, such as segmental duplication followed by neofunctionalization or pseudogenization, can lead to the proliferation of genes with similar sequences (Rastogi and Liberles 2005, Levasseur and Pontarotti 2011). Such processes can complicate the selection of a single-copy reference gene set, especially if the group of organisms being studied lacks genomic resources.

We considered the possibility that using a larger set of reference loci from *Bembidion haplogonum* may yield a better coverage estimate. To test this, we repeated read mapping on *Bembidion haplogonum* DNA2544 using the 1421 single-copy orthologs annotated by BUSCO and “No Mito” reads. However, this gave coverage and genome size estimates (94.89X and 1,115.9Mb, respectively) that were very similar to the Regier (96.18X, 1,101Mb) and ODB (93.36X, 1,134Mb) gene sets, which casts doubt on insufficient references as the cause.

### Comparison of Flow Cytometry and Sequence-based Genome Size Estimates

No single sequence-based estimation method proved to be accurate in all cases. Species with large genomes, such as *Bembidion haplogonum* and *Pterostichus melanarius*, appear to present the greatest difficulty for inference of genome size by sequence-based means alone. It is tempting to speculate that the same factors responsible for inaccuracy of k-mer methods are at work in read mapping, especially given that the two methods often yielded underestimates of similar magnitude; however, it is unclear what those factors may be, and they are not necessarily of equal consequence to the two methods. Although both methods use the same underlying reads to estimate coverage, they rely on somewhat different assumptions and portions of the genome. For example, read mapping focuses on coding regions, but k-mer analysis analyzes the entire genomes (Vurture et al. 2017).

The underestimation of the *Bembidion haplogonum* genome by half by most sequence-based methods, including read mapping, is particularly puzzling. A potential explanation that could have accounted for this is recent whole genome duplication or copy number variation; however, cytological examination showed no evidence of genome duplication, nor are such events known in *Bembidion* or near relatives (Maddison 1985, Serrano 1981, 1998). *Bembidion* chromosomes do, however, possess several unusual characteristics. The number and shape of chromosomes is remarkably consistent among species, with almost all species having males with 2n = 22+XY, a value shared by nearly all *Bembidion*, including *B. haplogonum* (Maddison 1985). Each autosome consists primarily of a large heterochromatic central region flanked by small euchromatic tails, and males exhibit achiasmatic meiosis (Maddison 1985, Serrano 1981, 1998). Several species of *Bembidion*, including *Bembidion lividulum*, are also known to possess highly replicated rDNA regions (Sproul and Maddison 2017). It is possible that some of these chromosomal properties are involved in the underestimation of genome size by sequence-based methods, but this is purely speculative, as a mechanism that could account for this unknown. Genomic data can exhibit a variety of sequencing biases (Benjamini and Speed 2012, Desvillechabrol et al. 2016), which could cause certain portions of the genome to be underrepresented relative to the rest, leading to misestimation of genome size. For example, consider an organism with a genome largely composed of AT-rich repetitive sequences. Because a highly AT- or GC-rich sequence is less likely to be amplified during library preparation (Benjamini and Speed 2012, Chen et al. 2013, Ross et al. 2013), the resulting library would contain fewer reads derived from these repeats and more from the more AT-balanced protein-coding regions, violating a core assumption of the sequence-based genome size estimation methods (i.e. all bases have an equal probability of being sequenced). The resulting coverage estimates would be artificially increased, causing a decrease in the subsequent genome size estimates.

This particular issue, however, does not seem to be the case with *Bembidion haplogonum*, as the GC content of the reads identified as repetitive elements by RepeatExplorer (33.0%) is nearly identical to the overall GC content of the library (32.2%). It was also unremarkable compared to the GC content of the other seven species analyzed, which was 32.3% on average and ranged from 29.0% to 36.5%.

A more interesting possibility is that some biological process is physically causing parts of the genome to be disproportionately represented within nuclei. Flow cytometric estimates compare the unreplicated DNA amount in the G1 peak of the sample with the G1 peak of the standard. The G1 peak is typically the major peak. If the major peak is G2 rather than G1 the estimate will be a 2C value or twice the value assigned as a 1C. An increase in DNA per nucleus is also created by underreplication, a process by which portions of the genome are replicated more slowly during cell growth and division, resulting in fewer copies of certain parts of the chromosomes (Belyaeva et al. 2008, Spradling and Orr-Weaver 1987). The most well-known example of this occurs in the polytene salivary glands of *Drosophila* (Belyakin 2005, Yarosh and Spradling 2014). Using the NGS genome data of Yarosh and Spradling (2014), we repeated several of the sequence-based analyses employed in this study (Table S14), and observed that genomic sequence from polytene salivary gland cells produced genome size estimates significantly (two-sample t-test, p=0.000028) lower than those derived from unreplicated embryonic cells, by an average of 36%. While cytological study of *Bembidion haplogonum* effectively rules out any polytene-like chromosomal structures in the tissues sequenced for this study, it is possible some other process, perhaps related to the unusual characteristics of *Bembidion* chromosomes, is causing significant underreplication in the genome.

## CONCLUSIONS

Increasing our knowledge of genome sizes across the tree of life is important for a deeper understanding of genomic evolution, but the pace of this increase is currently very slow. In this paper we have presented the first published flow cytometry estimates for five species of carabid beetles; although this nearly doubles the number of carabids with genome size estimates, we now have flow-cytometric estimates for less than 0.03% of carabid species.

With the explosion of short-read sequence data, and the development of methods to infer genome sizes from these, the pace may increase, if the sequence-based methods are accurate enough. We found that some of the sequence-based estimation techniques we investigated were consistent with flow cytometry in some species, but no single technique was uniformly congruent with flow cytometry. Flow cytometry should be the preferred option for estimating genome size when live material and adequate resources are available. In cases where this is not possible, especially when working with rare or extinct organisms, sequence-based methods can provide an initial estimate of the size of a genome, and they may be well suited for cases when the genome is likely to be small and non-repetitive. However, our work shows that these techniques can also be misleading, particularly in large or highly repetitive genomes. Further study on more genomes with different properties is necessary to better understand the strengths and weaknesses of these methods.

## ACKNOWLEDGEMENTS

We wish to thank Kipling W. Will for providing specimens of *Chlaenius sericeus*, and John S. Sproul for providing specimens and Illumina genomic data for *Bembidion lividulum*. We are also grateful to Mark Dasenko at the OSU CGRB for performing all the Illumina sequencing for this project, as well as constructing several Illumina libraries. This project was partially supported by Harold E. and Leona M. Rice Endowment Fund at Oregon State University and NSF Grant DEB-1702062 to JMP and DRM.

