## Supplementary figures and images for "Measuring genome sizes using read-depth, k-mers, and flow cytometry: methodological comparisons in beetles (Coleoptera)"

### plot.log.png

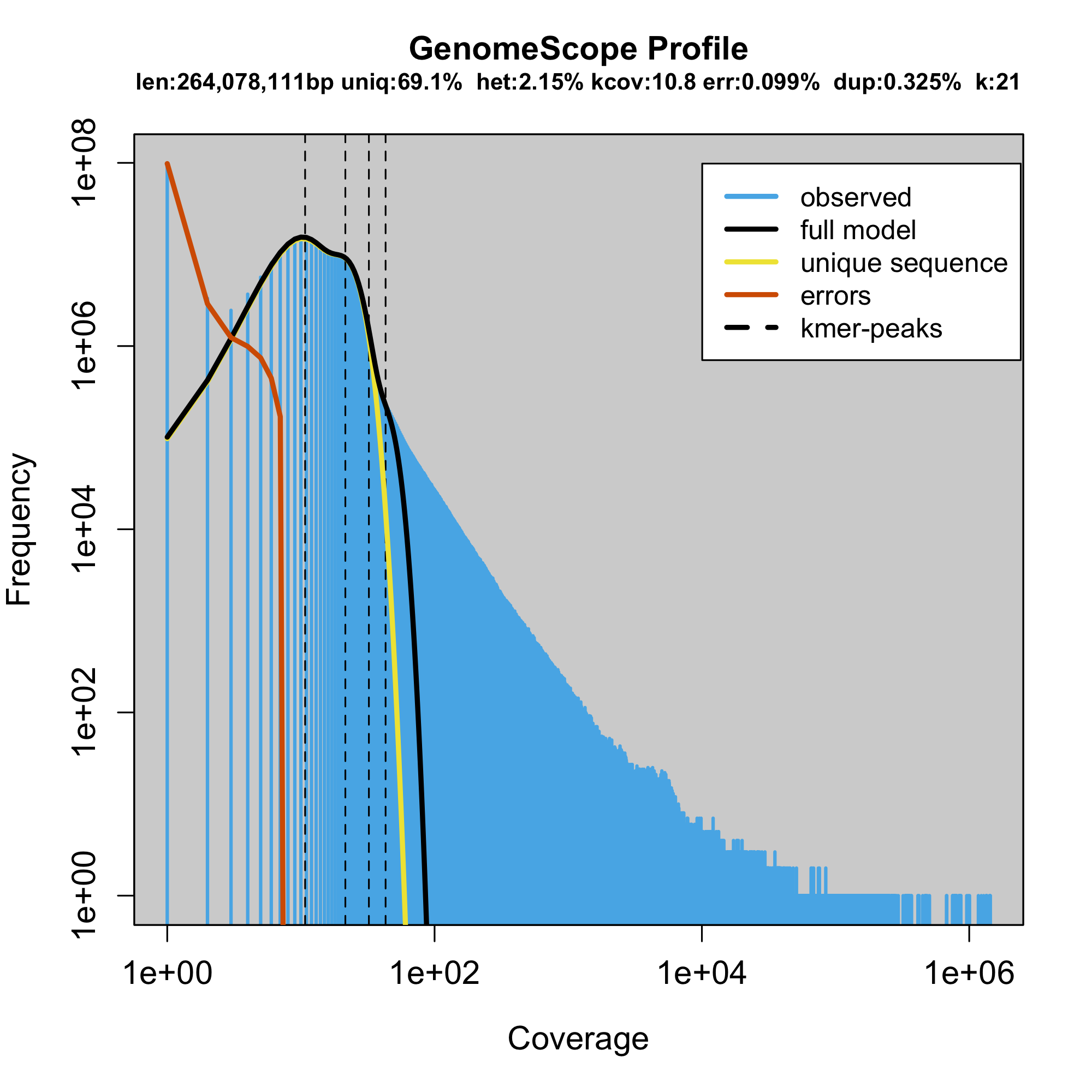

### plot.log.png

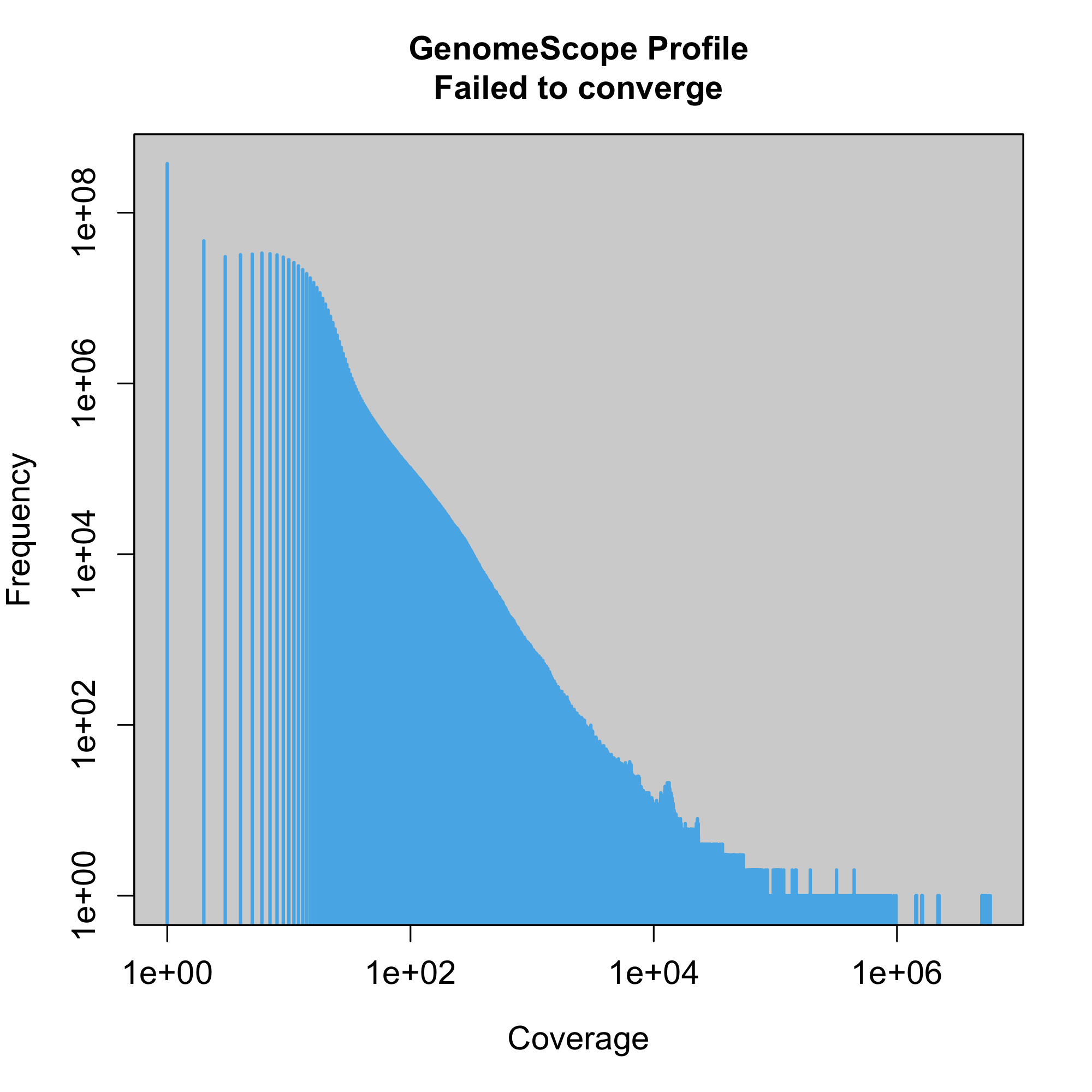

### plot.log.png

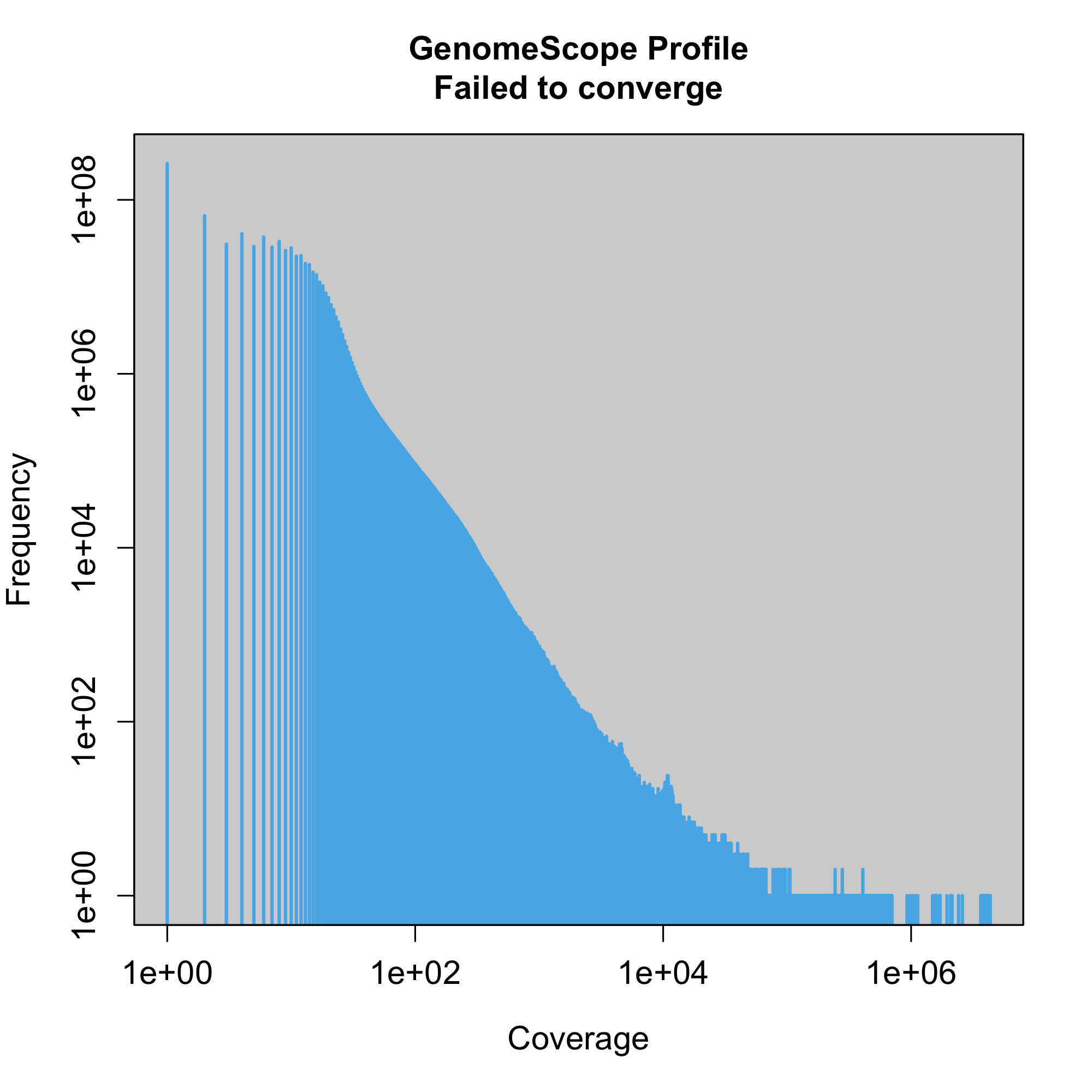

### plot.log.png

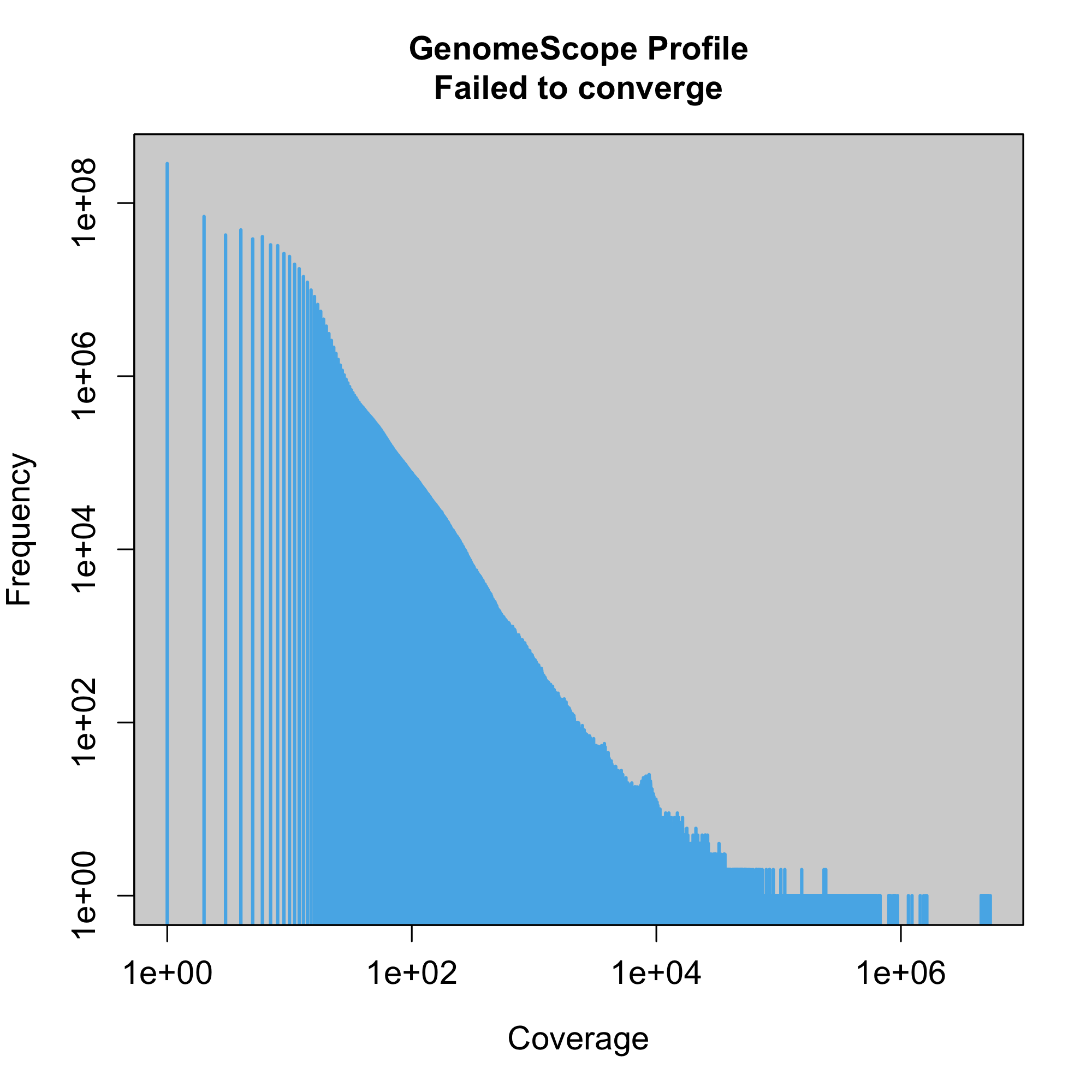

### plot.log.png

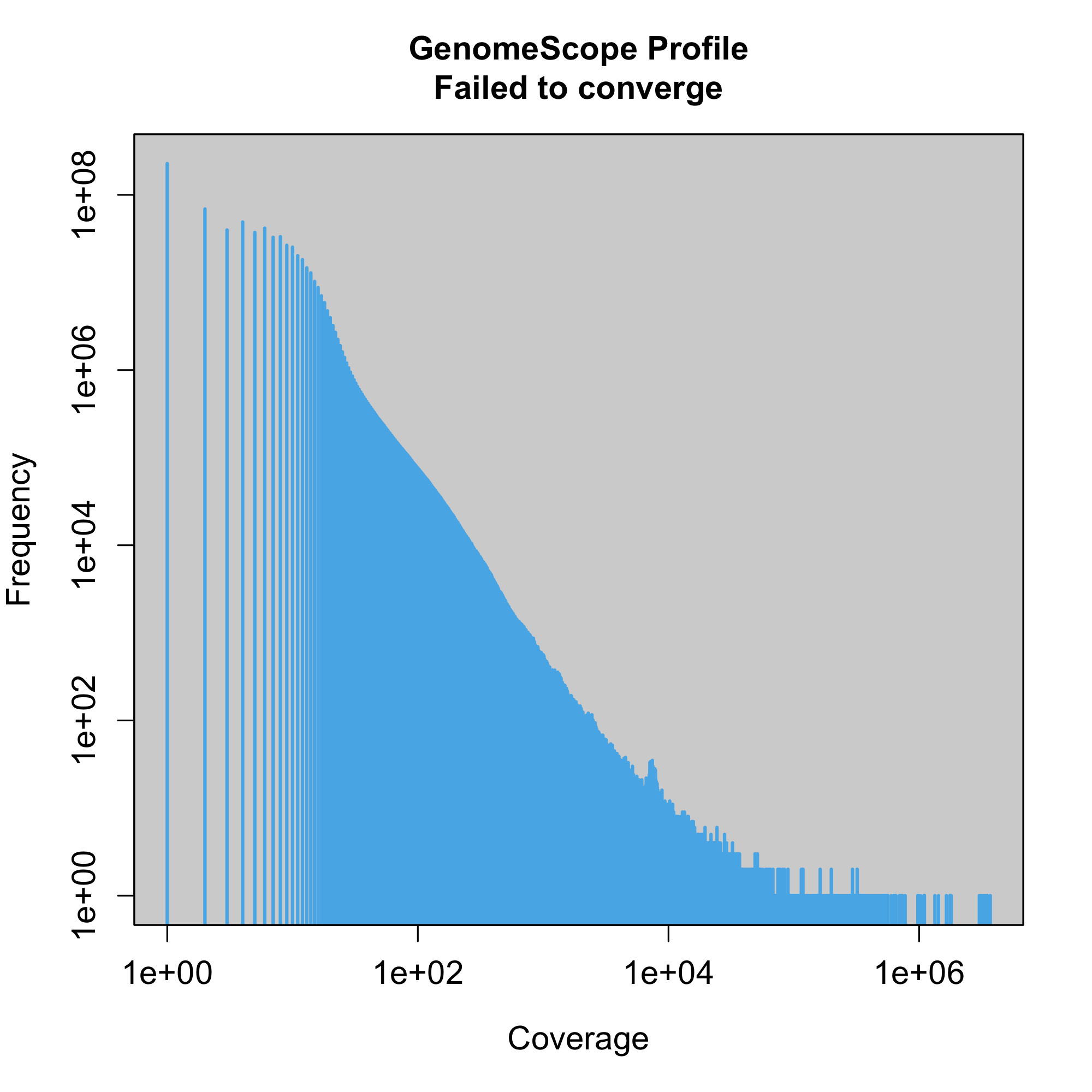

### plot.log.png

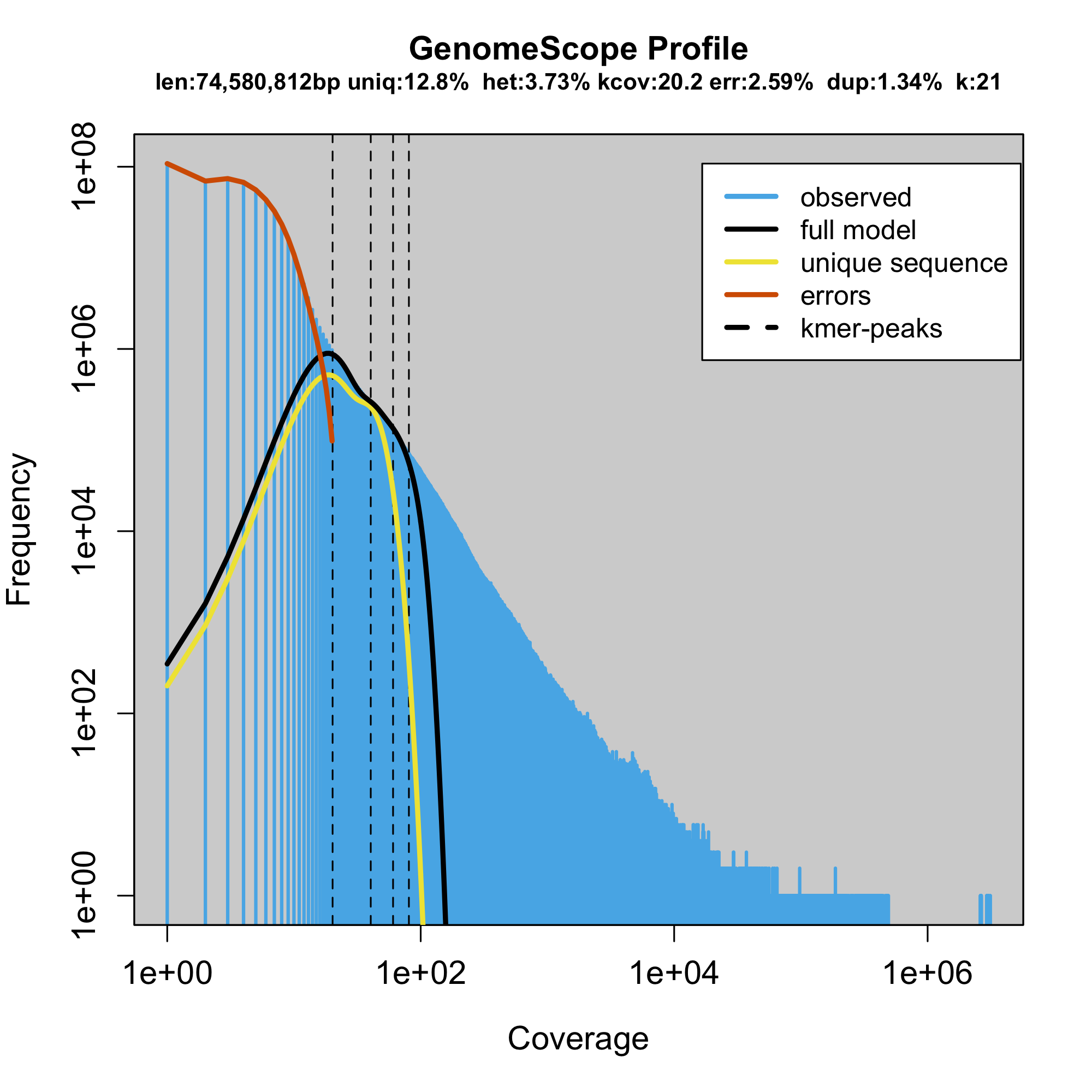

### plot.log.png

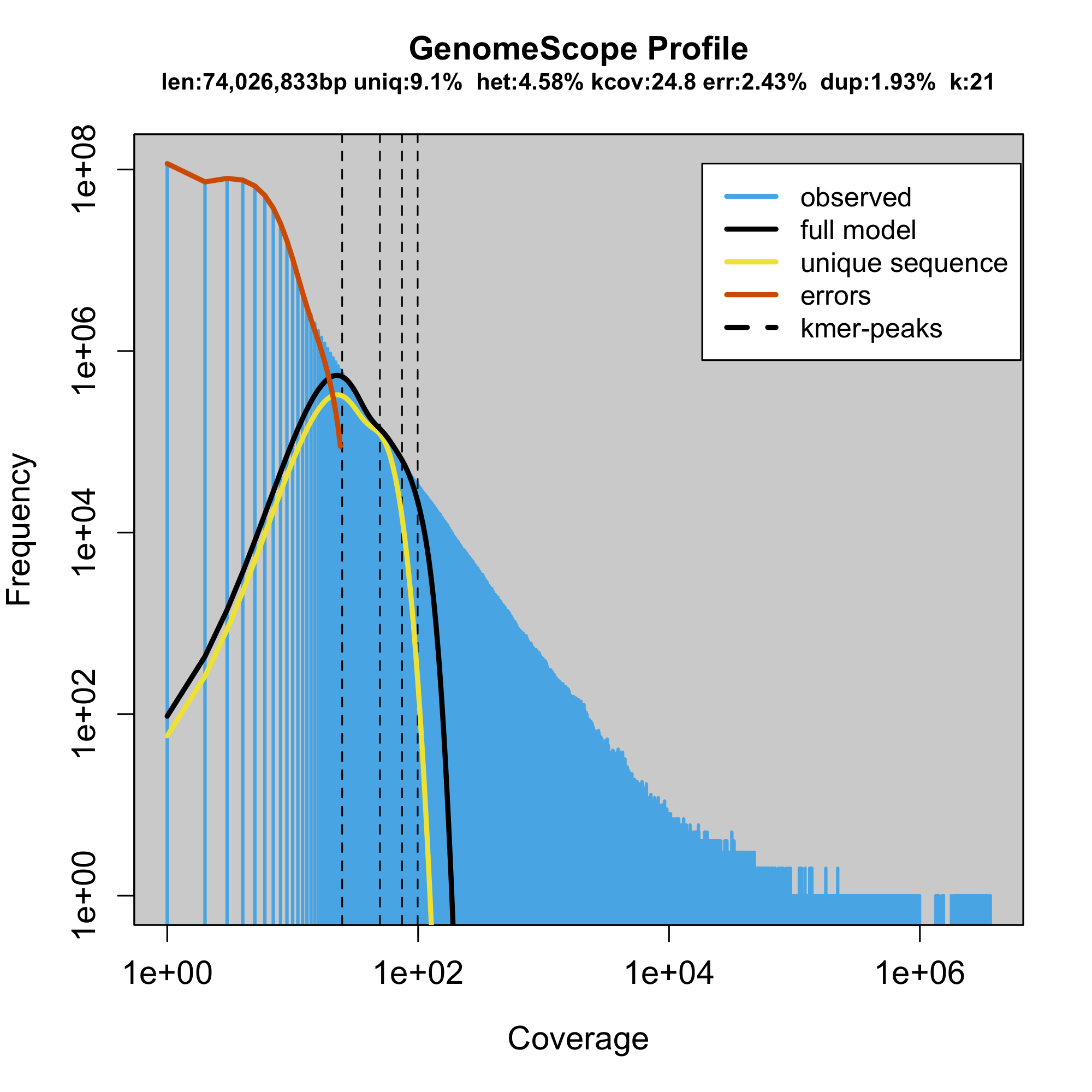

### plot.log.png

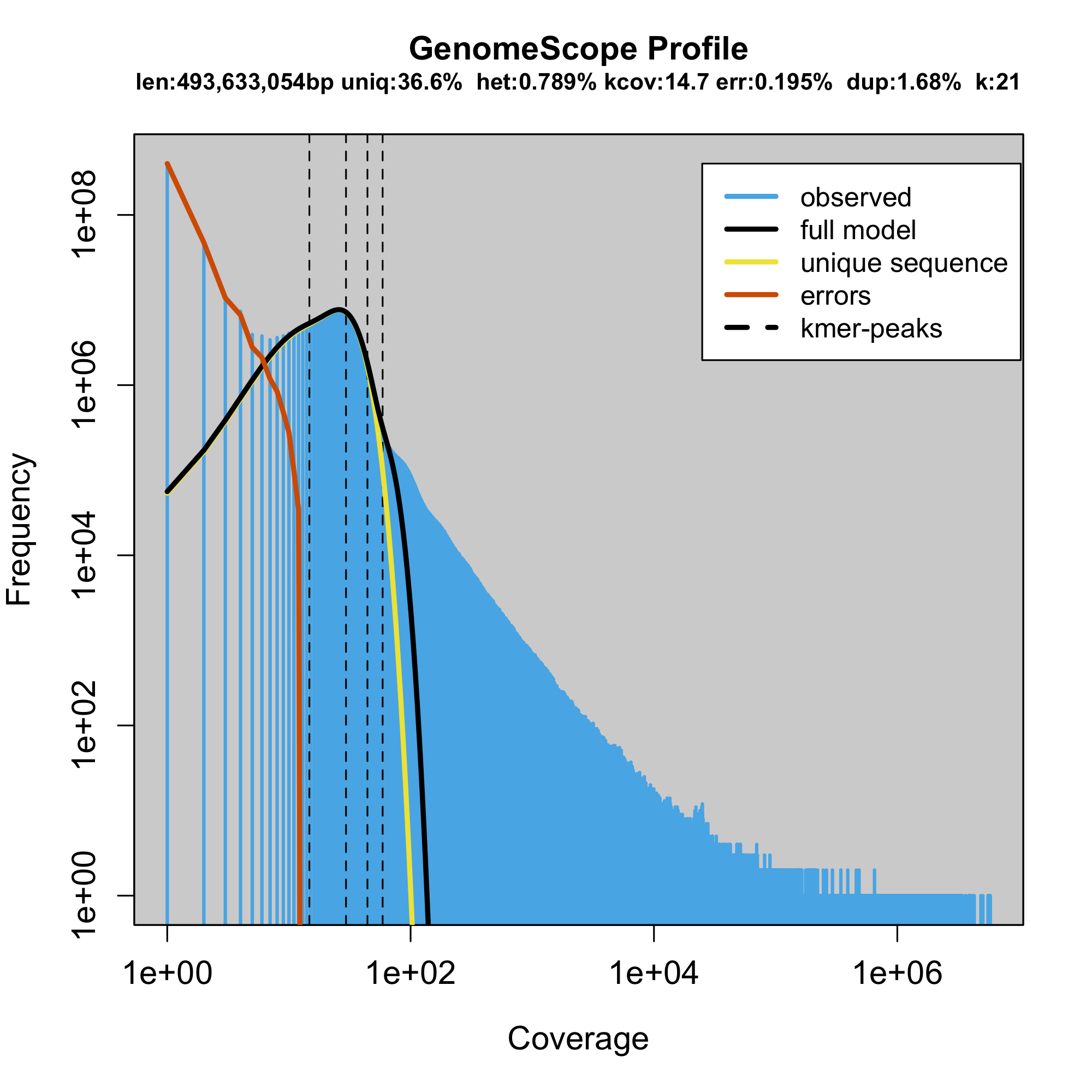

### plot.log.png

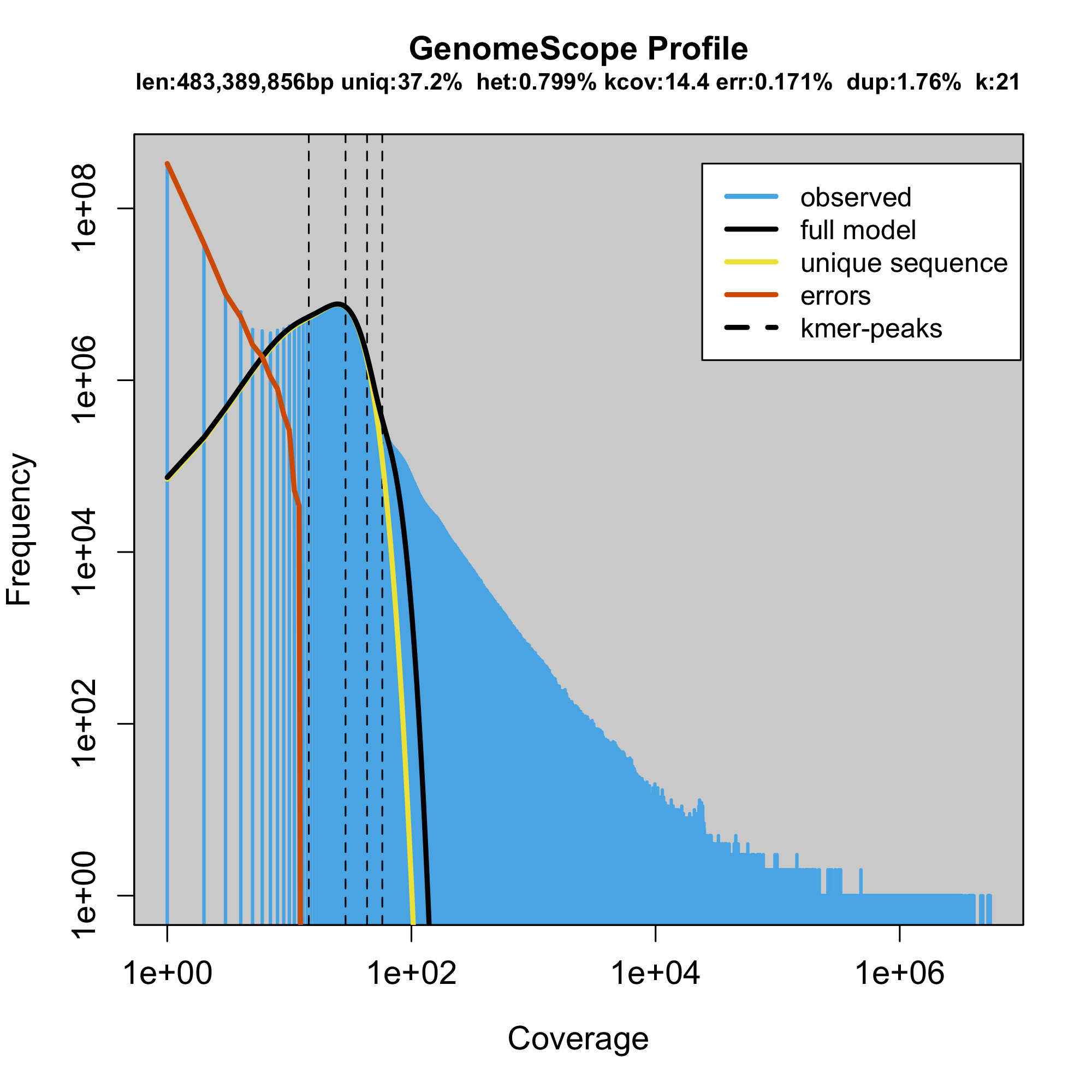

### plot.log.png

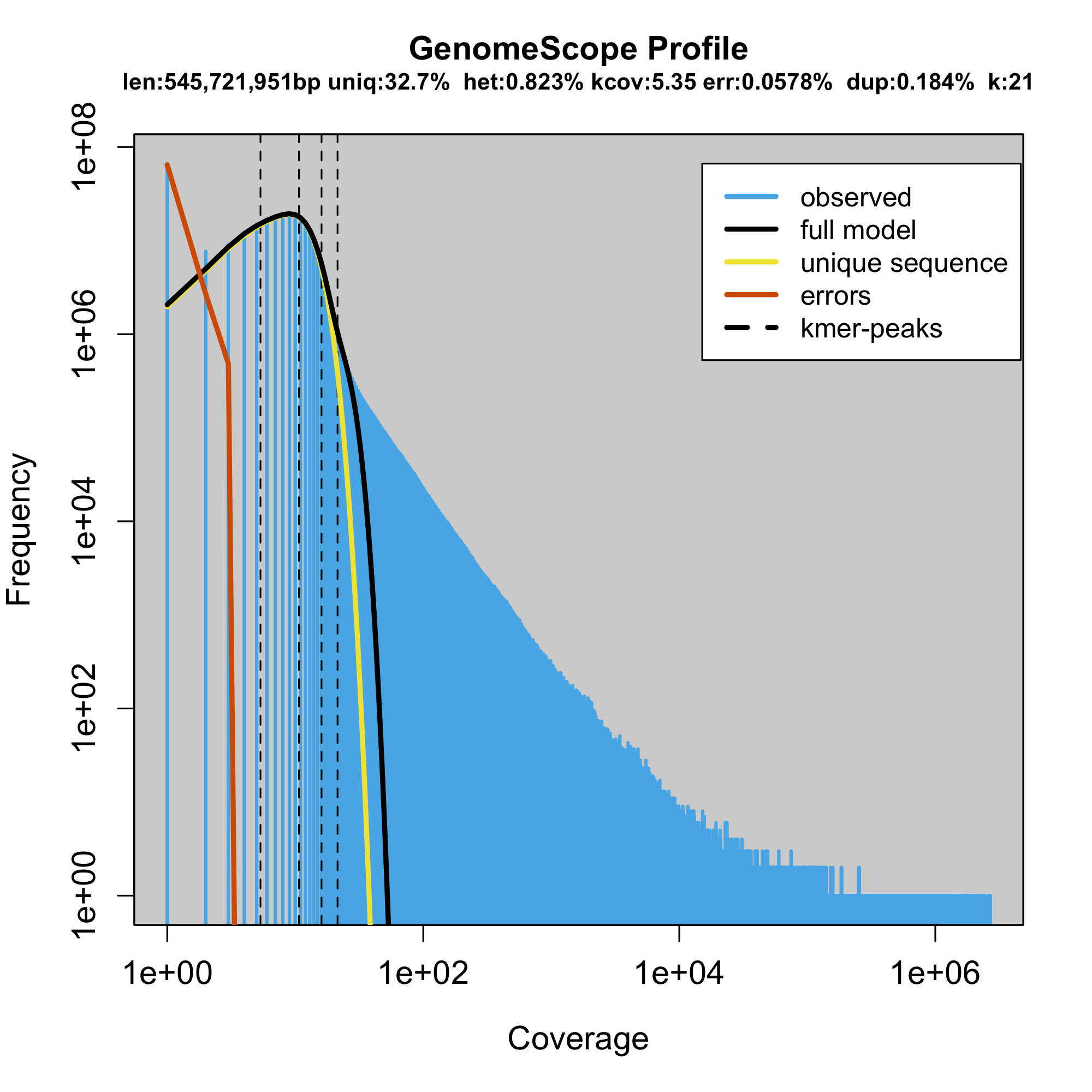

### plot.log.png

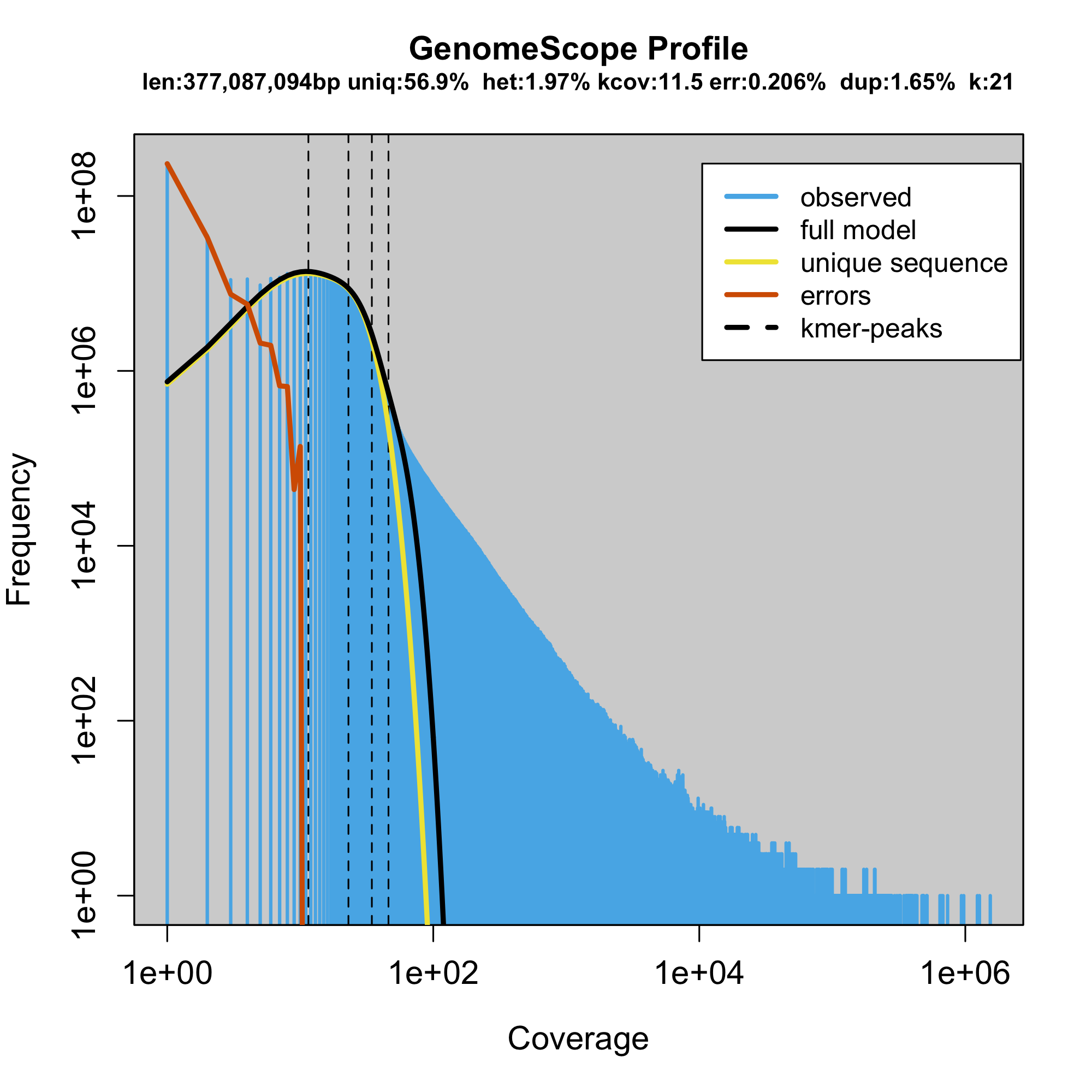

### plot.log.png

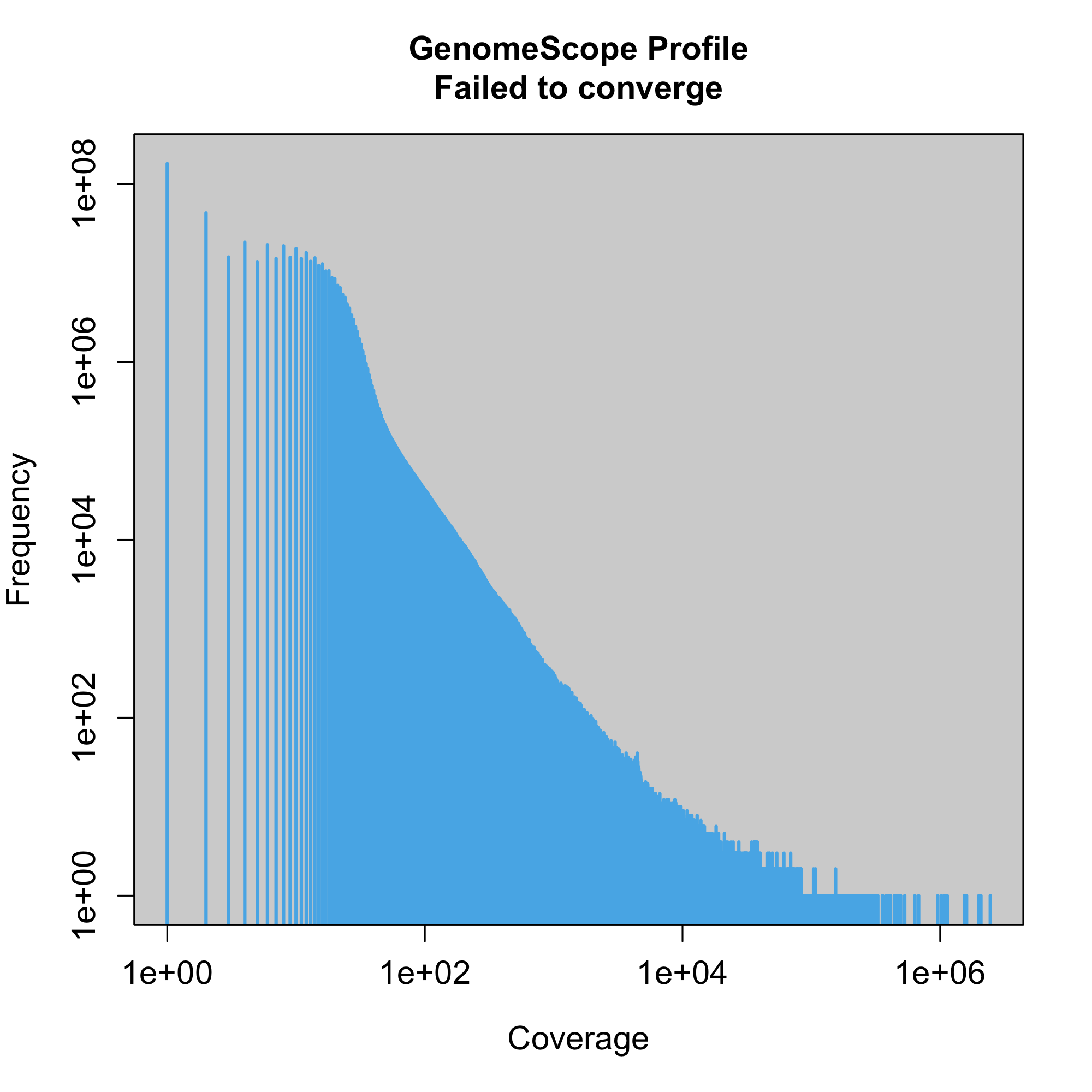

### plot.log.png

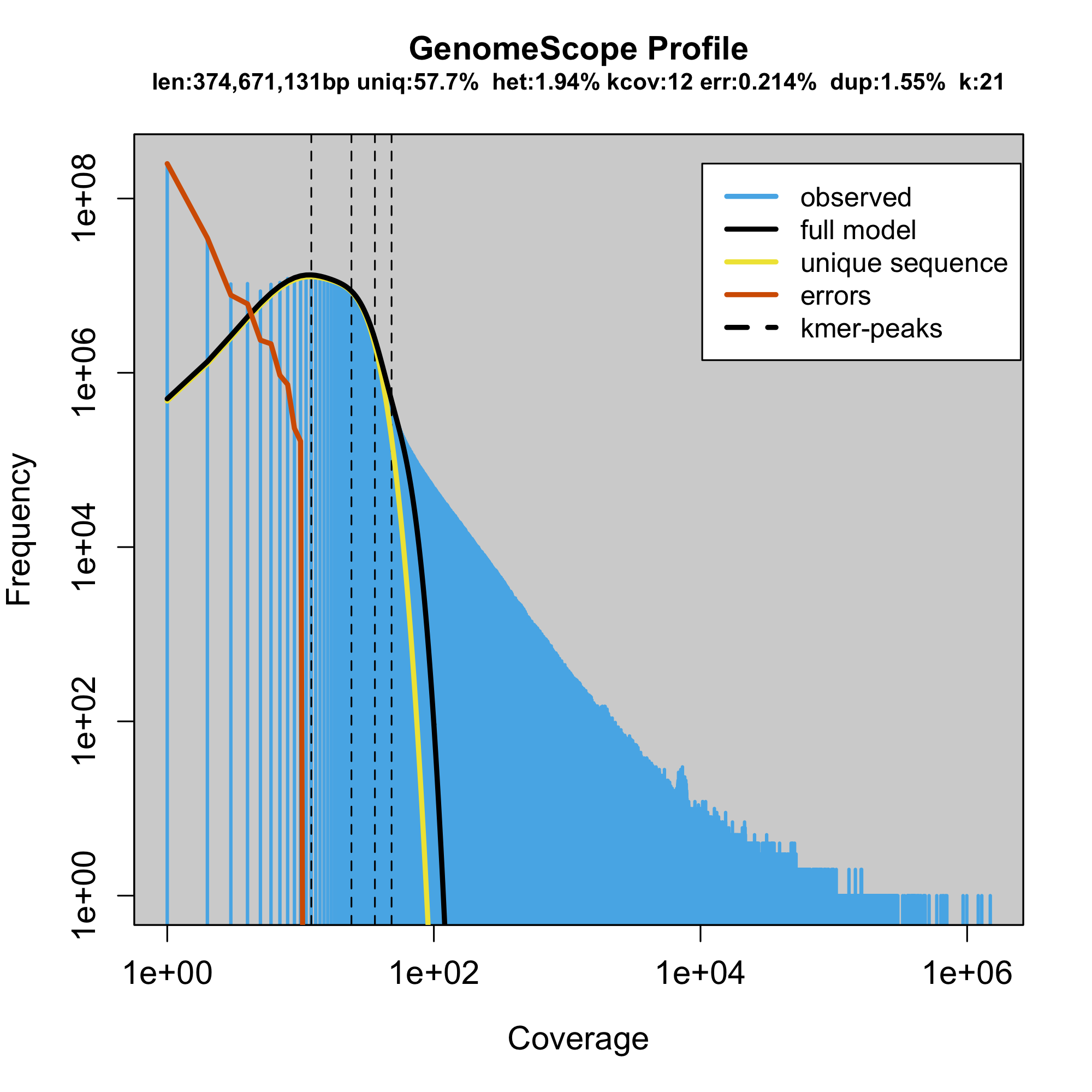

### plot.log.png

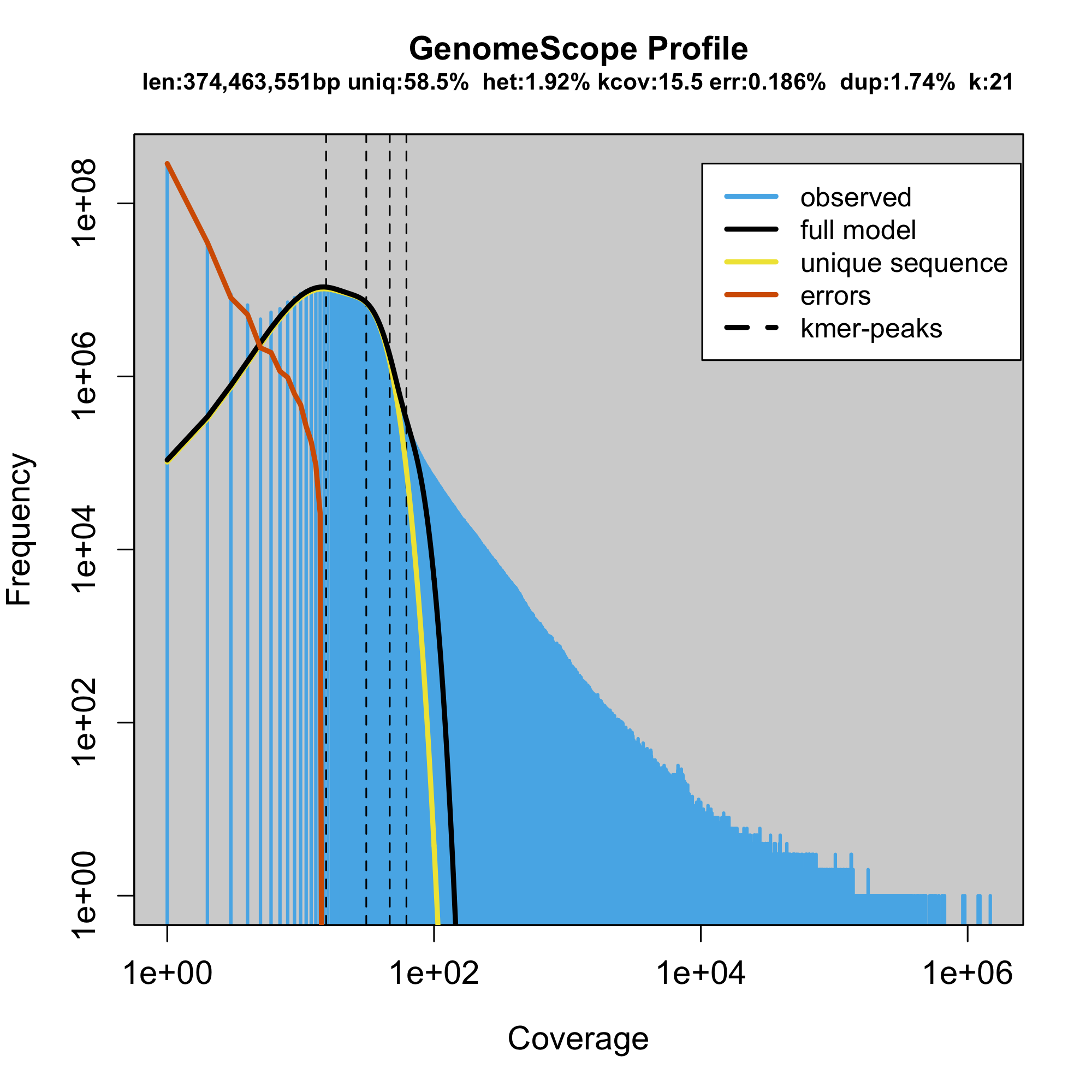

### plot.log.png

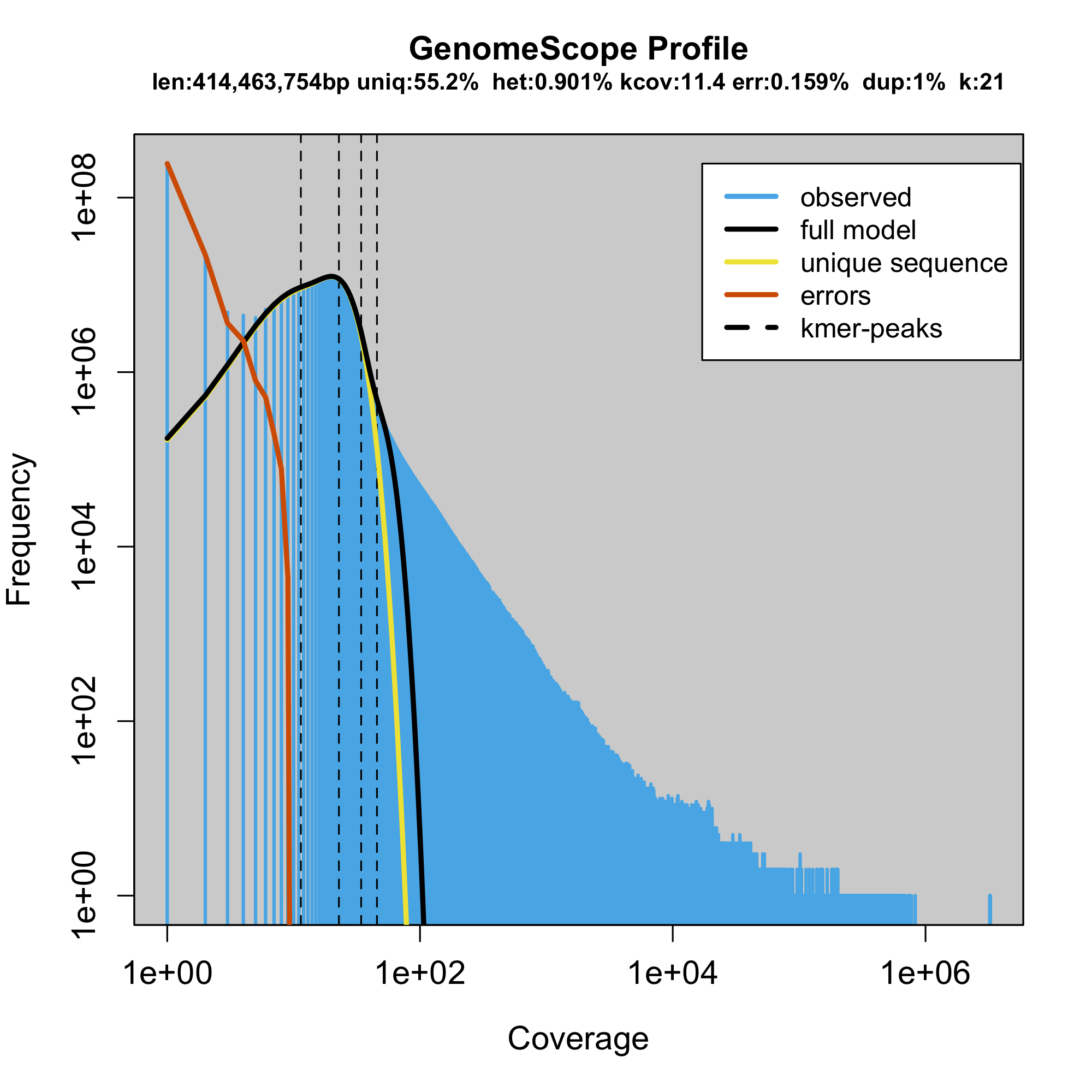

### plot.log.png

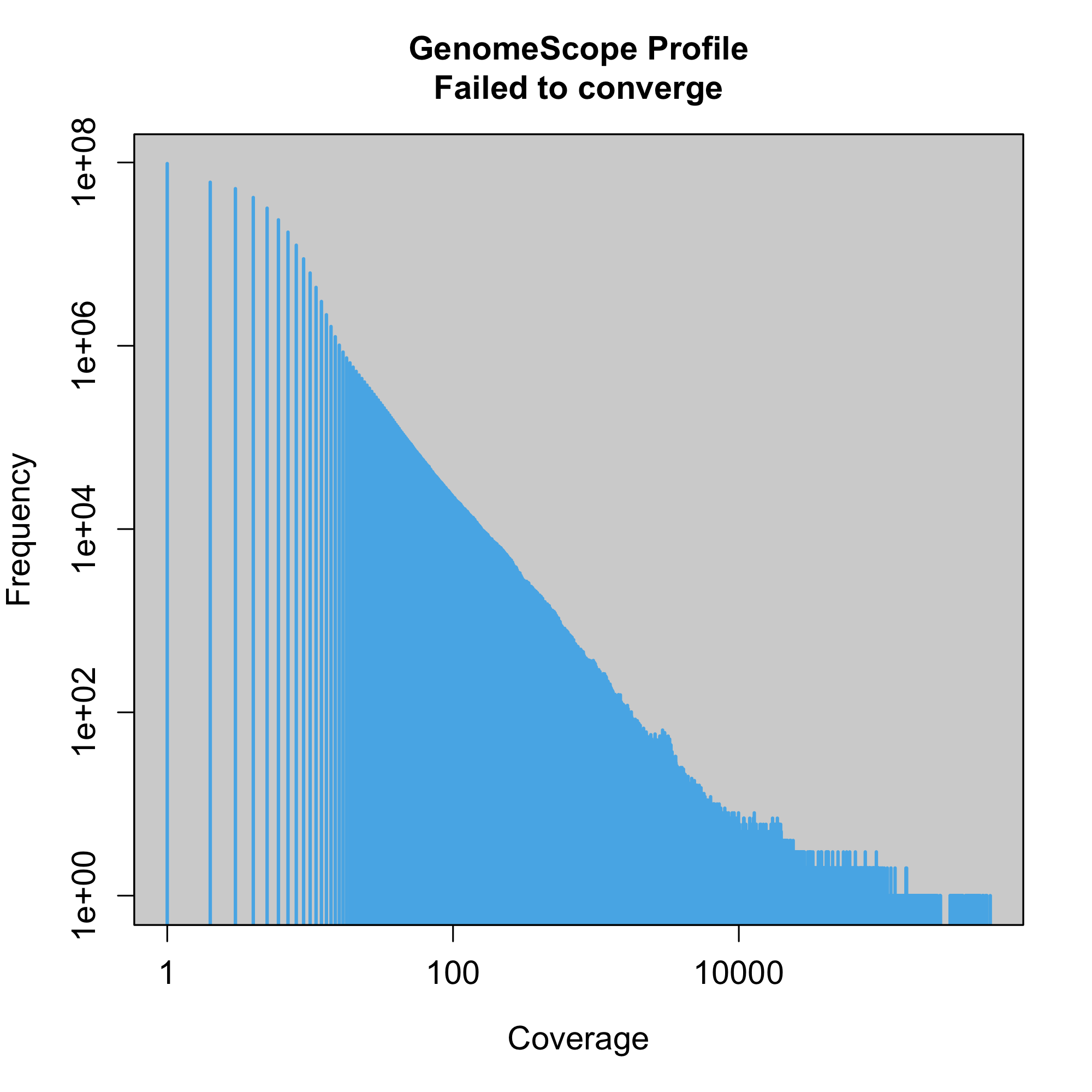

### plot.log.png

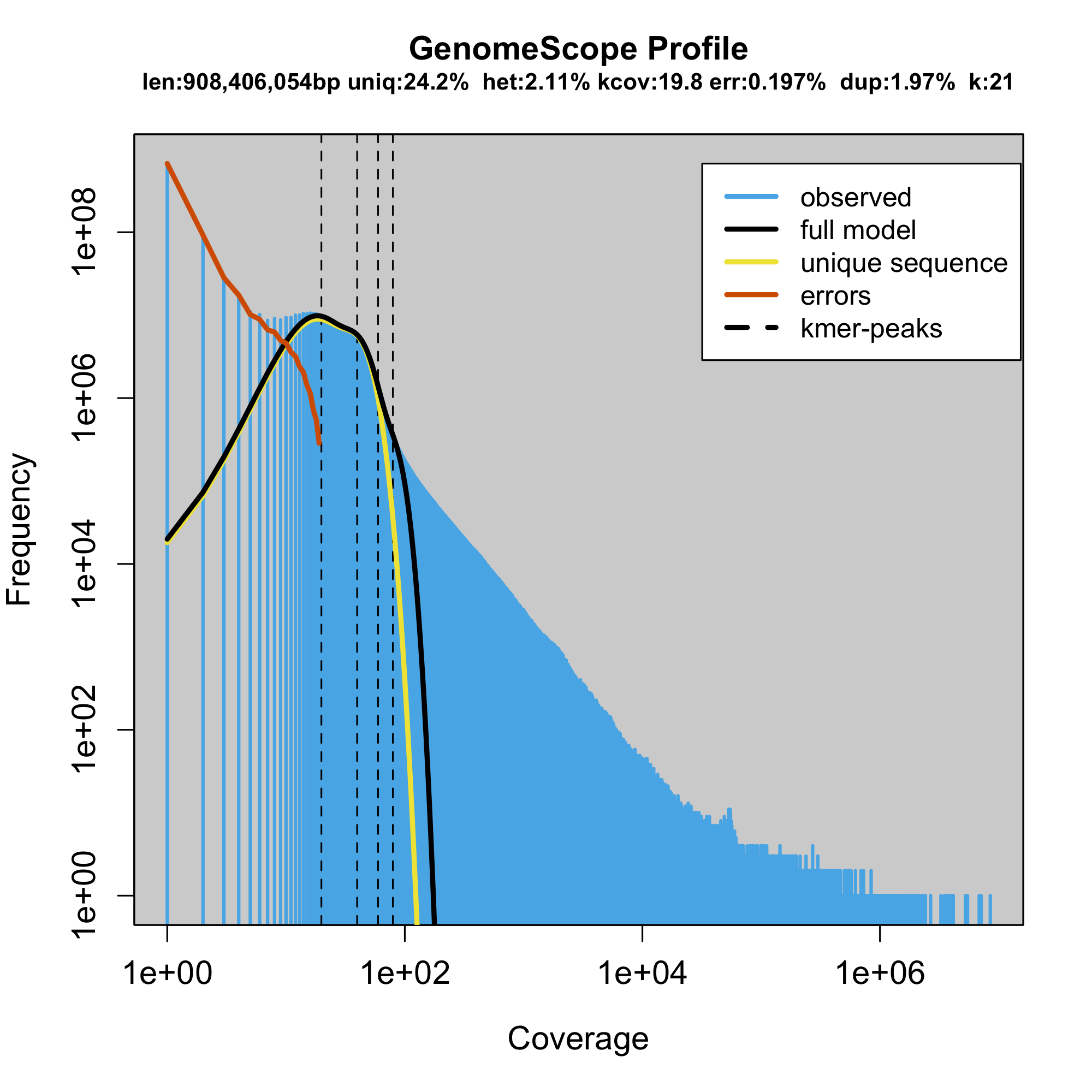

### plot.log.png

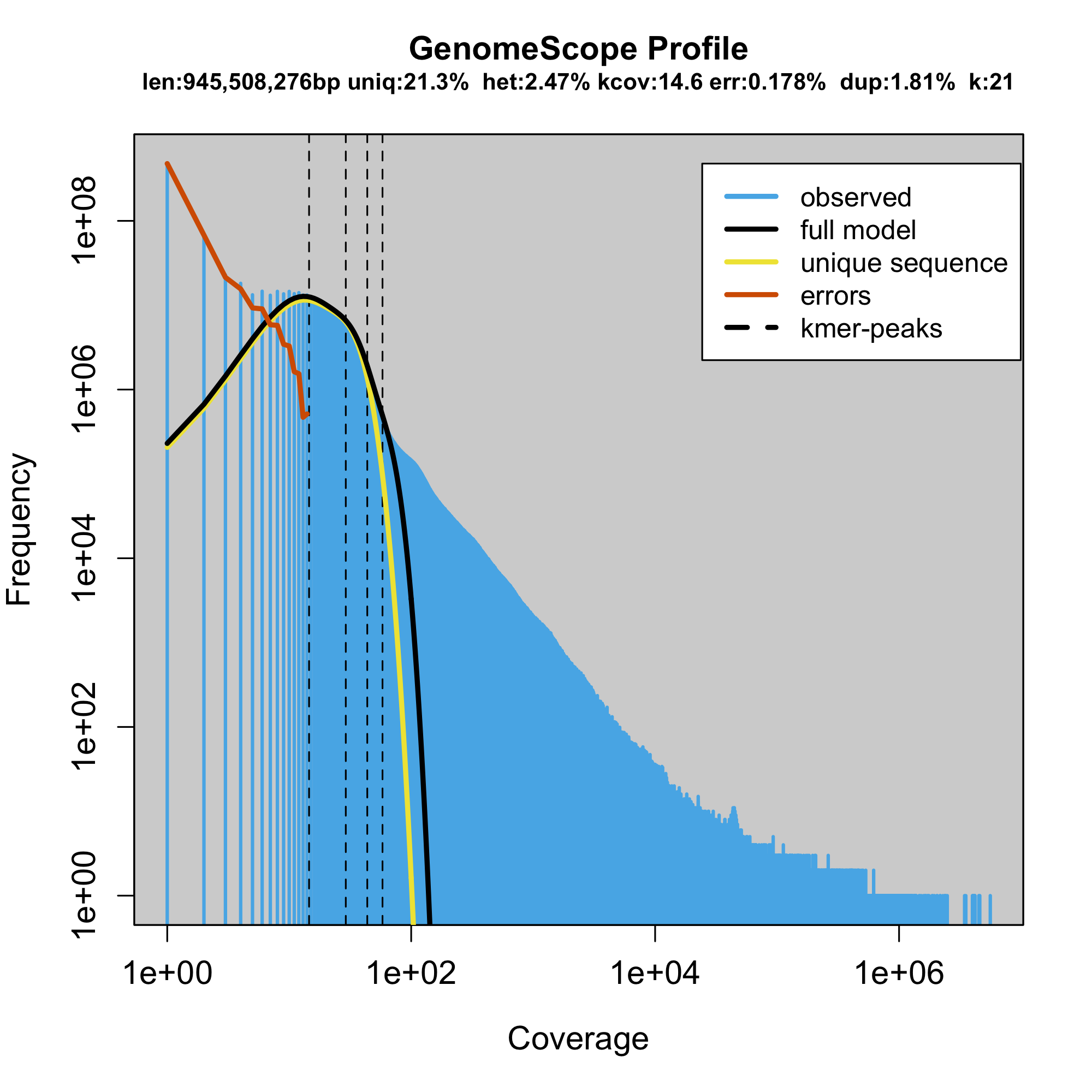

### plot.log.png

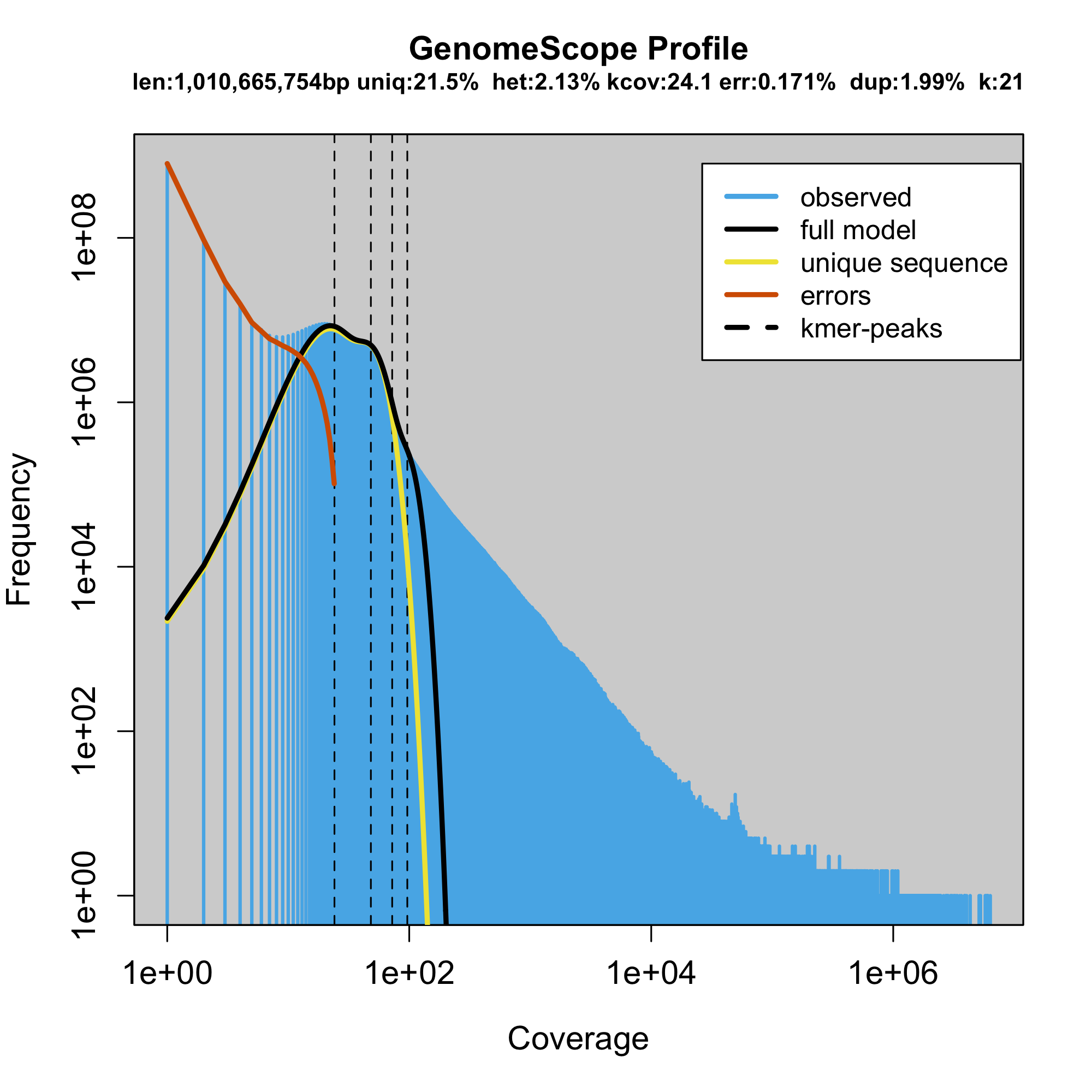

### plot.log.png

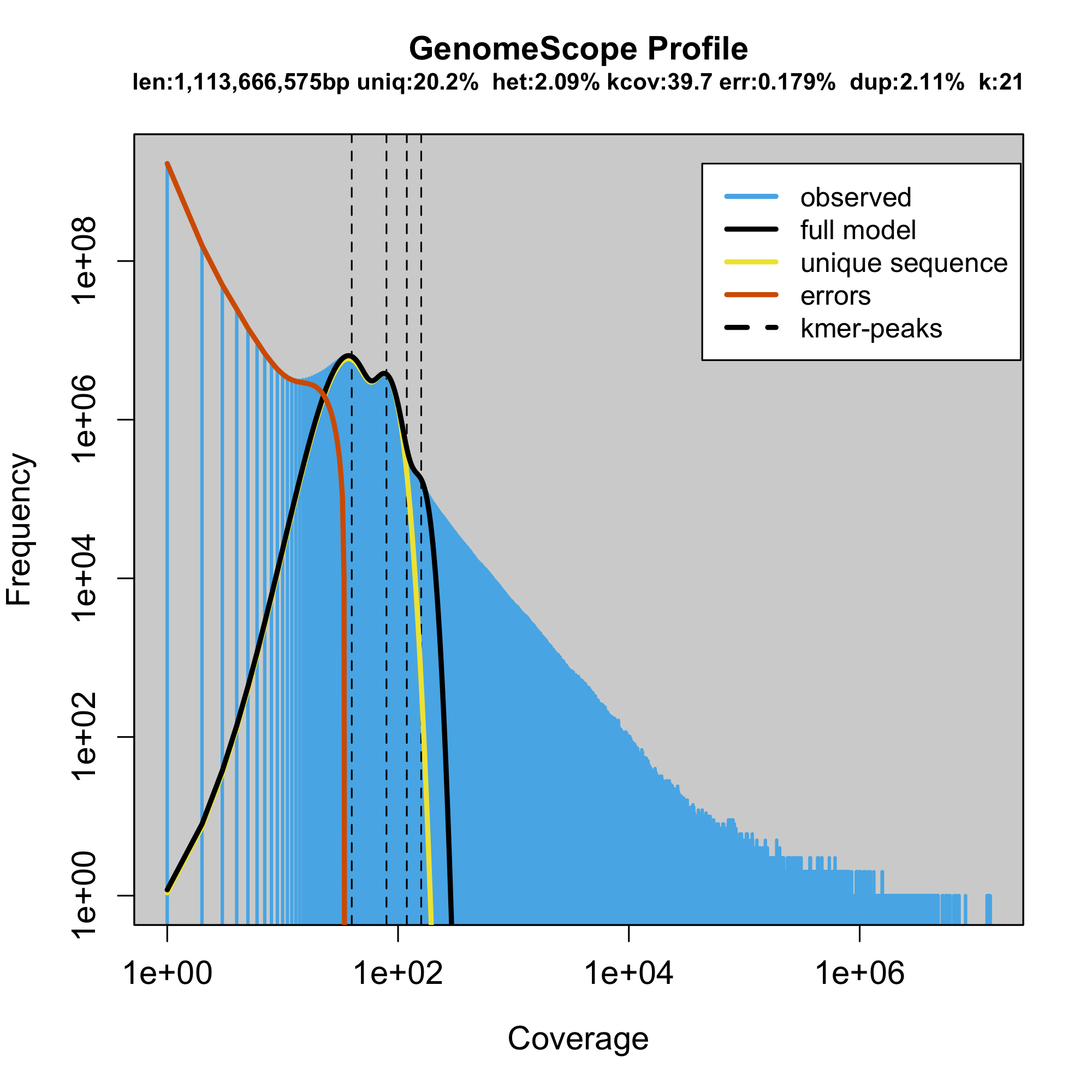

### plot.log.png

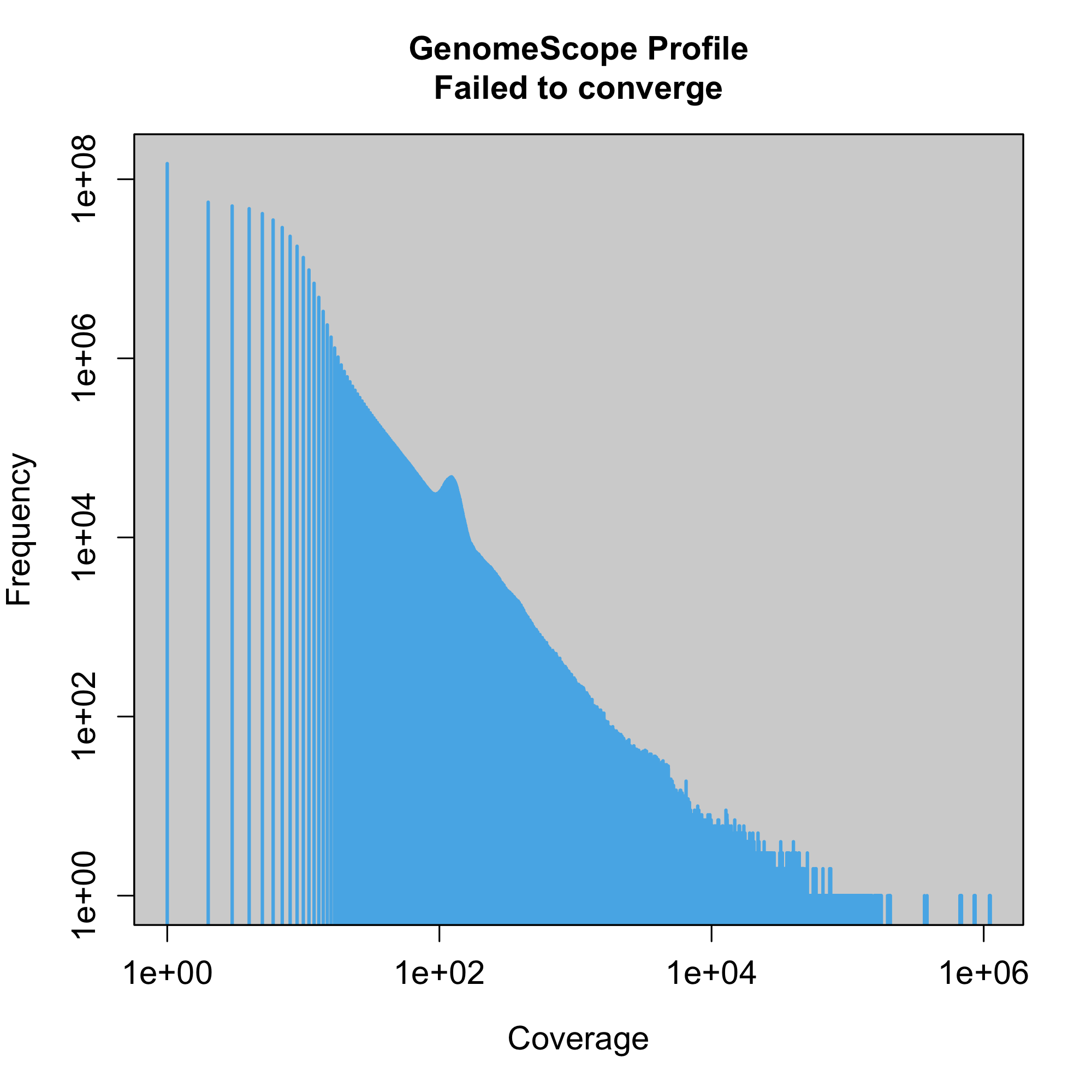

### plot.png

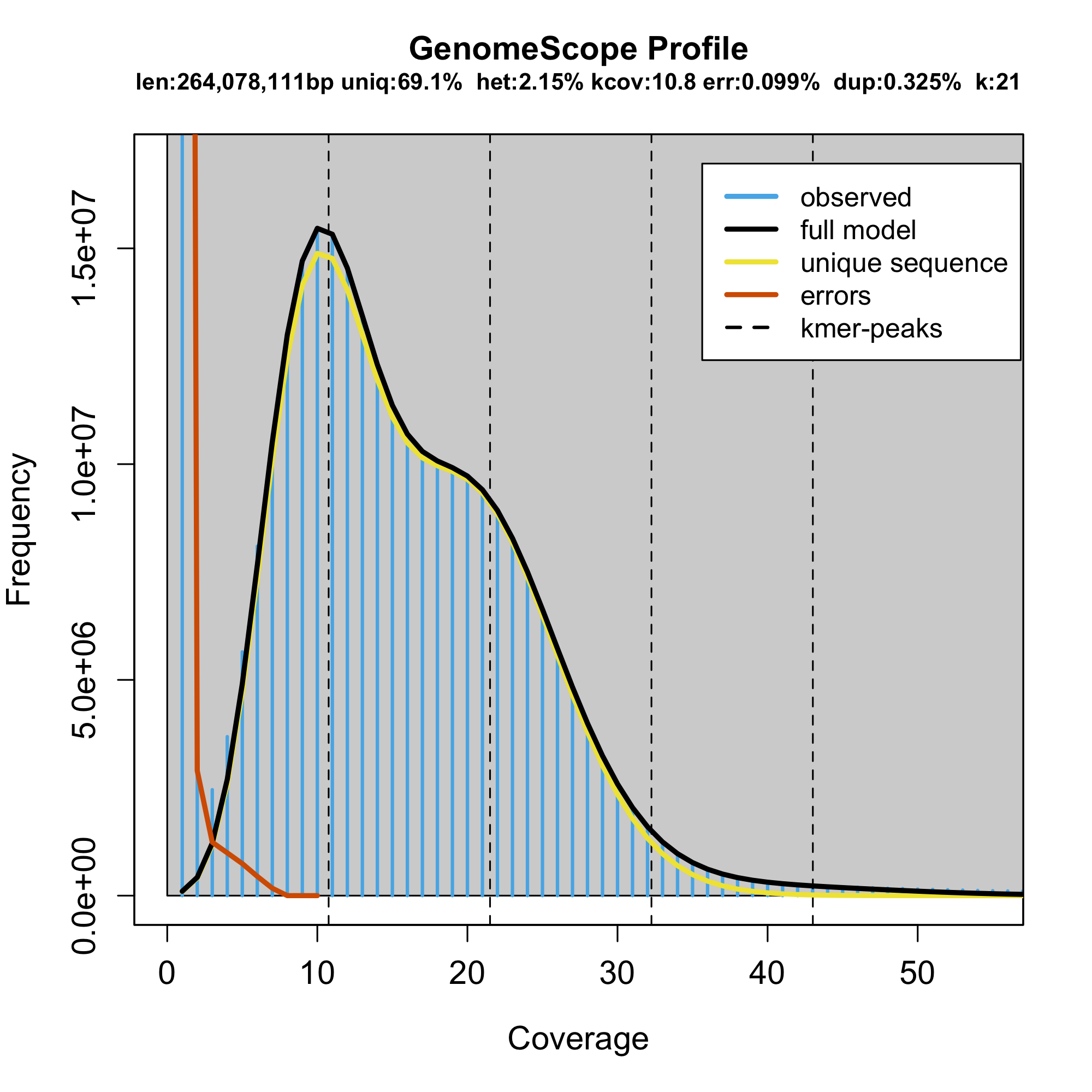

### plot.png

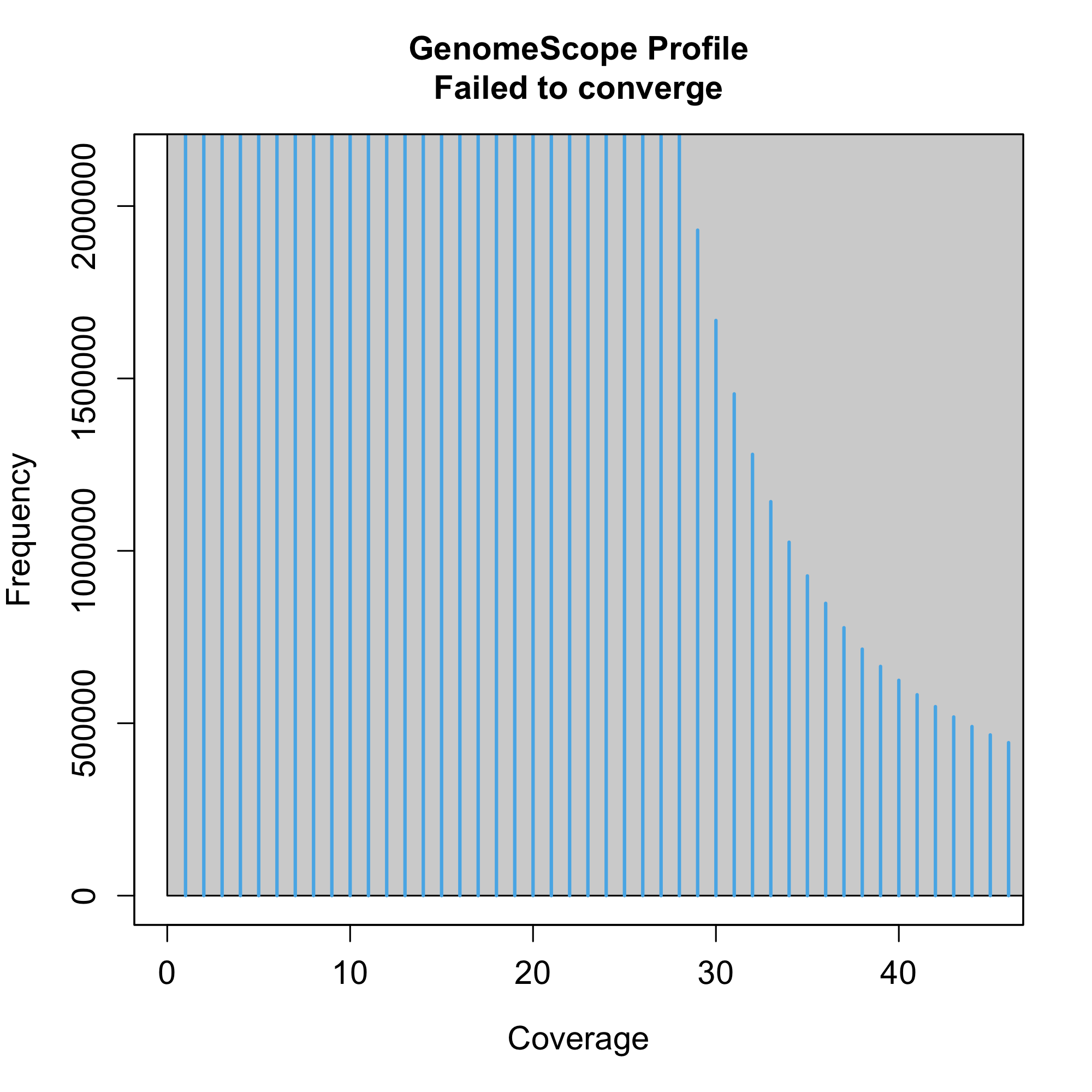

### plot.png

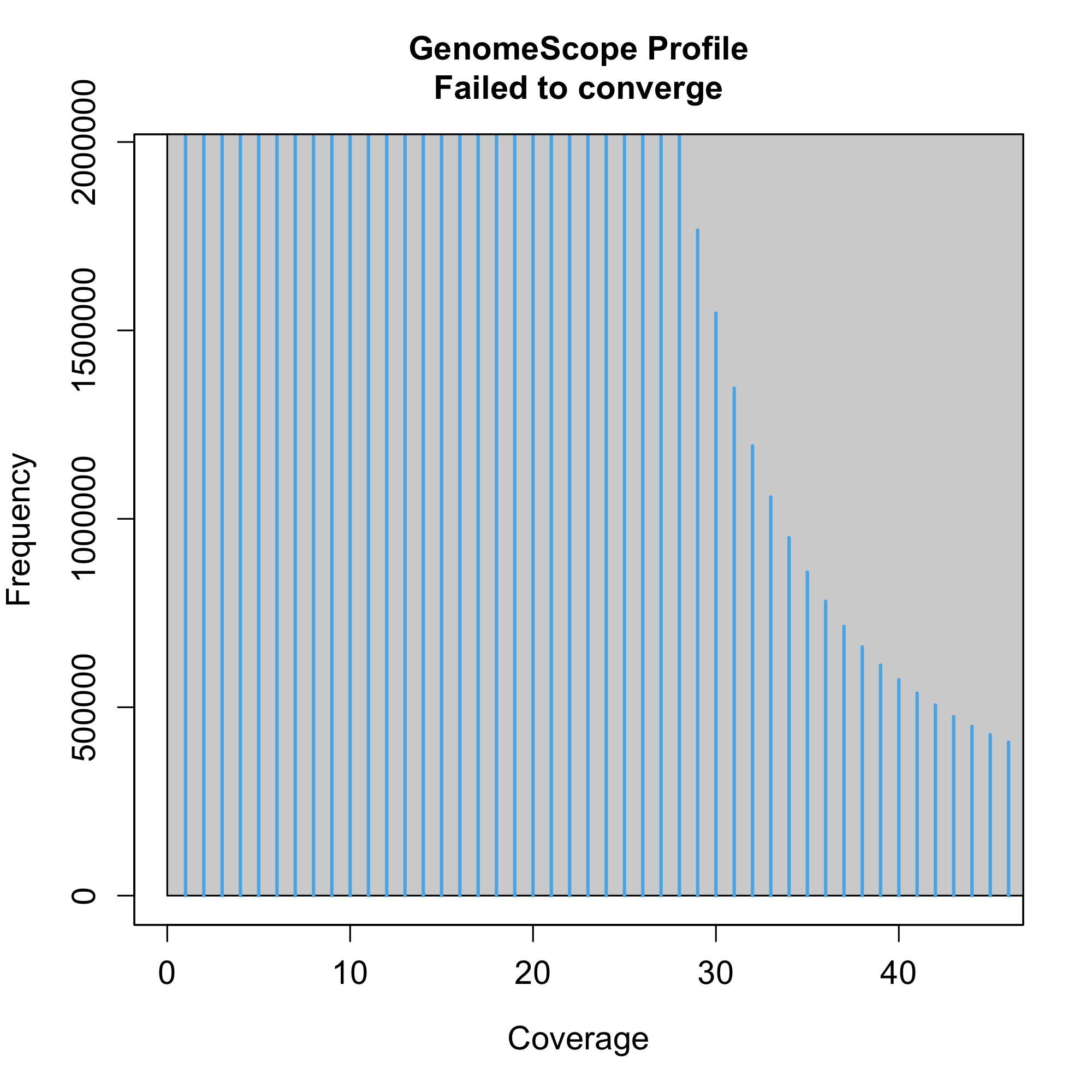

### plot.png

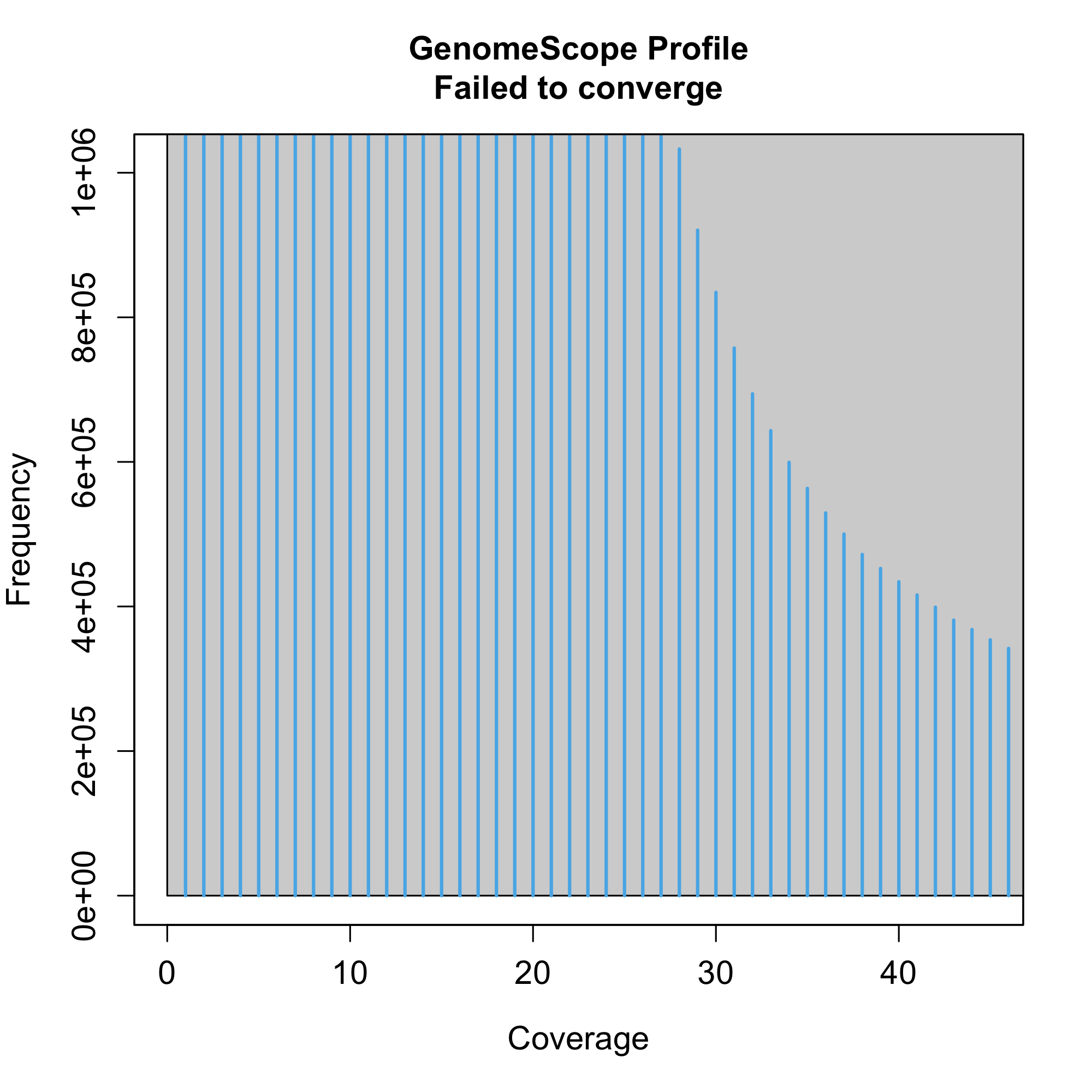

### plot.png

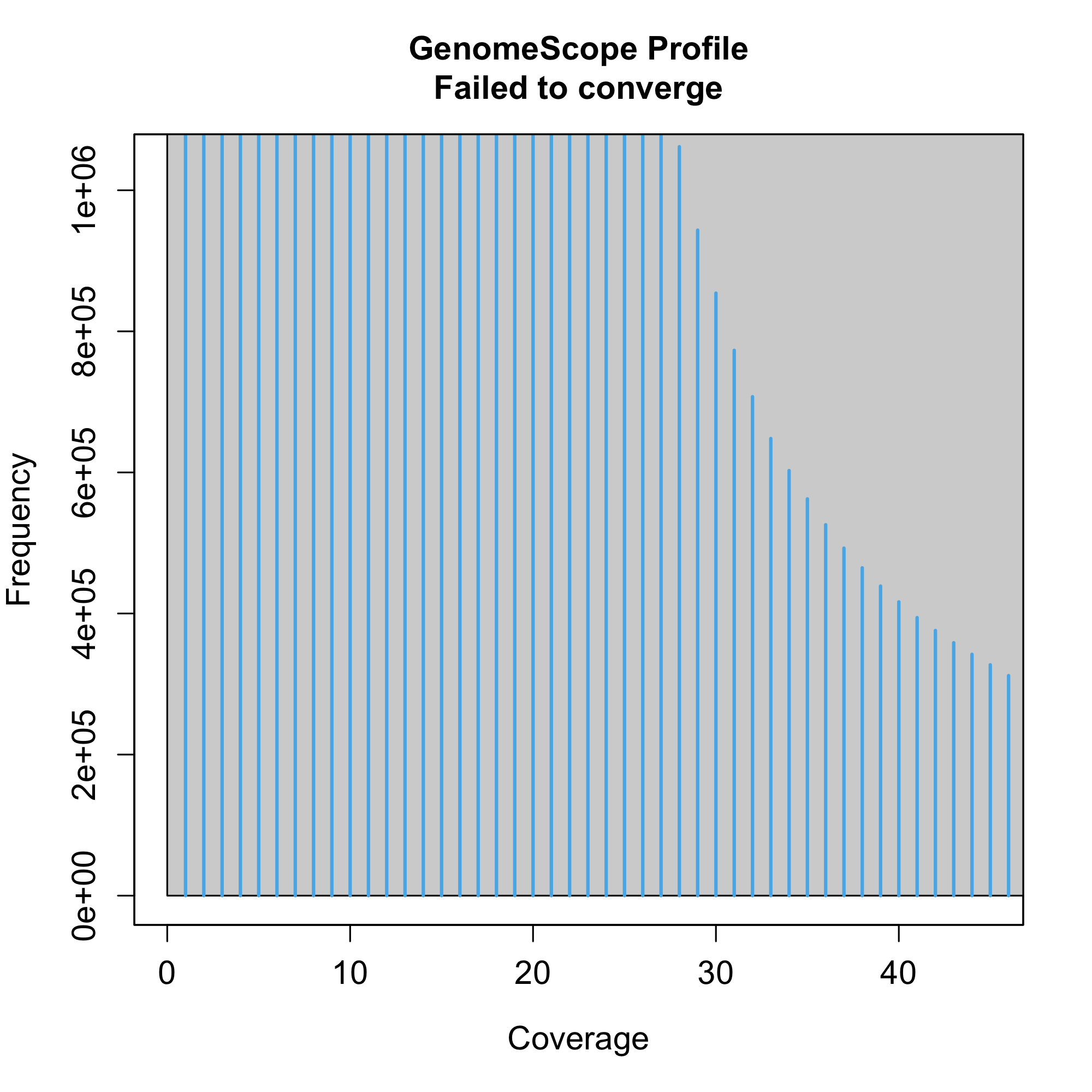

### plot.png

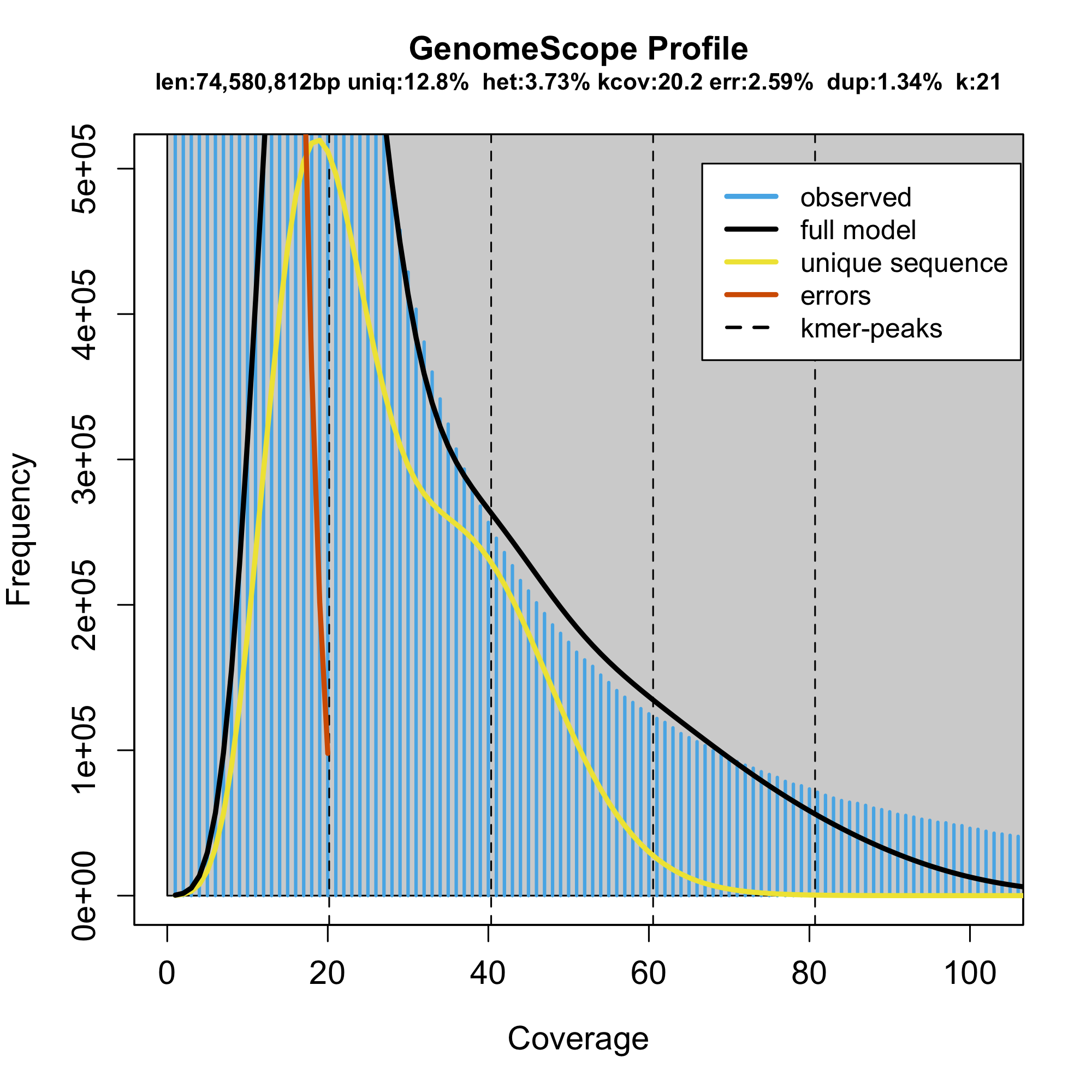

### plot.png

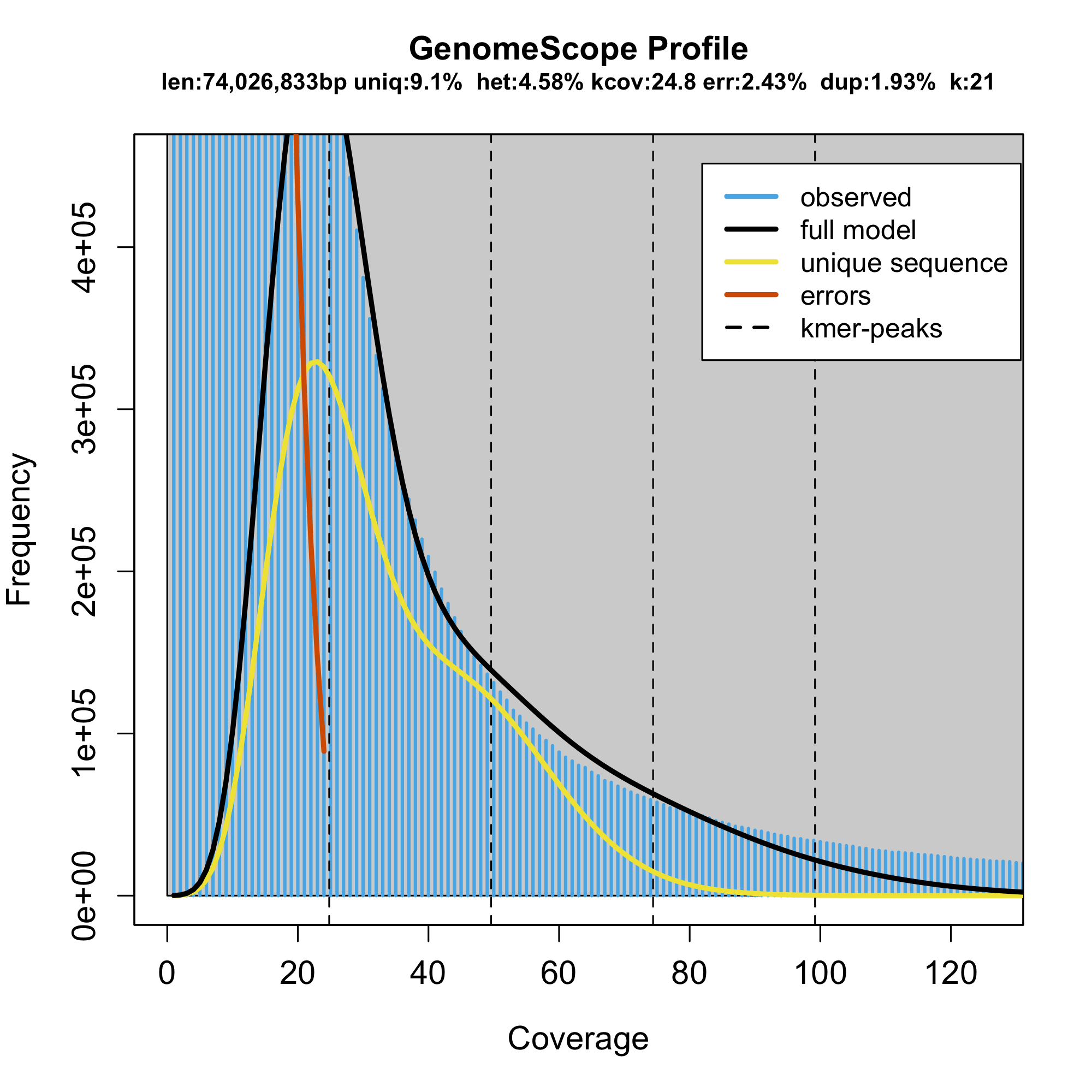

### plot.png

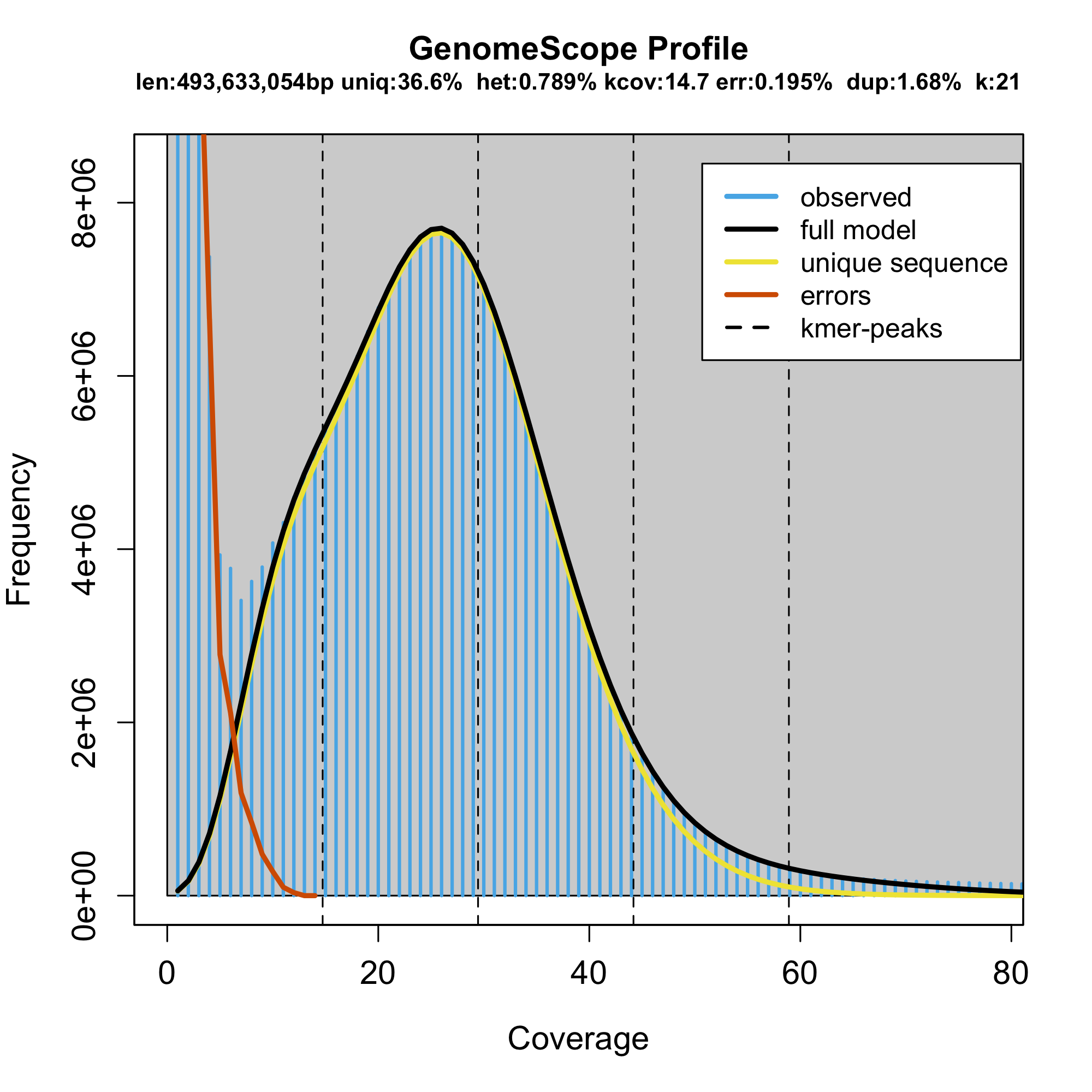

### plot.png

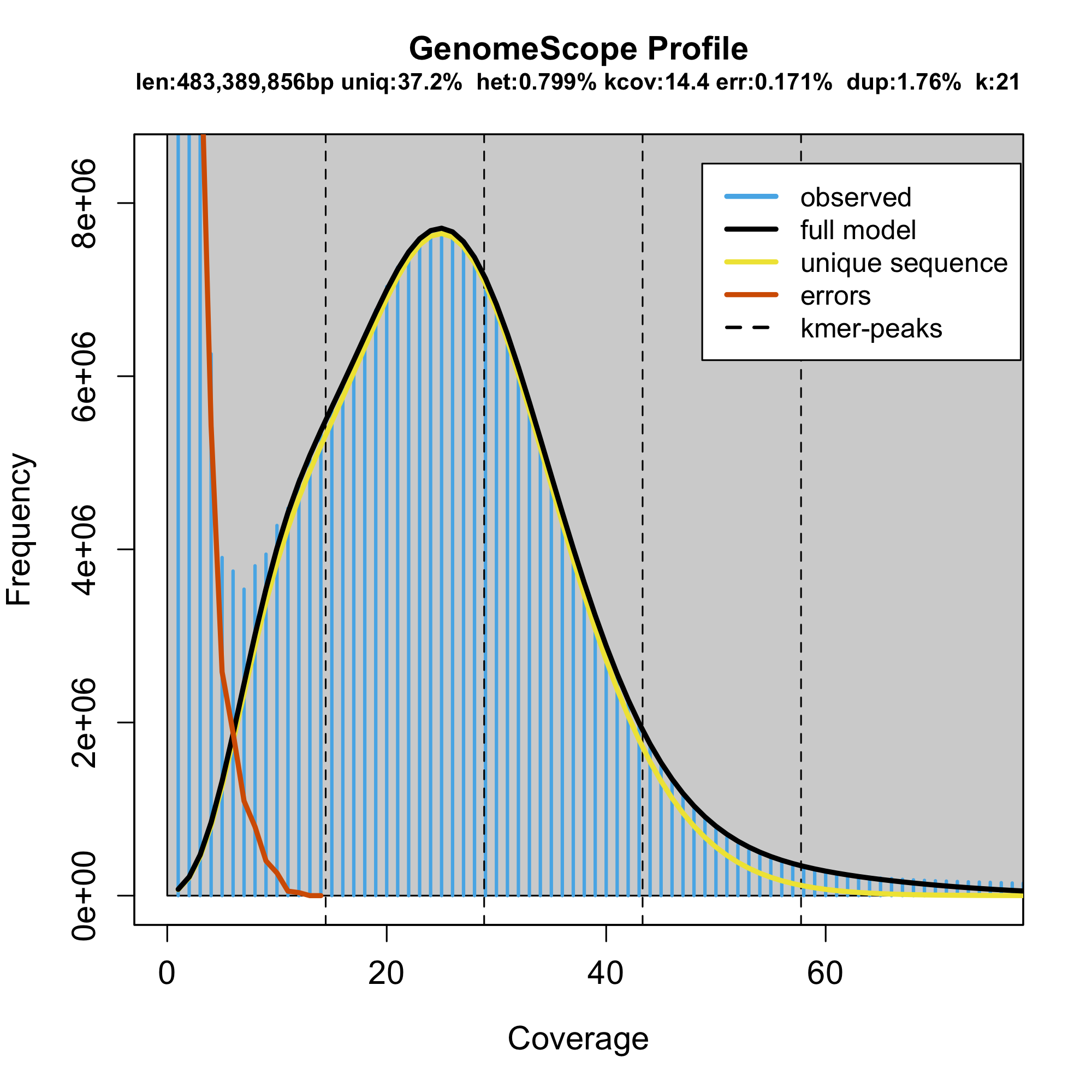

### plot.png

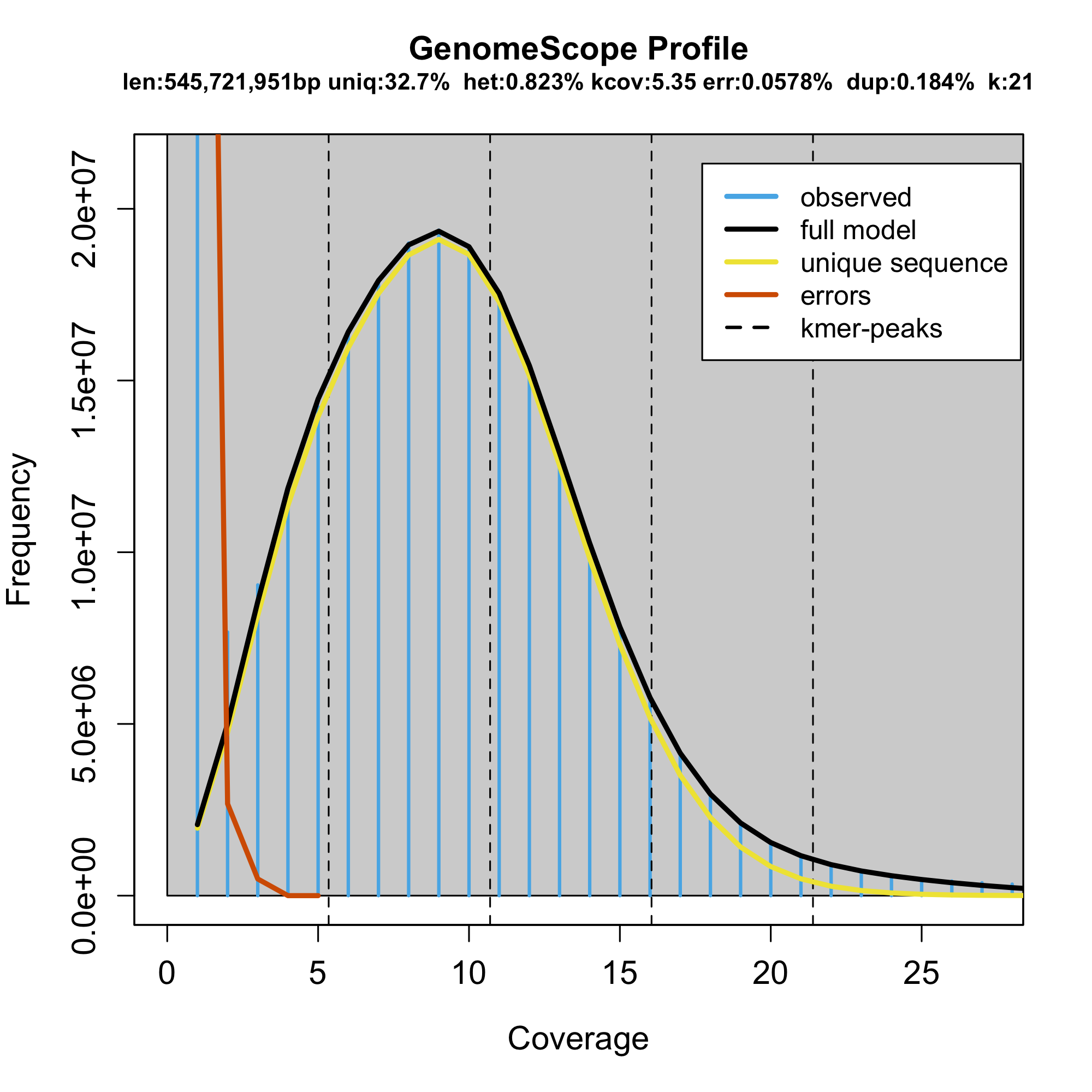

### plot.png

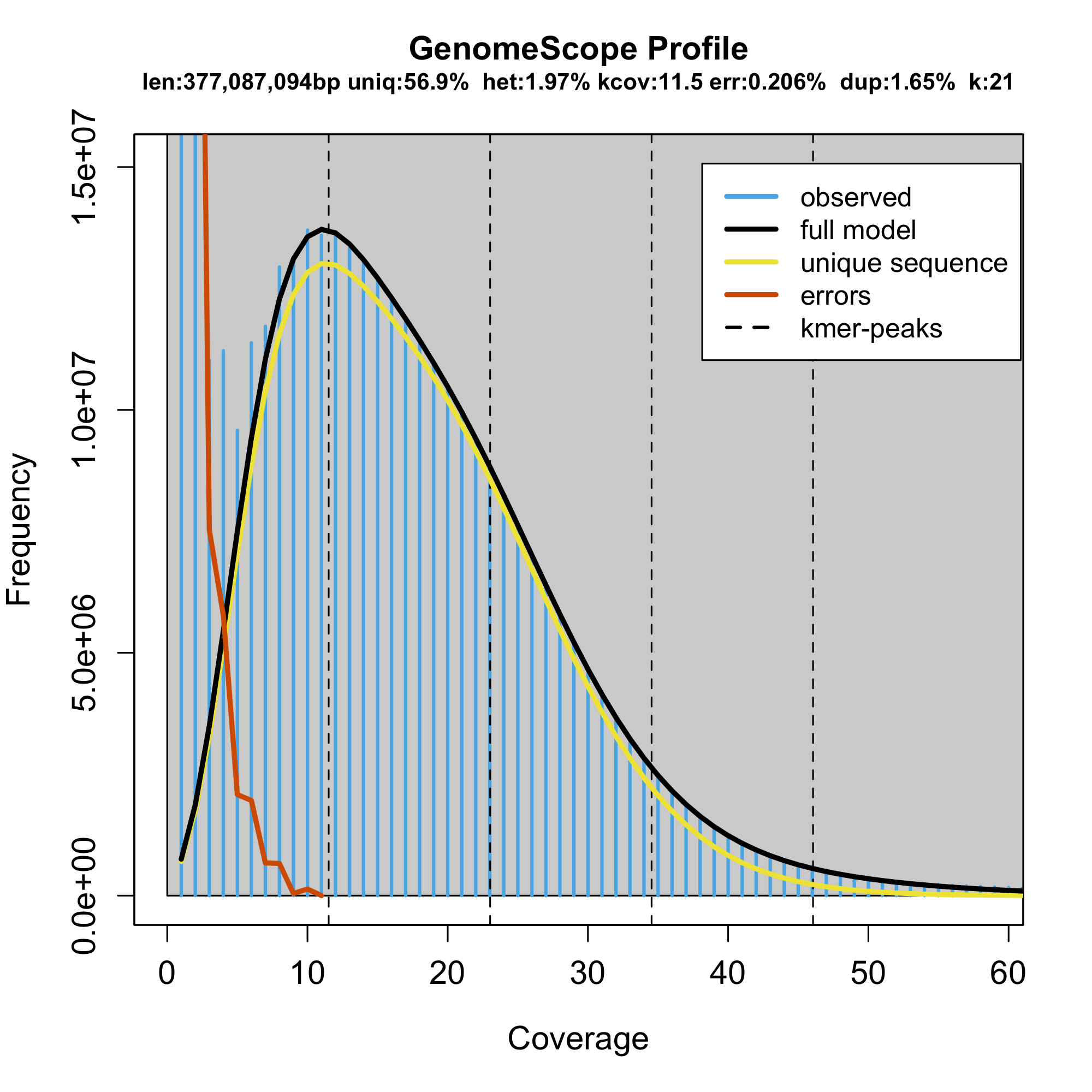

### plot.png

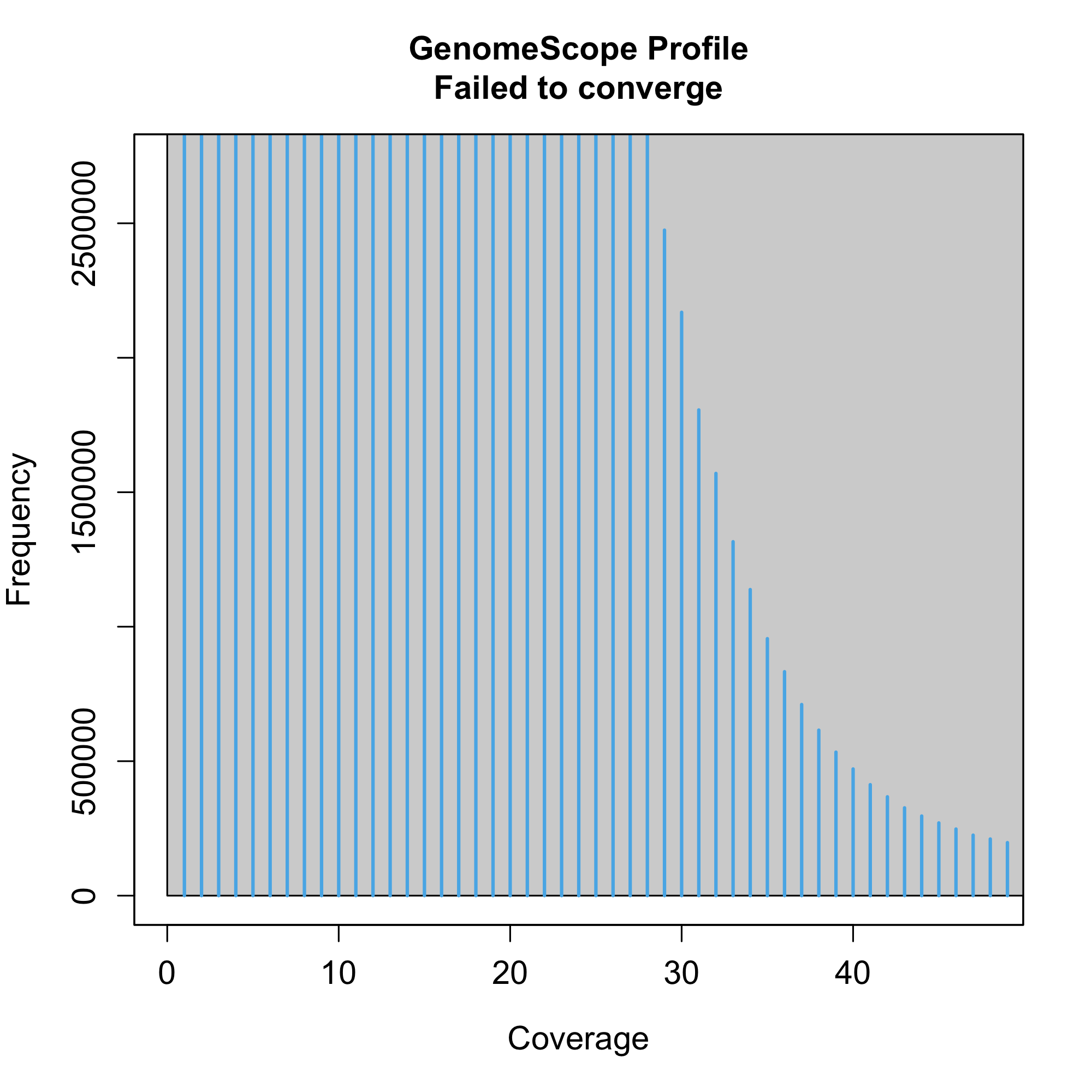

### plot.png

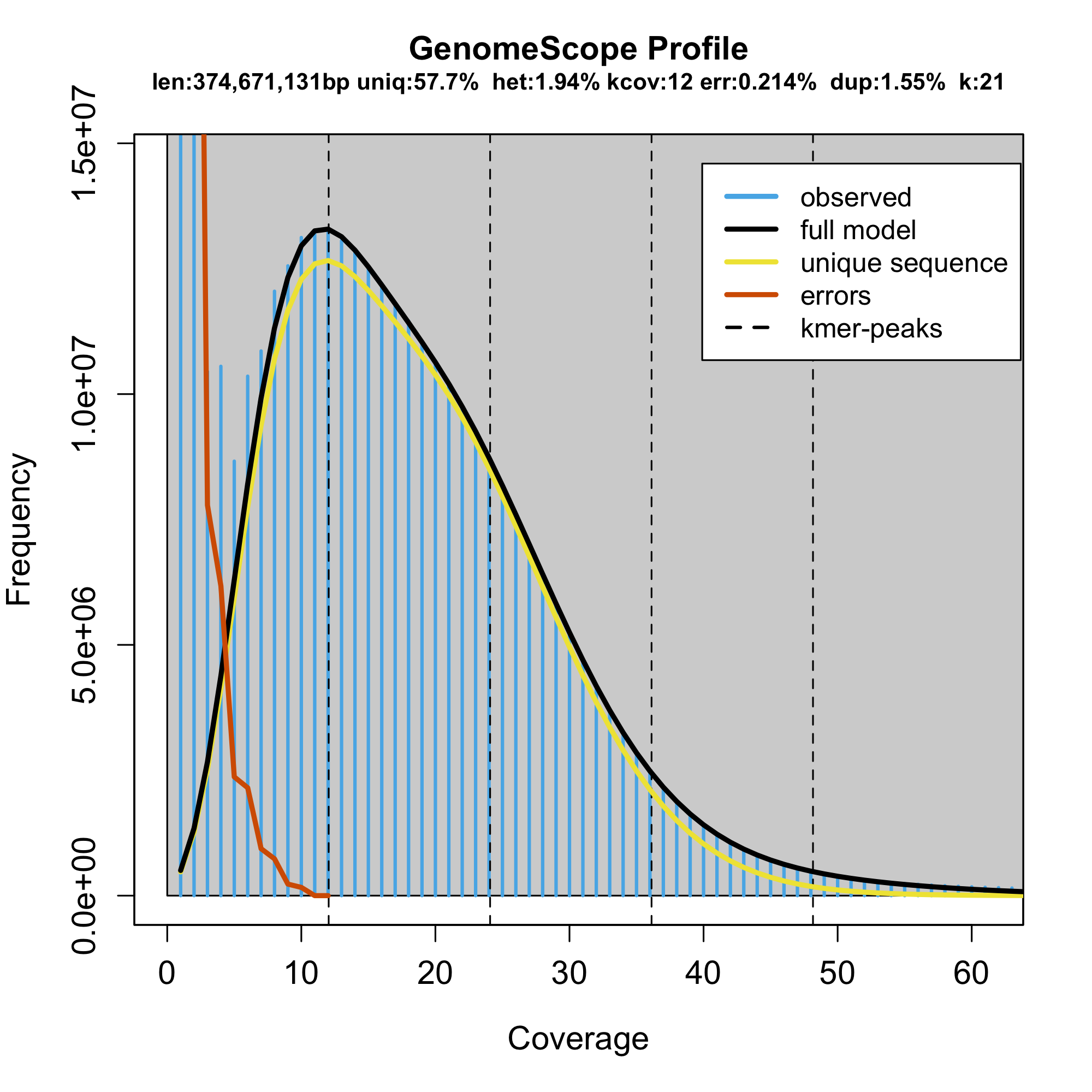

### plot.png

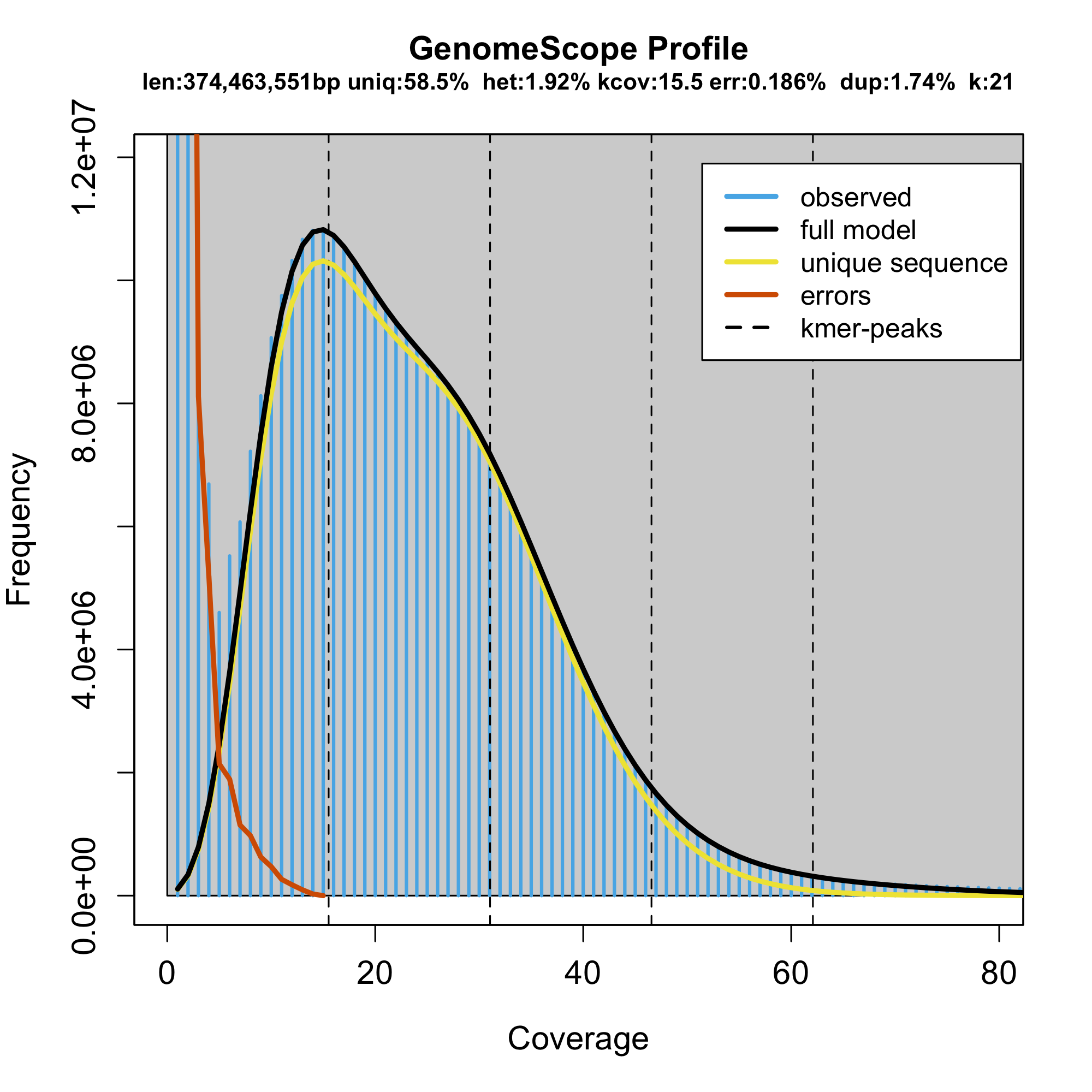

### plot.png

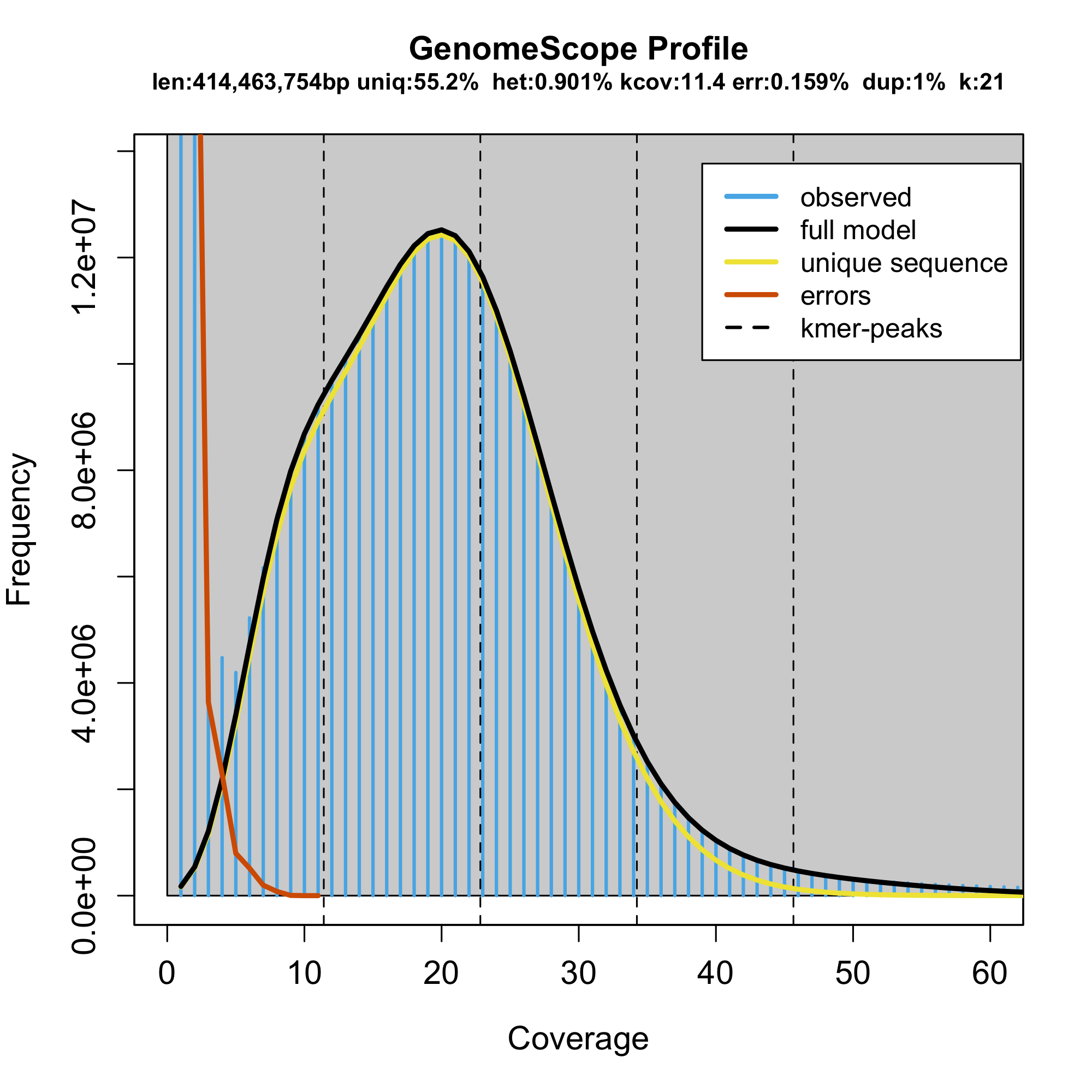

### plot.png

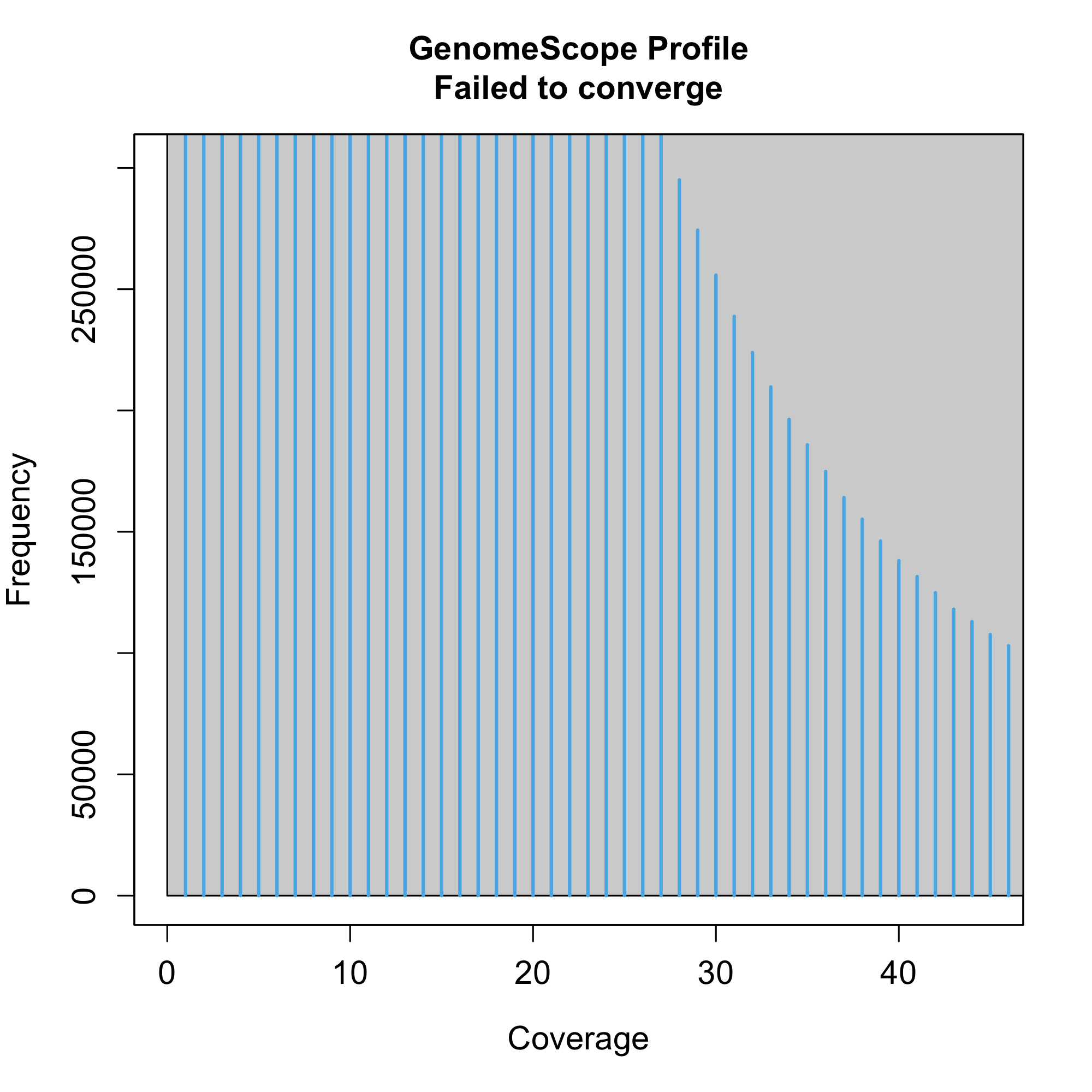

### plot.png

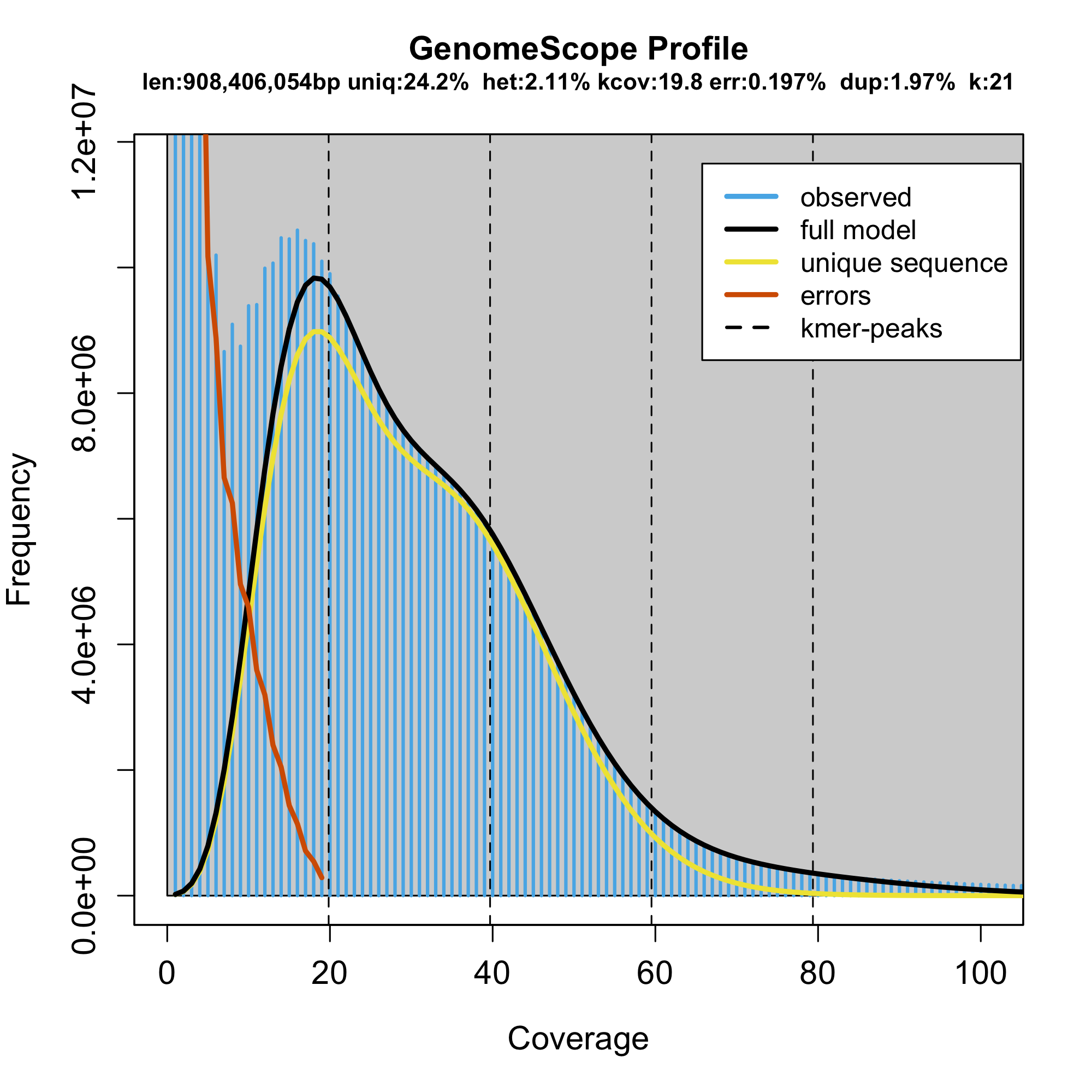

### plot.png

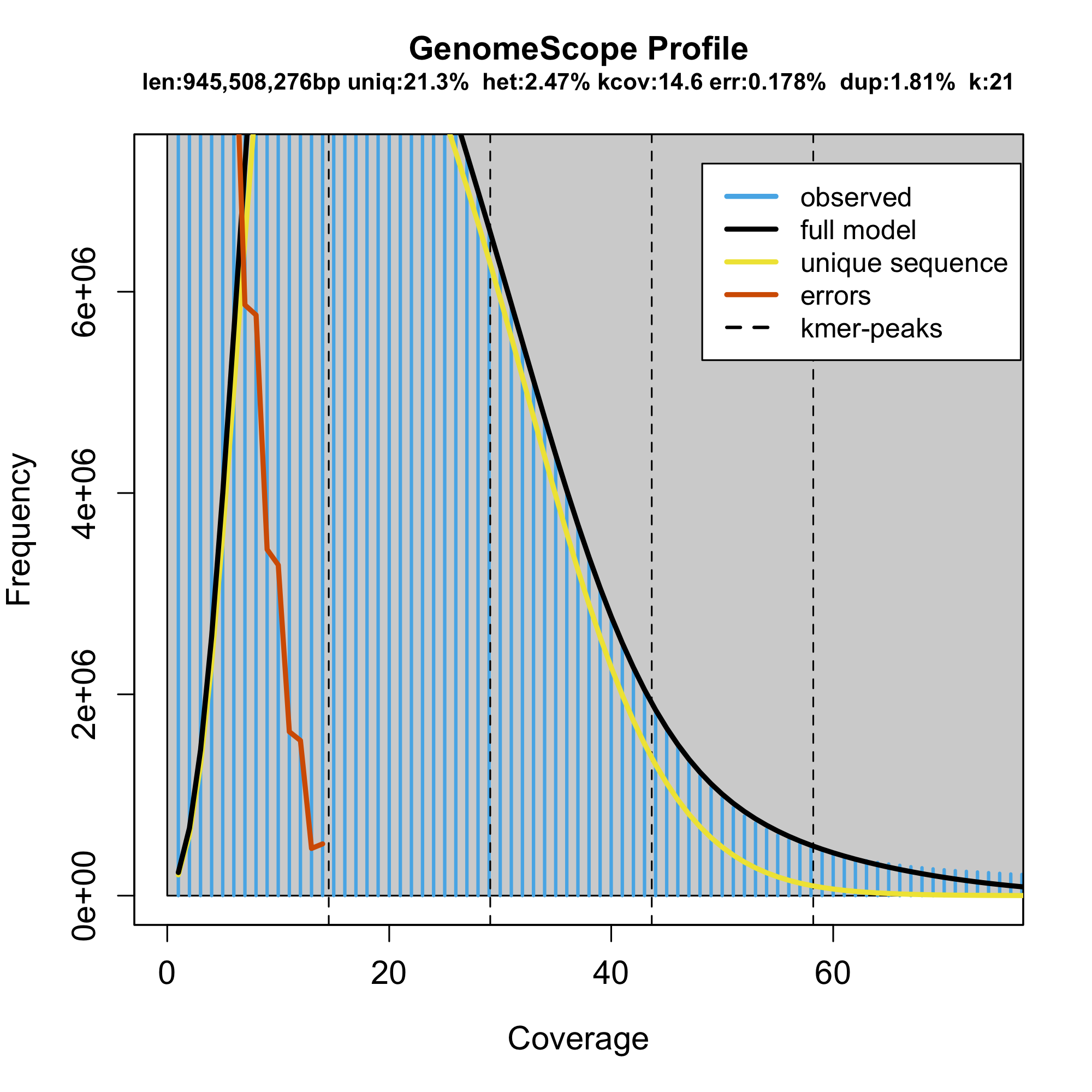

### plot.png

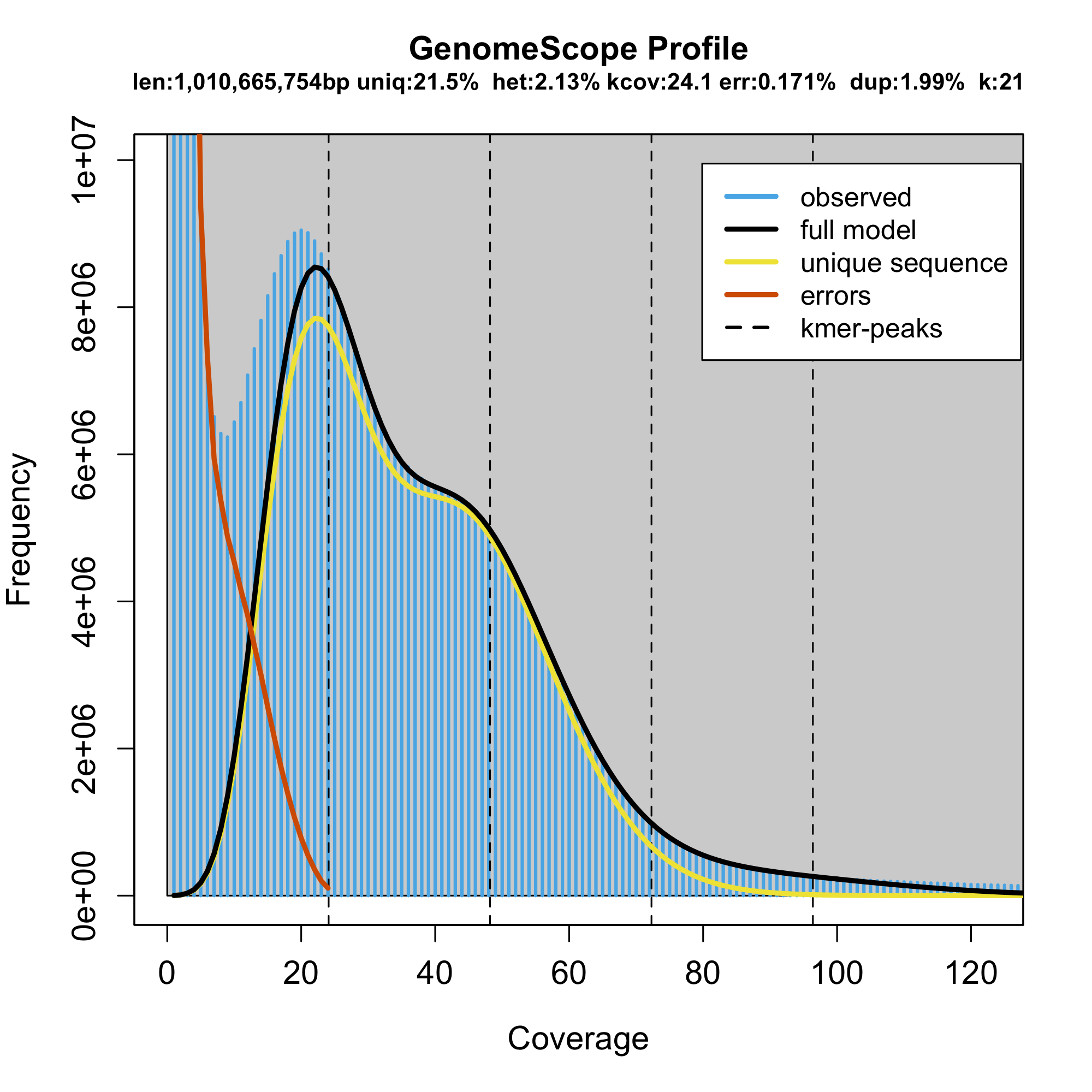

### plot.png

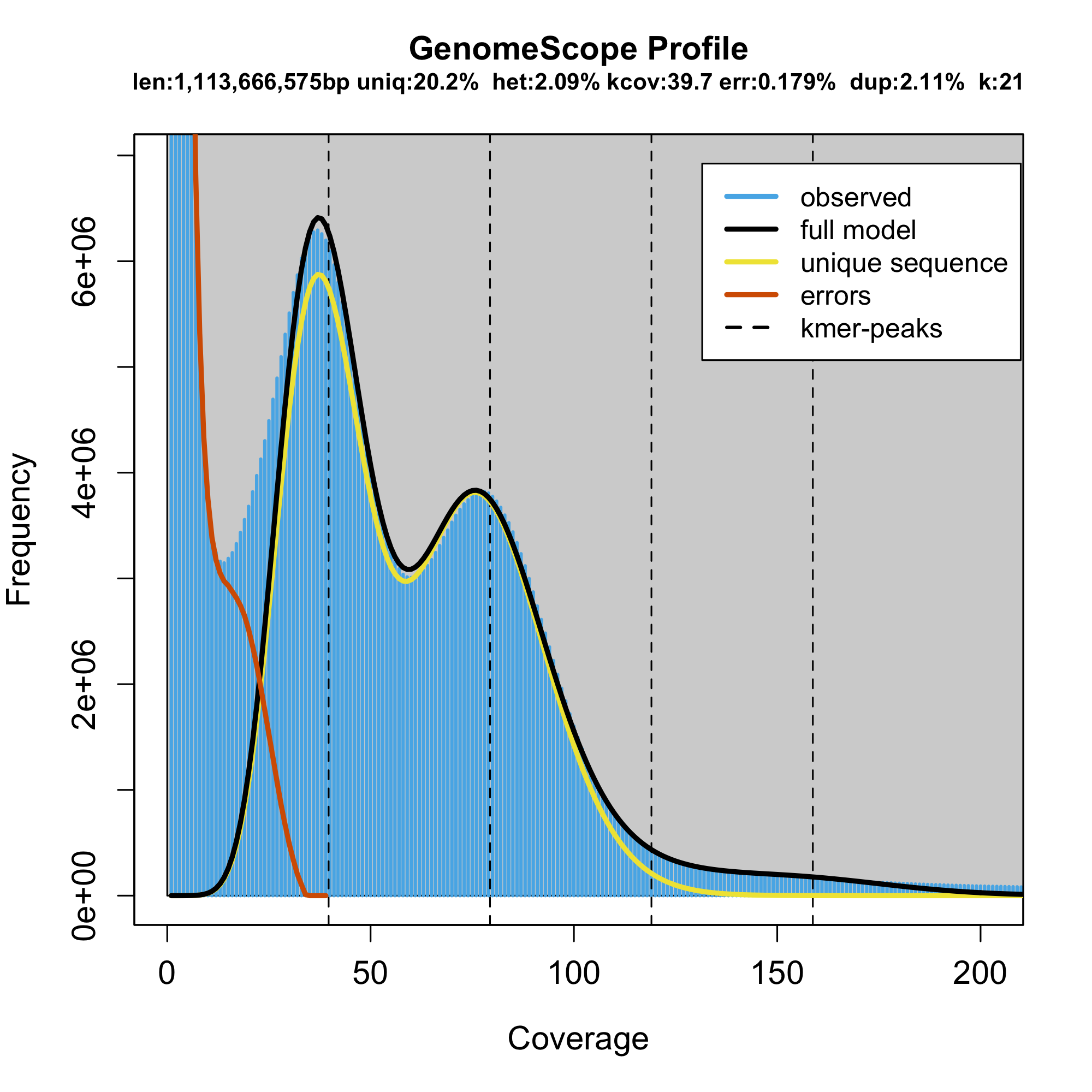

### plot.png

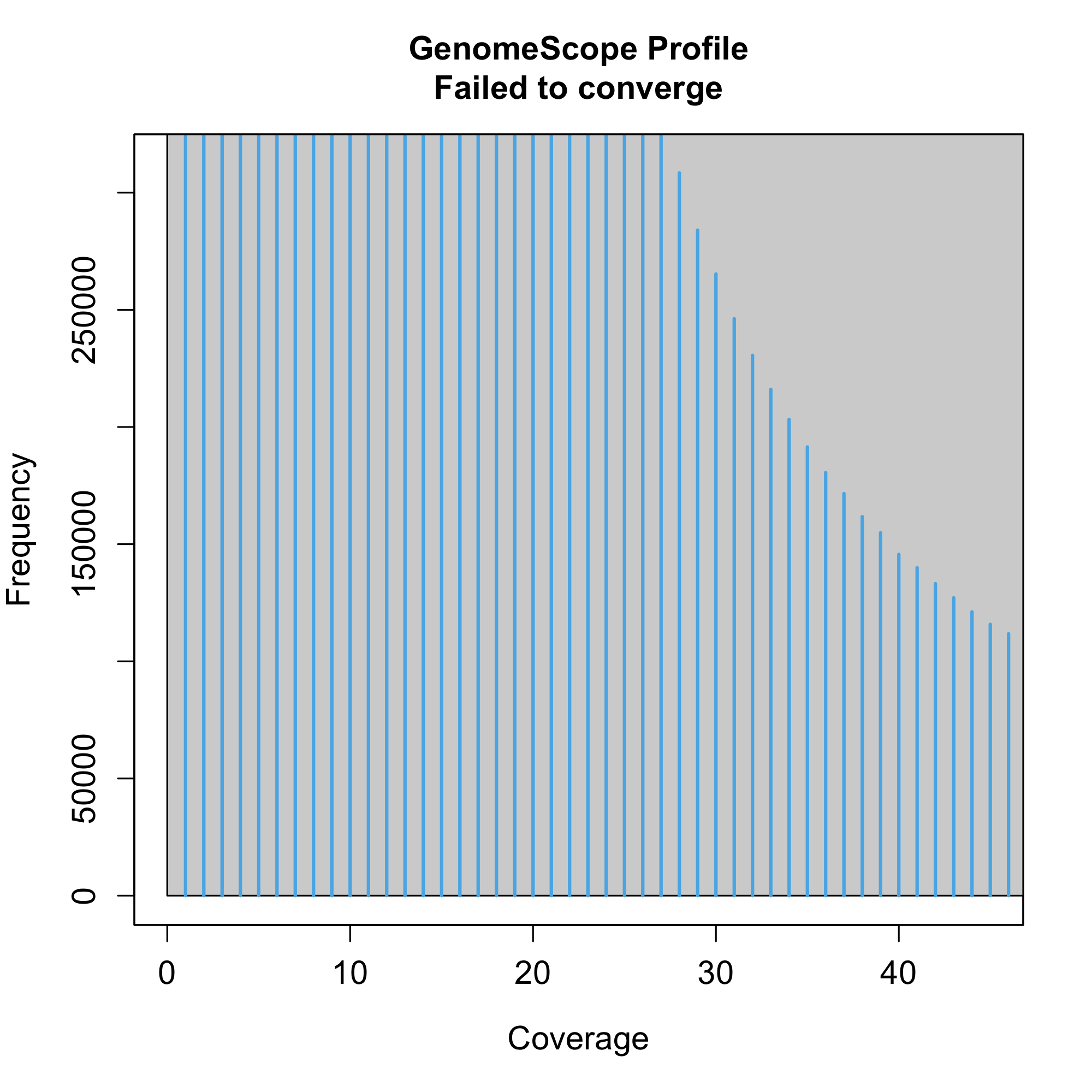

### Supplemental Figure 1

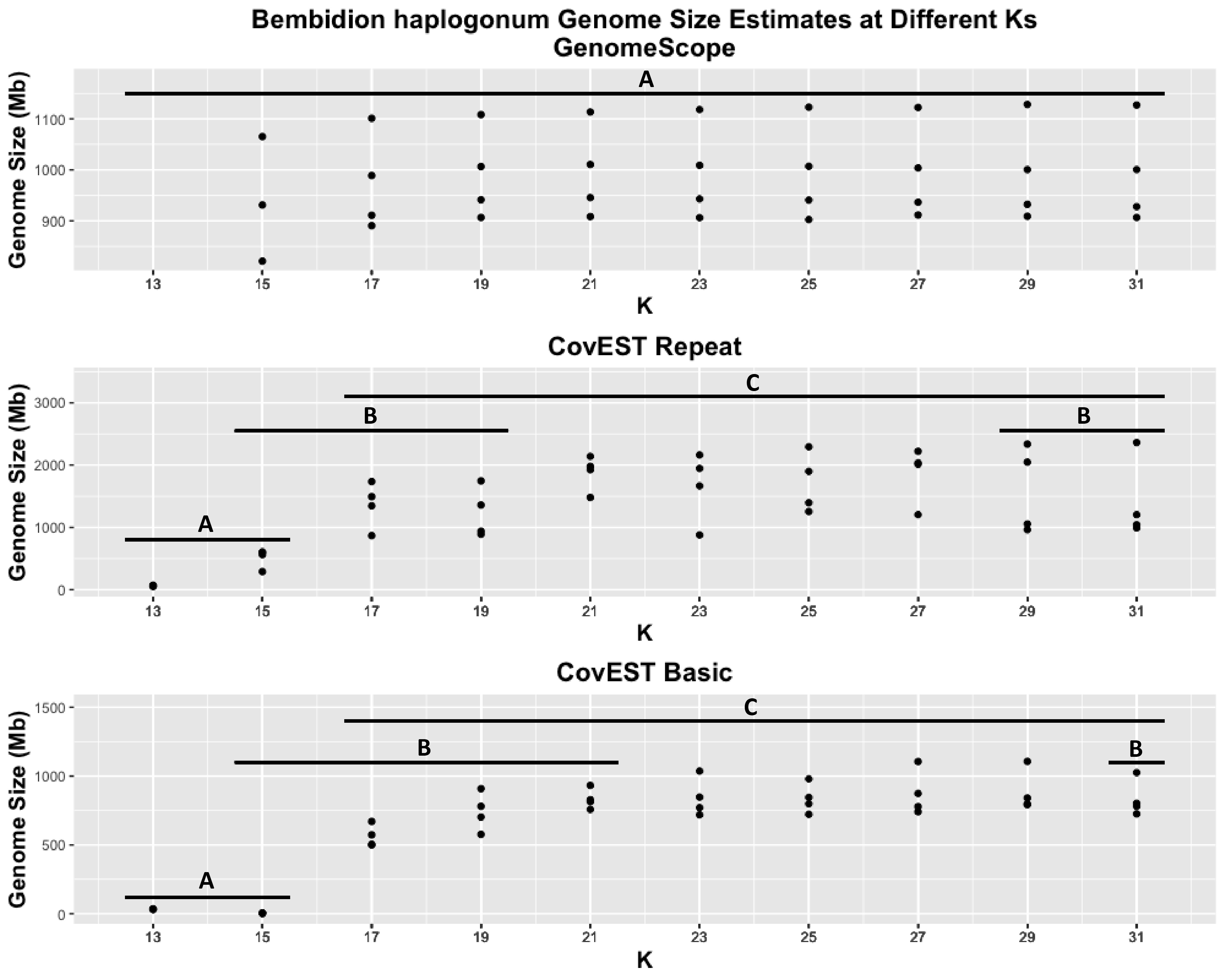

### Supplemental Figure 2

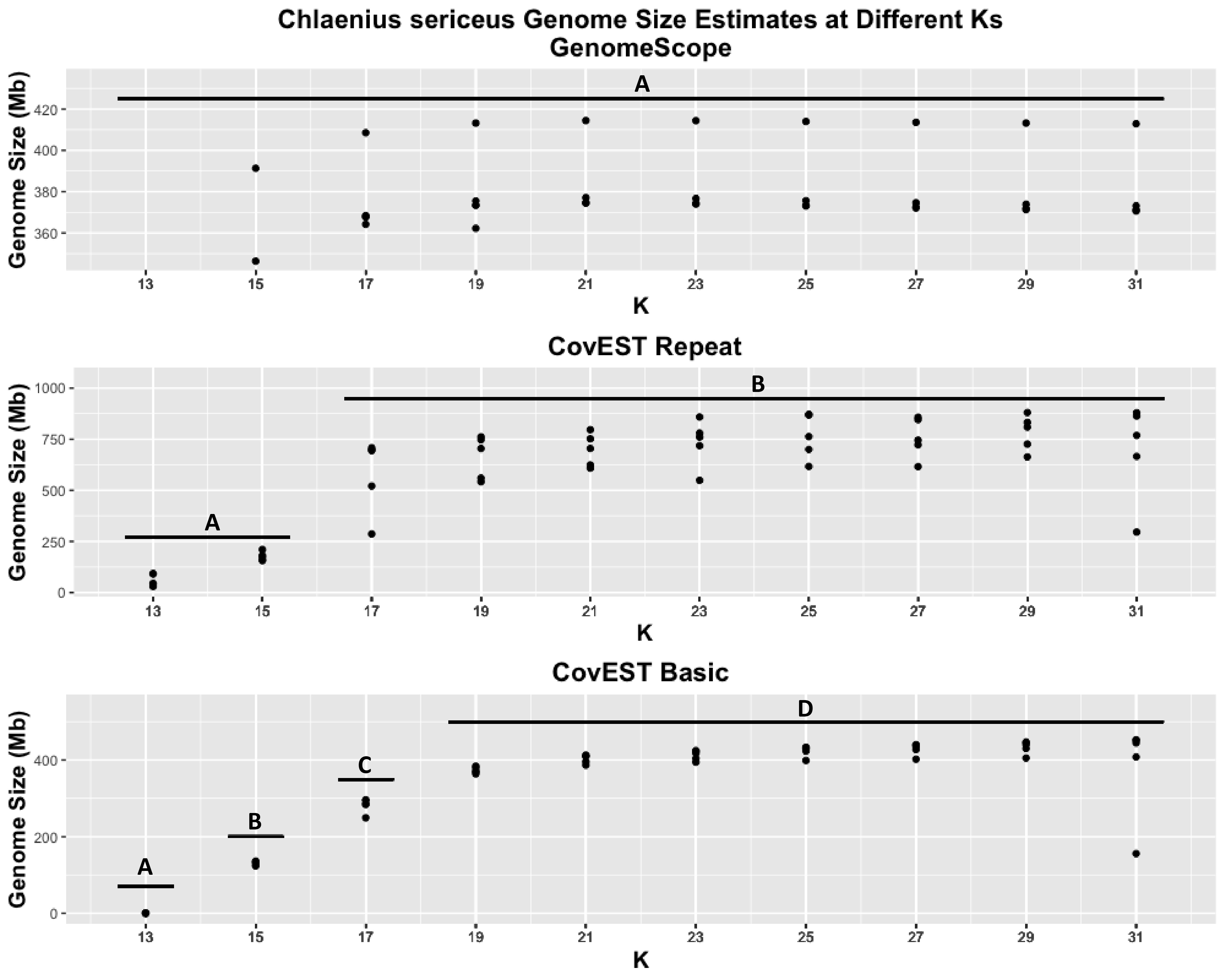

### Supplemental Figure 3

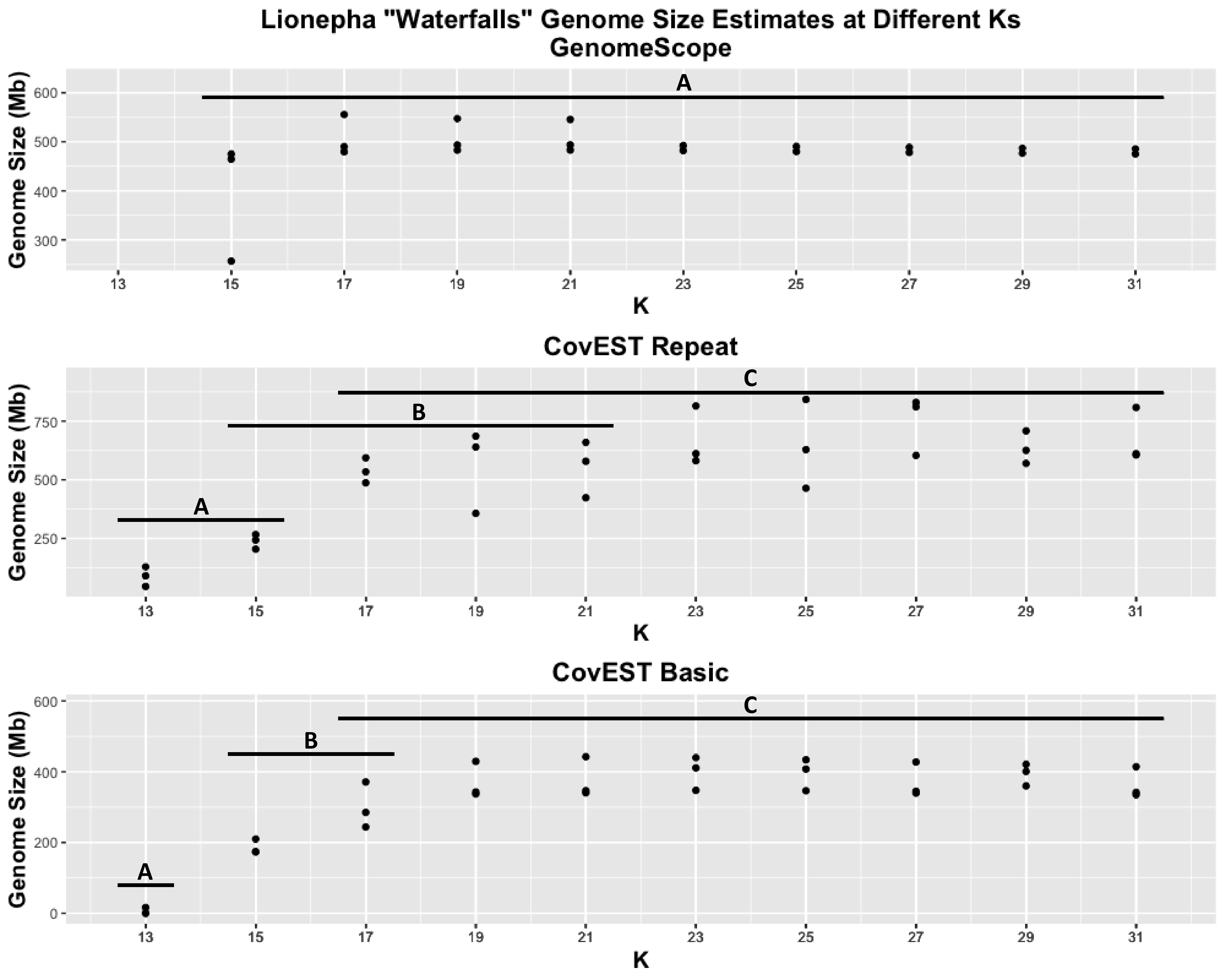

### Supplemental Figure 4

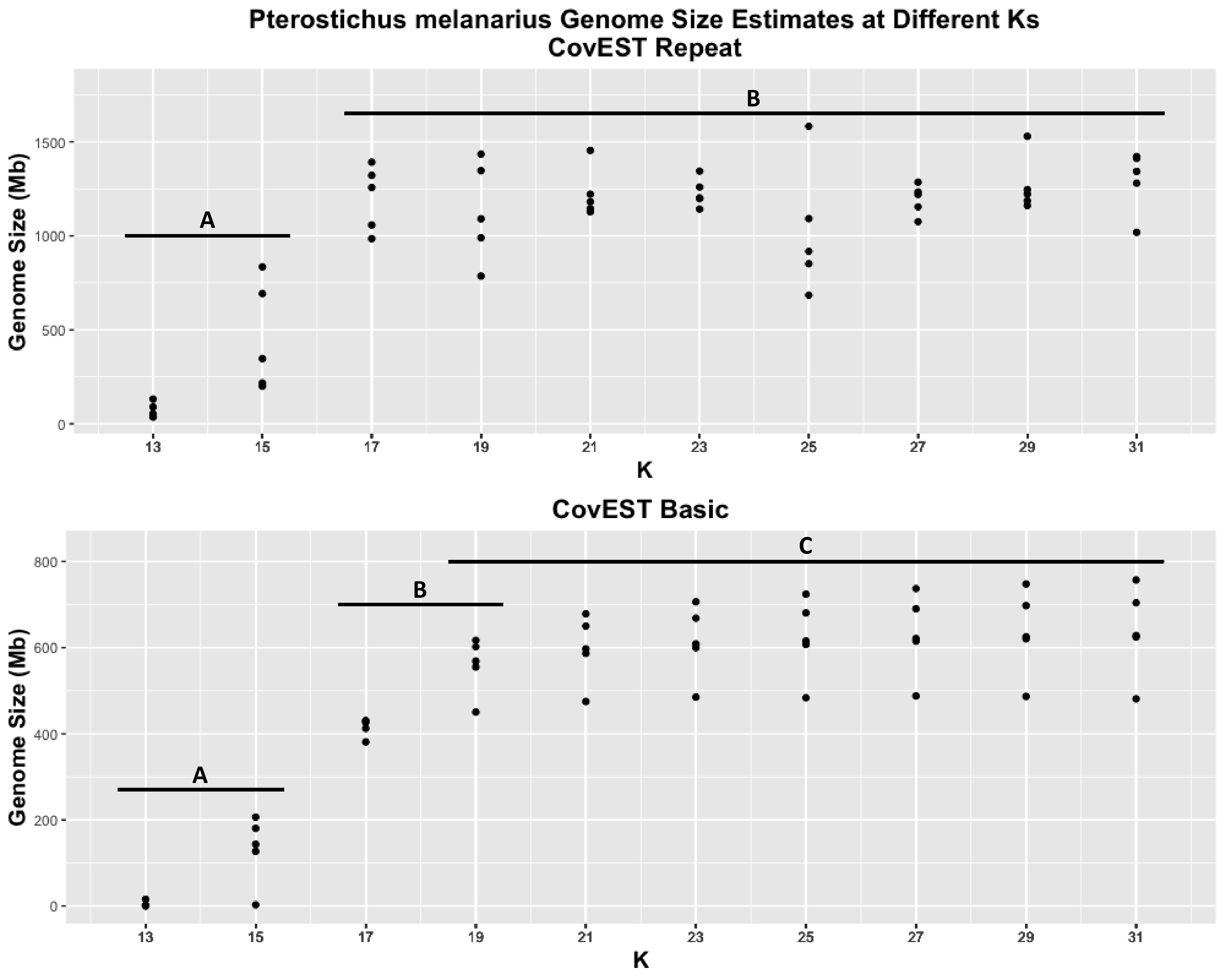

### Supplemental Figure 5

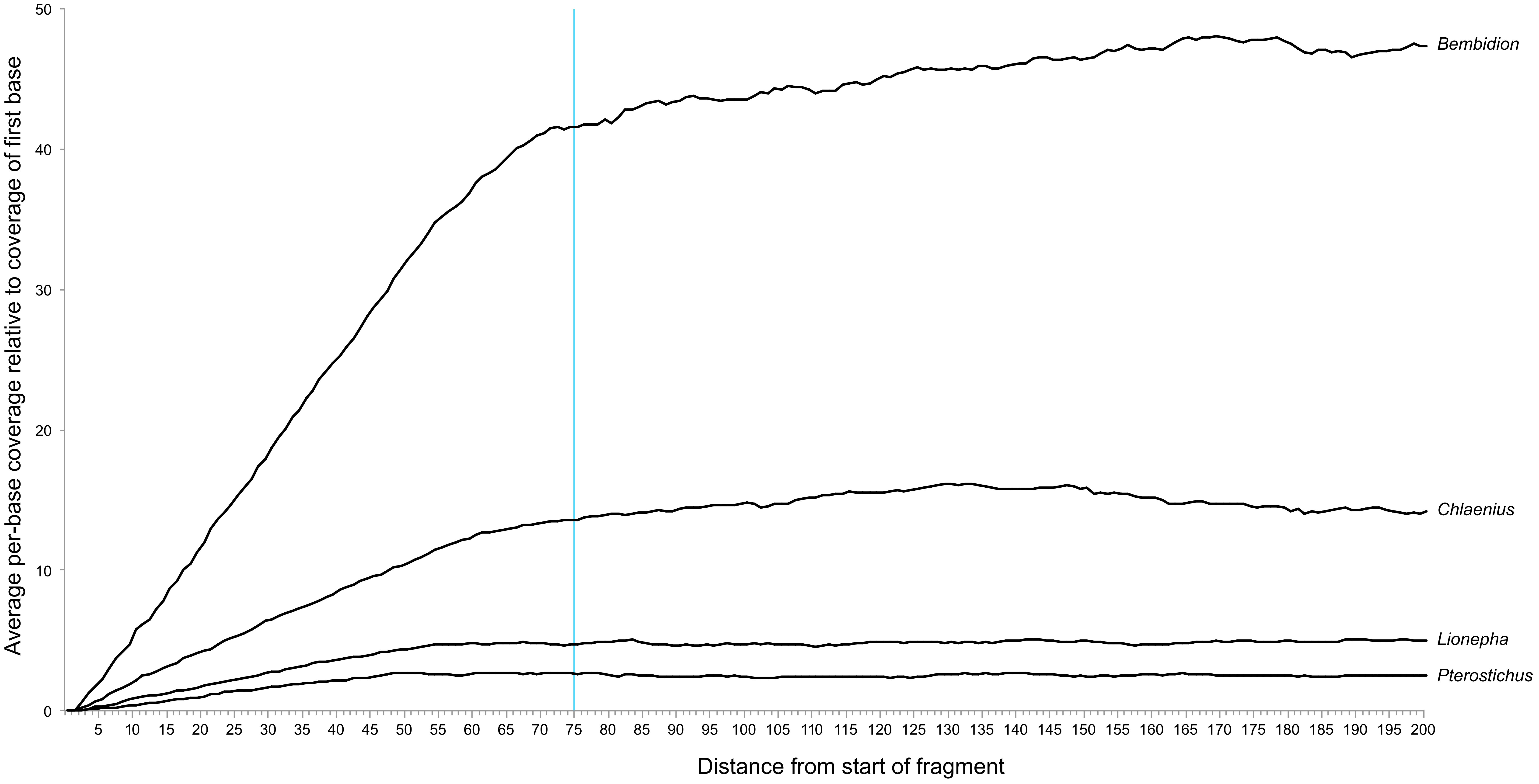

### Supplemental Figure 6

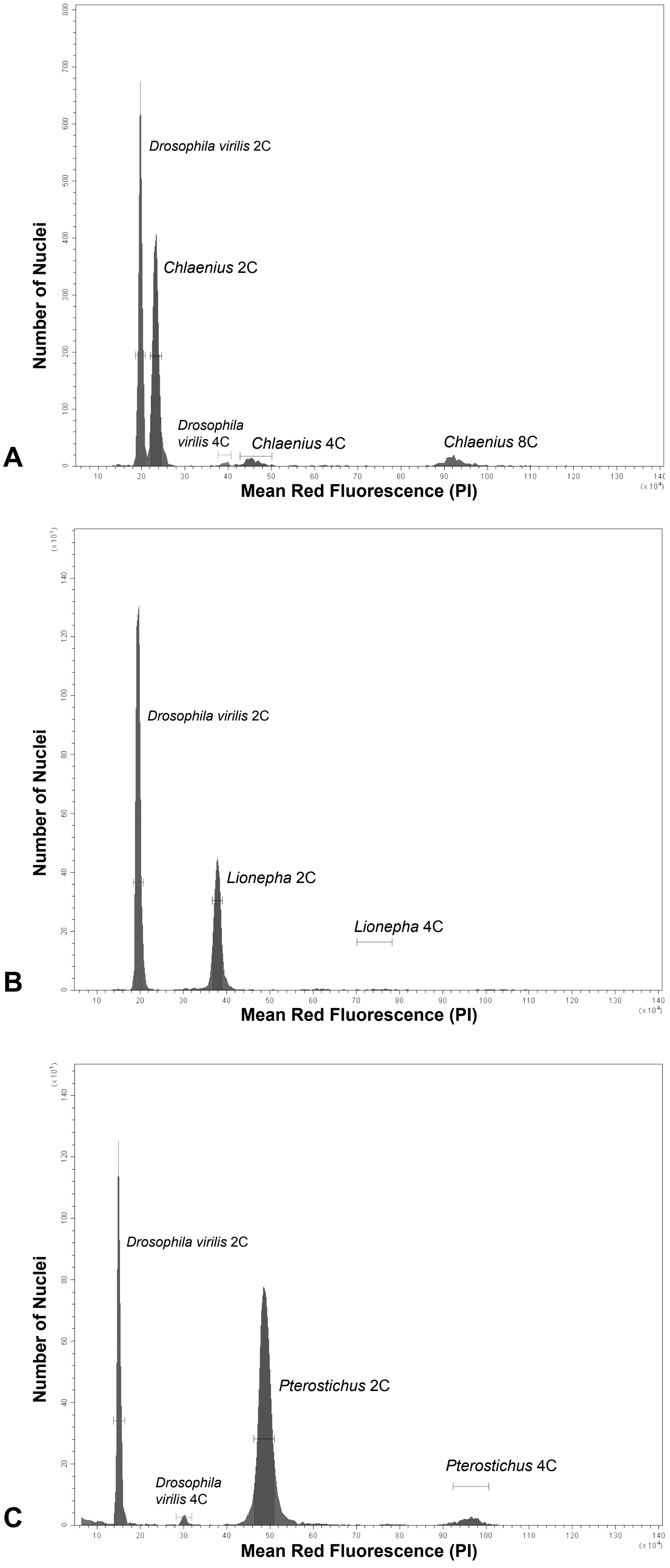

### Supplemental Figure 7

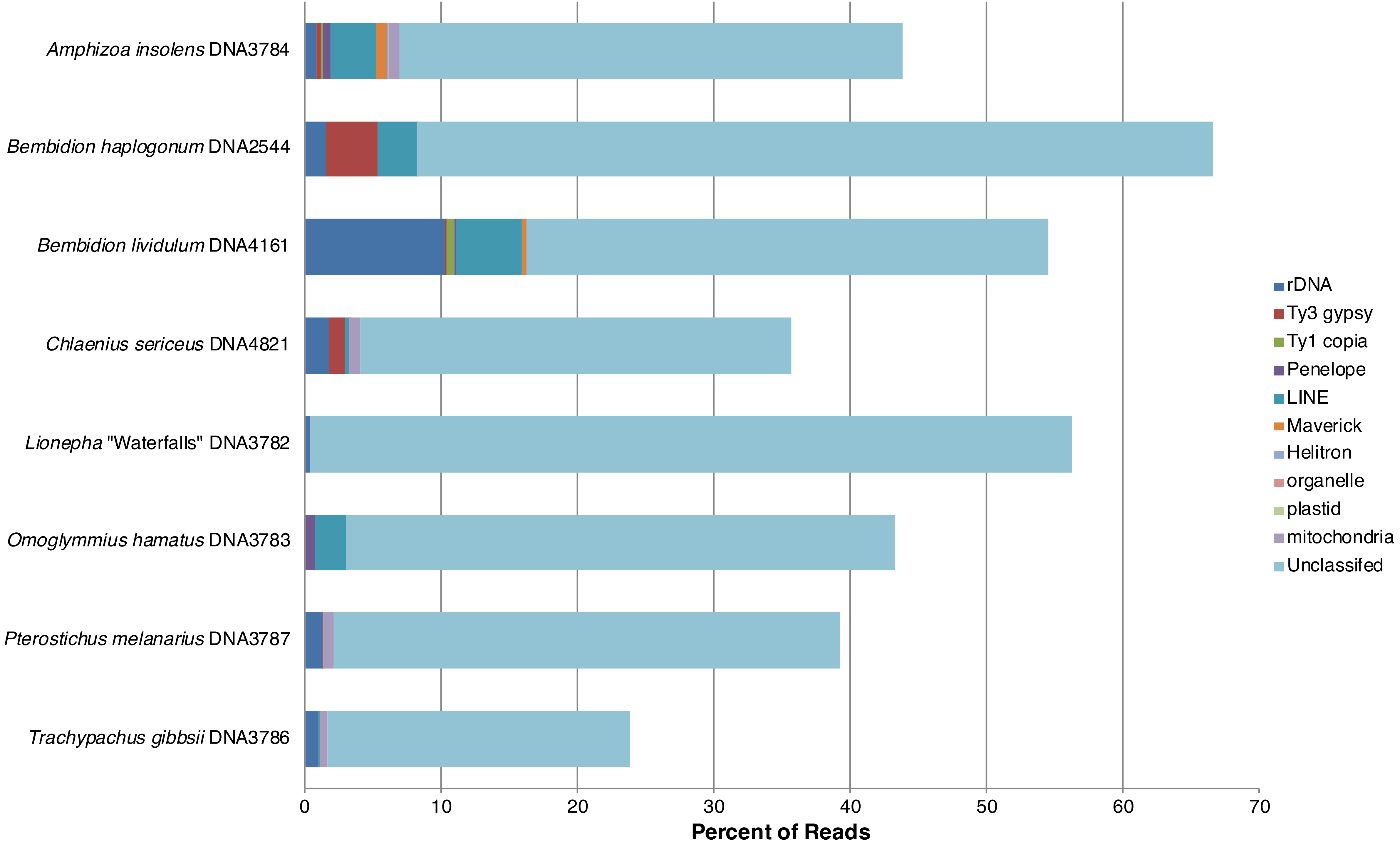
