## Supplemental Tables 1-14 for "Measuring genome sizes using read-depth, k-mers, and flow cytometry: methodological comparisons in beetles (Coleoptera)"

| **Sample Name** | **Sex** | **Collection Locality** | **Coordinates** | **Collection Date** | **SRA Accession Number** |
| --- | --- | --- | --- | --- | --- |
| *Amphizoa insolens* DNA3784 | Female | USA: Oregon: Sheep Creek | 44.4127 N 122.2092 W | 23-Aug-2013 | SRR8518617 |
| *Bembidion haplogonum* DNA2544 | Male | USA: Oregon: Corvallis | 44.5491 N 123.2449 W | 20-Sep-2009 | SRR8518612 |
| *Bembidion haplogonum* DNA5427 | Male | USA: Oregon: Corvallis | 44.5478 N 123.2430 W | 31-Aug-2018 | SRR8518625 |
| *Bembidion haplogonum* DNA5428 | Male | USA: Oregon: Corvallis | 44.5478 N 123.2430 W | 31-Aug-2018 | SRR8518626 |
| *Bembidion haplogonum* DNA5433 | Male | USA: Oregon: Corvallis | 44.5478 N 123.2430 W | 31-Aug-2018 | SRR8518631 |
| *Bembidion lividulum* DNA4161 | Male | USA: California: Ebbetts Pass | 38.5445 N 119.8115 W | 27-Jun-2014 | SRR8518619 |
| *Chlaenius sericeus* DNA4821 | Male | USA: California: Wilcox Ranch | 38.2497 N 121.8783 W | 29-Jun-2014 | SRR8518613 |
| *Chlaenius sericeus* JMP067 | Male | USA: Oregon: Prineville Reservoir | 44.1294 N 120.6971 W | 2-Aug-2018 | SRR8518630 |
| *Chlaenius sericeus* JMP068 | Male | USA: Oregon: Prineville Reservoir | 44.1294 N 120.6971 W | 2-Aug-2018 | SRR8518629 |
| *Chlaenius sericeus* JMP070 | Male | USA: Oregon: Prineville Reservoir | 44.1294 N 120.6971 W | 2-Aug-2018 | SRR8518623 |
| *Chlaenius sericeus* JMP071 | Male | USA: Oregon: Prineville Reservoir | 44.1294 N 120.6971 W | 2-Aug-2018 | SRR8518624 |
| *Lionepha* "Waterfalls" DNA3782 | Male | USA: Oregon: Marys Peak | 44.4746 N 123.5286 W | 4-Sep-2010 | SRR8518614 |
| *Lionepha* "Waterfalls" DNA5435^a^ | Male | USA: Oregon: Marys Peak | 44.4748 N 123.5280 W | 10-Oct-2018 | SRR8518632 |
| *Lionepha* "Waterfalls" DNA5436^a^ | Male | USA: Oregon: Marys Peak | 44.4748 N 123.5280 W | 10-Oct-2018 | SRR8518622 |
| *Omoglymmius hamatus* DNA3783 | Male | USA: California: Brightman Flat Cpgd | 38.3506 N 119.8451 W | 30-Jun-1996 | SRR8518618 |
| *Pterostichus melanarius* DNA3787 | Male | USA: Oregon: Corvallis | 44.5488 N 123.2482 W | 29-Aug-2013 | SRR8518615 |
| *Pterostichus melanarius* JMP059 | Male | USA: Oregon: Chip Ross Park | 44.6089 N 123.2813 W | 15-Jun-2018 | SRR8518620 |
| *Pterostichus melanarius* JMP060 | Male | USA: Oregon: Chip Ross Park | 44.6089 N 123.2813 W | 15-Jun-2018 | SRR8518621 |
| *Pterostichus melanarius* JMP061 | Male | USA: Oregon: Chip Ross Park | 44.6089 N 123.2813 W | 15-Jun-2018 | SRR8518627 |
| *Pterostichus melanarius* JMP062 | Male | USA: Oregon: Chip Ross Park | 44.6089 N 123.2813 W | 15-Jun-2018 | SRR8518628 |
| *Trachypachus gibbsii* DNA3786 | Male | USA: California: Carson Spur | 38.7047 N 120.1048 W | 1-Jun-2012 | SRR8518616 |

Table S1. Information on genomic specimens sequenced in this study.

a. Samples made with DNA extracted from different tissues of the same individual.

| **Sample Name** | **Genome  Size** | **Location** | **Coordinates** | **Collection Date** |
| --- | --- | --- | --- | --- |
| *Bembidion haplogonum* |  |  |  |  |
| Female |  |  |  |  |
| Bh F1 | 2190.61 | USA: Oregon: Corvallis | 44.5478 N 123.2430 W | 31-Aug-2018 |
| Bh F2 | 2190.54 | USA: Oregon: Corvallis | 44.5478 N 123.2430 W | 31-Aug-2018 |
| Bh F3 | 2217.67 | USA: Oregon: Corvallis | 44.5478 N 123.2430 W | 31-Aug-2018 |
| Bh F4 | 2138.03 | USA: Oregon: Corvallis | 44.5478 N 123.2430 W | 31-Aug-2018 |
| Bh F5 | 2208.21 | USA: Oregon: Corvallis | 44.5478 N 123.2430 W | 31-Aug-2018 |
| Bh F6 | 2186.22 | USA: Oregon: Corvallis | 44.5478 N 123.2430 W | 31-Aug-2018 |
| Bh F7 | 2228.84 | USA: Oregon: Corvallis | 44.5478 N 123.2430 W | 31-Aug-2018 |
| Bh F8 | 2212.52 | USA: Oregon: Corvallis | 44.5478 N 123.2430 W | 31-Aug-2018 |
| Bh F9 | 2168.05 | USA: Oregon: Corvallis | 44.5478 N 123.2430 W | 31-Aug-2018 |
| Male |  |  |  |  |
| Bh M1 | 2103.53 | USA: Oregon: Corvallis | 44.5478 N 123.2430 W | 31-Aug-2018 |
| Bh M2 | 2139.31 | USA: Oregon: Corvallis | 44.5478 N 123.2430 W | 31-Aug-2018 |
| Bh M3 | 2115.7 | USA: Oregon: Corvallis | 44.5478 N 123.2430 W | 31-Aug-2018 |
| Bh M4 | 2092.72 | USA: Oregon: Corvallis | 44.5478 N 123.2430 W | 31-Aug-2018 |
| Bh M5 | 2183.69 | USA: Oregon: Corvallis | 44.5478 N 123.2430 W | 31-Aug-2018 |
| Bh M6 | 2124.72 | USA: Oregon: Corvallis | 44.5478 N 123.2430 W | 31-Aug-2018 |
| Bh M7 | 2116.35 | USA: Oregon: Corvallis | 44.5478 N 123.2430 W | 31-Aug-2018 |
| Bh M8 | 2097.83 | USA: Oregon: Corvallis | 44.5478 N 123.2430 W | 31-Aug-2018 |
| Bh M9 | 2088.64 | USA: Oregon: Corvallis | 44.5478 N 123.2430 W | 31-Aug-2018 |
| *Lionepha* “Waterfalls” |  |  |  |  |
| Female |  |  |  |  |
| DRM V101282 | 597.6 | USA: Oregon: Marys Peak | 44.4748 N 123.5280 W | 10-Oct-2018 |
| Male |  |  |  |  |
| DRMV101276 | 584.62 | USA: Oregon: Marys Peak | 44.4748 N 123.5280 W | 10-Oct-2018 |
| DRMV101277 | 589.53 | USA: Oregon: Marys Peak | 44.4748 N 123.5280 W | 10-Oct-2018 |
| DRMV101278 | 593.98 | USA: Oregon: Marys Peak | 44.4748 N 123.5280 W | 10-Oct-2018 |
| DRMV101279 | 581.06 | USA: Oregon: Marys Peak | 44.4748 N 123.5280 W | 10-Oct-2018 |
| DRMV101280 | 579.33 | USA: Oregon: Marys Peak | 44.4748 N 123.5280 W | 10-Oct-2018 |
| *Bembidion lividulum* |  |  |  |  |
| Female |  |  |  |  |
| DRMV101289 | 837.39 | USA: Oregon: Creek nr Todd Lake | 44.0281 N 121.6711 W | 17-Jul-18 |
| Male |  |  |  |  |
| DRMV101284 | 835.03 | USA: Oregon: Creek nr Todd Lake | 44.0281 N 121.6711 W | 17-Jul-18 |
| DRMV101285 | 828.91 | USA: Oregon: Creek nr Todd Lake | 44.0281 N 121.6711 W | 17-Jul-18 |
| DRMV101286 | 822.42 | USA: Oregon: Creek nr Todd Lake | 44.0281 N 121.6711 W | 17-Jul-18 |
| DRMV101287 | 830.03 | USA: Oregon: Creek nr Todd Lake | 44.0281 N 121.6711 W | 17-Jul-18 |
| DRMV101288 | 842.09 | USA: Oregon: Creek nr Todd Lake | 44.0281 N 121.6711 W | 17-Jul-18 |
| *Pterostichus melanarius* |  |  |  |  |
| Female |  |  |  |  |
| Pm F2 | 1045.1 | USA: Oregon: Chip Ross Park | 44.6089 N 123.2813 W | 15-Jun-2018 |
| Pm F3 | 1018.3 | USA: Oregon: Chip Ross Park | 44.6089 N 123.2813 W | 15-Jun-2018 |
| Pm F4 | 1068 | USA: Oregon: Chip Ross Park | 44.6089 N 123.2813 W | 15-Jun-2018 |
| Pm F5 | 1031.2 | USA: Oregon: Chip Ross Park | 44.6089 N 123.2813 W | 15-Jun-2018 |
| Male |  |  |  |  |
| *Pterostichus melanarius* JMP059^a^ | 982.1 | USA: Oregon: Chip Ross Park | 44.6089 N 123.2813 W | 15-Jun-2018 |
| *Pterostichus melanarius* JMP060^a^ | 982.1 | USA: Oregon: Chip Ross Park | 44.6089 N 123.2813 W | 15-Jun-2018 |
| *Pterostichus melanarius* JMP061^a^ | 1014.7 | USA: Oregon: Chip Ross Park | 44.6089 N 123.2813 W | 15-Jun-2018 |
| *Pterostichus melanarius* JMP062^a^ | 1024.8 | USA: Oregon: Chip Ross Park | 44.6089 N 123.2813 W | 15-Jun-2018 |
| *Chlaenius sericeus* |  |  |  |  |
| Female |  |  |  |  |
| Cs F5 | 399.9 | USA: Oregon: Prineville Reservoir | 44.1294 N 120.6971 W | 2-Aug-2018 |
| Cs F6 | 414.1 | USA: Oregon: Prineville Reservoir | 44.1294 N 120.6971 W | 2-Aug-2018 |
| Cs F7 | 432.1 | USA: Oregon: Prineville Reservoir | 44.1294 N 120.6971 W | 2-Aug-2018 |
| Cs F8 | 409.3 | USA: Oregon: Prineville Reservoir | 44.1294 N 120.6971 W | 2-Aug-2018 |
| Cs F9 | 394.4 | USA: Oregon: Prineville Reservoir | 44.1294 N 120.6971 W | 2-Aug-2018 |
| Cs F10 | 400.4 | USA: Oregon: Prineville Reservoir | 44.1294 N 120.6971 W | 2-Aug-2018 |
| Male |  |  |  |  |
| *Chlaenius sericeus* JMP068^a^ | 385.81 | USA: Oregon: Prineville Reservoir | 44.1294 N 120.6971 W | 2-Aug-2018 |
| *Chlaenius sericeus* JMP069^a^ | 390.05 | USA: Oregon: Prineville Reservoir | 44.1294 N 120.6971 W | 2-Aug-2018 |
| *Chlaenius sericeus* JMP070^a^ | 396.53 | USA: Oregon: Prineville Reservoir | 44.1294 N 120.6971 W | 2-Aug-2018 |
| *Chlaenius sericeus* JMP071^a^ | 393.6 | USA: Oregon: Prineville Reservoir | 44.1294 N 120.6971 W | 2-Aug-2018 |

Table S2. Flow Cytometry values for specimens examined. Each value is the average of two technical replicates. Samples labeled with an asterisk were also analyzed by sequence-based methods.

a. Specimen also assessed with sequenced-based methods.

| **Sample Name** | **Sex** | **Collection Locality** | **Coordinates** | **Collection Date** | **SRA Accession Number** |
| --- | --- | --- | --- | --- | --- |
| *Bembidion haplogonum* DNA3229 | Male | USA: Oregon: Corvallis | 44.5491 N 123.2451 W | 28-Jul-2011 | SRR8801541 |
| *Bembidion lividulum* DNA4279 | Male | USA: California: Sonora Pass | 38.3322 N 119.6500 W | 26-Jun-2014 | SRR8801544 |
| *Chlaenius sericeus* JMPR007 | Male | USA: California: Wilcox Ranch | 38.2497 N 121.8783 W | 29-Jun-2014 | SRR8801540 |
| *Lionepha casta* DNA4602 | Male | USA: Oregon: Marys Peak | 44.4746 N 123.5286 W | 3-Sep-2011 | SRR8801542 |
| *Pterostichus melanarius* DNA4765 | Male | USA: Oregon: Corvallis | 44.5488 N 123.2482 W | 29-Aug-2013 | SRR8801543 |
| *Trachypachus gibbsii* DNA4436 | Male | USA: California: Carson Spur | 38.7047 N 120.1048 W | 1-Jun-2012 | SRR8801545 |

Table S3. Information on transcriptomic specimens sequenced in this study.

|  | **Sample Preservation** | **Library Prep Kit** | **Illumina Platform** | **Read Length** |
| --- | --- | --- | --- | --- |
| *Bembidion* *haplogonum* DNA3229 | Fresh | TruSeq RNA | HiSeq 2000 | 50 |
| *Bembidion* *lividulum* DNA4279 | RNAlater@ -20C | NEBNext RNA Ultra II | HiSeq 2500 | 101 |
| *Chlaenius* *sericeus* JMPR007 | RNAlater @ -20C | NEBNext RNA Ultra II | HiSeq 2500 | 101 |
| *Lionepha* *casta* DNA4602 | RNAlater @ -20C | NEBNext RNA Ultra II | HiSeq 2500 | 100 |
| *Pterostichus* *melanarius* DNA4765 | Fresh | NEBNext RNA Ultra II | HiSeq 2500 | 100 |
| *Trachypachus* *gibbsii* DNA4436 | RNAlater @ -20C | NEBNext RNA Ultra II | HiSeq 2500 | 101 |

Table S4. Additional details for methods used on transcriptomic sequencing specimens. All extractions were performed using a homogenate of the entire body.

|  | **Sample Preservation** | **Tissue** | **Library Prep Kit** | **Illumina**  **Platform** | **Read Length** |
| --- | --- | --- | --- | --- | --- |
| *Amphizoa* *insolens* DNA3784 | EtOH @ -20C | Muscle | Illumina TruSeq | HiSeq 2500 | 101 |
| *Bembidion haplogonum* DNA2544 | EtOH @ -20C | Reproductive | NEBNext DNA Ultra II | HiSeq 3000 | 151 |
| *Bembidion* *haplogonum* DNA5427 | Frozen, EtOH @ -20C | Reproductive | NEBNext DNA Ultra II | HiSeq 3000 | 151 |
| *Bembidion* *haplogonum* DNA5428 | Frozen, EtOH @ -20C | Muscle | NEBNext DNA Ultra II | HiSeq 3000 | 151 |
| *Bembidion* *haplogonum* DNA5433 | Frozen, EtOH @ -20C | Reproductive | NEBNext DNA Ultra II | HiSeq 3000 | 151 |
| *Bembidion* *lividulum* DNA4161 | EtOH @ -20C | Whole body | NEBNext DNA Ultra II | HiSeq 2500 | 101 |
| *Chlaenius sericeus* DNA4821 | EtOH @ -20C | Muscle | NEBNext DNA Ultra II | HiSeq 3000 | 151 |
| *Chlaenius* *sericeus* JMP068 | Frozen, EtOH @ -20C | Muscle | NEBNext DNA Ultra II | HiSeq 3000 | 151 |
| *Chlaenius* *sericeus* JMP069 | Frozen, EtOH @ -20C | Muscle | NEBNext DNA Ultra II | HiSeq 3000 | 151 |
| *Chlaenius* *sericeus* JMP070 | Frozen, EtOH @ -20C | Muscle | NEBNext DNA Ultra II | HiSeq 3000 | 151 |
| *Chlaenius* *sericeus* JMP071 | Frozen, EtOH @ -20C | Muscle | NEBNext DNA Ultra II | HiSeq 3000 | 151 |
| *Lionepha* "Waterfalls" DNA3782 | EtOH @ -20C | Muscle | Illumina TruSeq | HiSeq 2500 | 101 |
| *Lionepha* "Waterfalls" DNA5435^a^ | Frozen, EtOH @ -20C | Reproductive | NEBNext DNA Ultra II | HiSeq 3000 | 151 |
| *Lionepha* "Waterfalls" DNA5436^a^ | Frozen, EtOH @ -20C | Muscle | NEBNext DNA Ultra II | HiSeq 3000 | 151 |
| *Omoglymmius* *hamatus* DNA3783 | EtOH @ -20C | Muscle | Illumina TruSeq | HiSeq 2500 | 101 |
| *Pterostichus* *melanarius* DNA3787 | EtOH @ -20C | Muscle | Illumina TruSeq | HiSeq 2500 | 101 |
| *Pterostichus* *melanarius* JMP059 | Frozen, EtOH @ -20C | Muscle | NEBNext DNA Ultra II | HiSeq 3000 | 151 |
| *Pterostichus* *melanarius* JMP060 | Frozen, EtOH @ -20C | Muscle | NEBNext DNA Ultra II | HiSeq 3000 | 151 |
| *Pterostichus* *melanarius* JMP061 | Frozen, EtOH @ -20C | Muscle | NEBNext DNA Ultra II | HiSeq 3000 | 151 |
| *Pterostichus* *melanarius* JMP062 | Frozen, EtOH @ -20C | Muscle | NEBNext DNA Ultra II | HiSeq 3000 | 151 |
| *Trachypachus* *gibbsii* DNA3786 | EtOH @ -20C | Muscle | Illumina TruSeq | HiSeq 2500 | 101 |

Table S5. Additional details for methods used on genomic sequencing specimens.

a. Samples made with DNA extracted from different tissues of the same individual.

|  | **Summary** | **Complete** | **Fragmented** | **Missing** |
| --- | --- | --- | --- | --- |
| *Amphizoa insolens* DNA3784 | C:34.7%[S:34.5%,D:0.2%],F:31.2%,M:34.1%,n:2442 | 848 | 761 | 833 |
| *Bembidion haplogonum* DNA2544 | C:59.3%[S:57.5%,D:1.8%],F:27.2%,M:13.5%,n:2442 | 1,446 | 664 | 332 |
| *Bembidion lividulum* DNA4161 | C:29.8%[S:29.6%,D:0.2%],F:35.4%,M:34.8%,n:2442 | 726 | 865 | 851 |
| *Chlaenius sericeus* DNA4821 | C:78.7%[S:78.2%,D:0.5%],F:15.6%,M:5.7%,n:2442 | 1,923 | 380 | 139 |
| *Lionepha* "Waterfalls" DNA3782 | C:68.5%[S:68.3%,D:0.2%],F:20.1%,M:11.4%,n:2442 | 1,673 | 491 | 278 |
| *Omoglymmius hamatus* DNA3783 | C:7.3%[S:7.3%,D:0.0%],F:28.3%,M:64.4%,n:2442 | 178 | 690 | 1,574 |
| *Pterostichus melanarius* DNA3787 | C:30.7%[S:30.7%,D:0.0%],F:33.5%,M:35.8%,n:2442 | 750 | 817 | 875 |
| *Trachypachus gibbsii* DNA3786 | C:45.8%[S:45.2%,D:0.6%],F:31.5%,M:22.7%,n:2442 | 1,119 | 770 | 553 |

Table S6. Results of BUSCO analysis on eight genome assemblies using the 2442 gene Endopterygota odb9 reference set. Summary column shows results in BUSCO notation: C:complete [S:single-copy, D:duplicated], F:fragmented, M:missing, n:number of genes used. Genes classified as “complete” have lengths within two standard deviations of the BUSCO group mean length. “Complete” genes found only once are classified as “single-copy”, while those that are found multiple times are classified as “duplicated.” Partially recovered genes are classified as “fragmented,” and genes not recovered at all are classified as “missing.”

|  | **Summary** | **Complete** | **Fragmented** | **Missing** |
| --- | --- | --- | --- | --- |
| *Bembidion haplogonum* DNA3229 | C:85.0%[S:50.8%,D:34.2%],F:10.8%,M:4.2%,n:2442 | 2077 | 264 | 101 |
| *Bembidion lividulum* DNA4279 | C:79.5%[S:61.1%,D:18.4%],F:11.6%,M:8.9%,n:2442 | 1941 | 283 | 218 |
| *Chlaenius sericeus* JMPR007 | C:81.5%[S:64.6%,D:16.9%],F:9.6%,M:8.9%,n:2442 | 1990 | 234 | 218 |
| *Lionepha casta* DNA4602 | C:84.4%[S:57.5%,D:26.9%],F:10.5%,M:5.1%,n:2442 | 2063 | 257 | 122 |
| *Pterostichus melanarius* DNA4765 | C:90.1%[S:53.8%,D:36.3%],F:5.1%,M:4.8%,n:2442 | 2201 | 125 | 116 |
| *Trachypachus gibbsii* DNA4436 | C:87.1%[S:61.5%,D:25.6%],F:8.0%,M:4.9%,n:2442 | 2127 | 196 | 119 |

Table S7. Results of BUSCO analysis on six transcriptome assemblies using the 2442 gene Endopterygota odb9 reference set. See caption of Table TX03 for additional explanation.

|  | **Genome Size** | **Repeat** | **Unique** | **Error min** | **Error max** | **Fit Min** | **Fit Max** |
| --- | --- | --- | --- | --- | --- | --- | --- |
| *Amphizoa insolens* DNA3784 | - | - | - | 0% | 0% | 0% | 0% |
| *Bembidion haplogonum* DNA2544 | 1,113.67 | 888.97 | 224.70 | 0.18% | 0.18% | 90.29% | 97.26% |
| *Bembidion haplogonum* DNA5427 | 1,010.67 | 793.81 | 216.85 | 0.17% | 0.17% | 91.22% | 98.97% |
| *Bembidion haplogonum* DNA5428 | 945.51 | 744.29 | 201.21 | 0.18% | 0.18% | 91.16% | 99.06% |
| *Bembidion haplogonum* DNA5433 | 908.41 | 688.79 | 219.61 | 0.20% | 0.20% | 92.10% | 99.23% |
| *Bembidion lividulum* DNA4161 | - | - | - | 0% | 0% | 0% | 0% |
| *Chlaenius sericeus* DNA4821 | 414.46 | 185.88 | 228.58 | 0.16% | 0.16% | 96.68% | 99.73% |
| *Chlaenius sericeus* JMP068 | 374.46 | 155.32 | 219.14 | 0.19% | 0.19% | 96.57% | 99.64% |
| *Chlaenius sericeus* JMP069 | 374.67 | 158.34 | 216.33 | 0.21% | 0.21% | 96.60% | 99.48% |
| *Chlaenius sericeus* JMP070 | - | - | - | 0% | 0% | 0% | 0% |
| *Chlaenius sericeus* JMP071 | 377.09 | 162.36 | 214.72 | 0.21% | 0.21% | 96.57% | 99.58% |
| *Lionepha* "Waterfalls" DNA3782 | 545.72 | 367.11 | 178.61 | 0.06% | 0.06% | 95.81% | 99.49% |
| *Lionepha* "Waterfalls" DNA5435^a^ | 483.39 | 303.42 | 179.97 | 0.17% | 0.17% | 94.90% | 99.32% |
| *Lionepha* "Waterfalls" DNA5436^a^ | 493.63 | 312.77 | 180.86 | 0.20% | 0.20% | 94.77% | 99.35% |
| *Omoglymmius hamatus* DNA3783 | 74.03 | 67.29 | 6.74 | 2.43% | 2.43% | 73.44% | 91.11% |
| *Pterostichus melanarius* DNA3787 | 74.58 | 65.06 | 9.52 | 2.59% | 2.59% | 75.63% | 91.59% |
| *Pterostichus melanarius* JMP059 | - | - | - | 0% | 0% | 0% | 0% |
| *Pterostichus melanarius* JMP060 | - | - | - | 0% | 0% | 0% | 0% |
| *Pterostichus melanarius* JMP061 | - | - | - | 0% | 0% | 0% | 0% |
| *Pterostichus melanarius* JMP062 | - | - | - | 0% | 0% | 0% | 0% |
| *Trachypachus gibbsii* DNA3786 | 264.08 | 81.61 | 182.47 | 0.10% | 0.10% | 97.62% | 99.60% |

Table S8. GenomeScope results. Cells containing dashes indicate that GenomeScope failed to converge. Genome size estimates are given in Mb. Analyses were conducted using a k value of 21. See Vurture et al. 2017 for an explanation of the meaning of these values.

|  |  | **Regier** | | **OrthoDB** | |
| --- | --- | --- | --- | --- | --- |
|  | **Read Count** | **Coverage** | **Genome Size** | **Coverage** | **Genome Size** |
| **Filtered** | | | | | |
| ***Amphizoa insolens* DNA3784** | **53,029,104** | **8.04** | **666.25** | **8.35** | **641.21** |
| *Bembidion haplogonum* DNA2544 | 712,011,300 | 96.18 | 1,117.80 | 93.57 | 1,149.06 |
| *Bembidion haplogonum* DNA5427 | 390,795,034 | 61.13 | 965.30 | 56.65 | 1,041.69 |
| *Bembidion haplogonum* DNA5428 | 222,917,184 | 42.81 | 786.22 | 40.53 | 830.57 |
| *Bembidion haplogonum* DNA5433 | 290,855,186 | 51.43 | 854.01 | 47.93 | 916.33 |
| ***Bembidion lividulum* DNA4161** | **49,060,670** | **8.92** | **555.79** | **8.07** | **614.29** |
| *Chlaenius sericeus* DNA4821 | 75,464,330 | 31.78 | 358.52 | 28.93 | 393.90 |
| *Chlaenius sericeus* JMP068 | 93,341,372 | 38.52 | 365.95 | 40.67 | 346.59 |
| *Chlaenius sericeus* JMP069 | 72,591,474 | 30.31 | 361.69 | 31.64 | 346.47 |
| ***Chlaenius sericeus* JMP070** | **53,617,208** | **24.64** | **328.53** | **22.78** | **355.46** |
| *Chlaenius sericeus* JMP071 | 69,925,928 | 28.77 | 367.05 | 29.19 | 361.74 |
| *Lionepha* "Waterfalls" DNA3782 | 73,477,490 | 9.61 | 772.31 | 11.23 | 661.06 |
| *Lionepha* "Waterfalls" DNA5435^a^ | 110,976,098 | 35.12 | 477.16 | 34.07 | 491.78 |
| *Lionepha* "Waterfalls" DNA5436^a^ | 119,010,276 | 36.22 | 496.14 | 36.19 | 496.59 |
| ***Omoglymmius hamatus* DNA3783** | **76,303,218** | **5.99** | **1,285.63** | **5.92** | **1,300.79** |
| ***Pterostichus melanarius* DNA3787** | **65,498,994** | **6.35** | **1,041.58** | **6.8** | **972.20** |
| ***Pterostichus melanarius* JMP059** | **67,841,732** | **16.01** | **639.89** | **12.63** | **811.14** |
| ***Pterostichus melanarius* JMP060** | **68,305,154** | **13.7** | **753.11** | **13.6** | **758.14** |
| ***Pterostichus melanarius* JMP061** | **86,453,952** | **25.17** | **518.65** | **17.78** | **734.08** |
| ***Pterostichus melanarius* JMP062** | **97,447,446** | **16.29** | **903.50** | **17.19** | **856.00** |
| *Trachypachus gibbsii* DNA3786 | 72,656,132 | 22.83 | 321.46 | 22.81 | 321.76 |
| **No Mito** | | | | | |
| ***Amphizoa insolens* DNA3784** | **51,896,228** | **8.02** | **653.60** | **8.35** | **628.04** |
| *Bembidion haplogonum* DNA2544 | 701,179,000 | 96.18 | 1,100.87 | 93.36 | 1,134.10 |
| *Bembidion haplogonum* DNA5427 | 385,572,878 | 61.15 | 952.11 | 56.72 | 1,026.45 |
| *Bembidion haplogonum* DNA5428 | 218,017,252 | 42.83 | 768.69 | 40.55 | 811.80 |
| *Bembidion haplogonum* DNA5433 | 286,877,384 | 51.43 | 842.24 | 47.95 | 903.33 |
| ***Bembidion lividulum* DNA4161** | **48,990,136** | **8.92** | **554.99** | **8.07** | **613.41** |
| *Chlaenius sericeus* DNA4821 | 74,877,844 | 31.81 | 355.48 | 28.93 | 390.84 |
| *Chlaenius sericeus* JMP068 | 92,333,434 | 38.69 | 360.32 | 40.73 | 342.32 |
| *Chlaenius sericeus* JMP069 | 72,013,564 | 30.39 | 357.78 | 31.7 | 343.02 |
| ***Chlaenius sericeus* JMP070** | **53,119,248** | **24.69** | **324.89** | **22.82** | **351.43** |
| *Chlaenius sericeus* JMP071 | 69,184,928 | 28.88 | 361.80 | 29.11 | 358.92 |
| *Lionepha* "Waterfalls" DNA3782 | 73,040,060 | 9.61 | 767.71 | 11.23 | 657.13 |
| *Lionepha* "Waterfalls" DNA5435^a^ | 110,544,262 | 35.26 | 473.40 | 34.16 | 488.68 |
| *Lionepha* "Waterfalls" DNA5436^a^ | 115,669,748 | 36.51 | 478.34 | 36.19 | 482.67 |
| ***Omoglymmius hamatus* DNA3783** | **75,837,778** | **5.99** | **1,277.79** | **5.91** | **1,295.26** |
| ***Pterostichus melanarius* DNA3787** | **64,380,826** | **6.35** | **1,023.80** | **6.8** | **955.60** |
| ***Pterostichus melanarius* JMP059** | **67,306,770** | **16.06** | **632.64** | **12.66** | **802.69** |
| ***Pterostichus melanarius* JMP060** | **67,908,474** | **13.72** | **747.12** | **13.64** | **752.00** |
| ***Pterostichus melanarius* JMP061** | **86,110,430** | **25.23** | **515.26** | **17.81** | **730.23** |
| ***Pterostichus melanarius* JMP062** | **96,703,360** | **16.41** | **889.86** | **17.23** | **847.71** |
| *Trachypachus gibbsii* DNA3786 | 71,720,174 | 22.81 | 317.63 | 22.8 | 317.71 |

Table S9. Summary of read mapping genome size estimates for the Regier and OrthoDB gene sets using three different read filtering methods. Genome size is calculated by multiplying the read length by total read, then dividing by coverage. Genome sizes are given in Mb.

a. Samples made with DNA extracted from different tissues of the same individual.

|  | **Basic** | | **Repeat** | |
| --- | --- | --- | --- | --- |
|  | **Genome Size** | **Coverage** | **Genome Size** | **Coverage** |
| ***Amphizoa insolens* DNA3784** | **376.78** | **13.9** | **728.47** | **7.19** |
| *Bembidion haplogonum* DNA2544 | 932.04 | 113.49 | 2,140.04 | 49.43 |
| *Bembidion haplogonum* DNA5427 | 827.88 | 70.28 | 1,980.24 | 29.38 |
| *Bembidion haplogonum* DNA5428 | 813.44 | 40.44 | 1,924.27 | 17.1 |
| *Bembidion haplogonum* DNA5433 | 758.09 | 57.1 | 1,480.15 | 29.25 |
| ***Bembidion lividulum* DNA4161** | **359.94** | **13.77** | **790.81** | **6.27** |
| *Chlaenius sericeus* DNA4821 | 386.92 | 29.14 | 608.44 | 18.53 |
| *Chlaenius sericeus* JMP068 | 410.91 | 33.89 | 751.89 | 18.52 |
| *Chlaenius sericeus* JMP069 | 395.86 | 27.46 | 704.51 | 15.43 |
| ***Chlaenius sericeus* JMP070** | **409.81** | **19.57** | **796.39** | **10.07** |
| *Chlaenius sericeus* JMP071 | 412.78 | 25.3 | 624.53 | 16.72 |
| *Lionepha* "Waterfalls" DNA3782 | 442.32 | 16.66 | 659.97 | 11.17 |
| *Lionepha* "Waterfalls" DNA5435^a^ | 340.38 | 49 | 423.66 | 39.37 |
| *Lionepha* "Waterfalls" DNA5436^a^ | 346.52 | 50.39 | 578.96 | 30.16 |
| ***Omoglymmius hamatus* DNA3783** | **627.13** | **12.2** | **1,188.94** | **6.44** |
| ***Pterostichus melanarius* DNA3787** | **475.11** | **13.67** | **1,221.83** | **5.32** |
| ***Pterostichus melanarius* JMP059** | **586.67** | **17.32** | **1,127.86** | **9.01** |
| ***Pterostichus melanarius* JMP060** | **597.29** | **17.16** | **1,145.04** | **8.95** |
| ***Pterostichus melanarius* JMP061** | **650.22** | **19.99** | **1,454.54** | **8.94** |
| ***Pterostichus melanarius* JMP062** | **678.61** | **21.51** | **1,181.38** | **12.36** |
| *Trachypachus gibbsii* DNA3786 | 314.81 | 22.99 | 525.13 | 13.78 |

Table S10. CovEST genome size (in Mb) and coverage estimates for two models, Basic and Repeat, performed using a k value of 21.

a. Samples made with DNA extracted from different tissues of the same individual.

|  | **Regier Read Mapping Coverages** | | |
| --- | --- | --- | --- |
|  | **Untrimmed** | **IQR Trim** | **Difference** |
| *Amphizoa insolens* DNA3784 | 8.02 | 8.02 | -0.00056 |
| *Bembidion haplogonum* DNA2544 | 96.18 | 96.18 | -0.00371 |
| *Bembidion lividulum* DNA4161 | 8.92 | 8.92 | -0.00449 |
| *Chlaenius sericeus* DNA4821 | 31.81 | 31.81 | -0.00367 |
| *Lionepha* "Waterfalls" DNA3782 | 9.61 | 9.61 | -0.00082 |
| *Omoglymmius hamatus* DNA3783 | 5.99 | 11.64 | 5.65421 |
| *Pterostichus melanarius* DNA3787 | 6.35 | 6.35 | 0.00129 |
| *Trachypachus gibbsii* DNA3786 | 22.81 | 22.81 | -0.00433 |

Table S11. Regier read mapping mean coverages before and after removing outliers using the 3*IQR rule. The columns “Untrimmed” and “IQR Trim” indicate the mean coverage of each sample before and after, respecitively, removing outlier. The column “Difference” is the difference in coverage between “IQR Trim” and “Untrimmed.”

|  | **ODB Read Mapping Coverages** | | |
| --- | --- | --- | --- |
|  | **Untrimmed** | **IQR Trim** | **Difference** |
| *Amphizoa insolens* DNA3784 | 8.35 | 8.09 | -0.259 |
| *Bembidion haplogonum* DNA2544 | 93.36 | 87.99 | -5.374 |
| *Bembidion lividulum* DNA4161 | 8.07 | 7.82 | -0.254 |
| *Chlaenius sericeus* DNA4821 | 28.93 | 25.82 | -3.111 |
| *Lionepha* "Waterfalls" DNA3782 | 11.23 | 10.93 | -0.299 |
| *Omoglymmius hamatus* DNA3783 | 5.91 | 5.12 | -0.794 |
| *Pterostichus melanarius* DNA3787 | 6.8 | 6.61 | -0.190 |
| *Trachypachus gibbsii* DNA3786 | 22.8 | 22.80 | 0.000 |

Table S12. OrthoDB read mapping mean coverages before (“Untrimmed”) and after (“IQR Trim”) removing outliers more than three interquartiles from the median. The column “Difference” is the difference between “IQR Trim” and “Untrimmed.”

|  | RepeatExplorer (Mb) | GenomeScope (Mb) |
| --- | --- | --- |
| *Bembidion haplogonum* DNA2544 | 1,410.4 | 889.0 |
| *Chlaenius sericeus* DNA4821 | 139.6 | 185.9 |
| *Lionepha* "Waterfalls" DNA3782 | 329.7 | 367.1 |

Table S13. Comparison of total repetitive DNA sequence (in Mb) estimated by RepeatExplorer and GenomeScope. Values for RepeatExplorer were extrapolated by multiplying the total fraction of repeats by the genome size estimated with flow cytometry.

| **Accession** | **Strain** | **Tissue** | **GenomeScope estimate (Mb)** |
| --- | --- | --- | --- |
| SRR1515985 | y ; cn bw sp | embryo | 202,210,977 |
| SRR1516222 | y ; cn bw sp | embryo | 163,268,455 |
| SRR1516224 | y ; cn bw sp | embryo | 176,703,147 |
| SRR1519053 | In(1)scV2, scV2; SuURES | 0.5-2.5hr embryo | 198,516,951 |
| SRR1516221 | y ; cn bw sp | salivary gland | 117,956,035 |
| SRR1516223 | y ; cn bw sp | salivary gland | 118,452,887 |
| SRR1516225 | y ; cn bw sp | salivary gland | 118,396,656 |
| SRR1516226 | y w | salivary gland | 89,033,495 |
| SRR1516227 | y ; ry[506] | salivary gland | 129,031,716 |
| SRR1516228 | y ; cn bw sp | salivary gland | 120,086,988 |
| SRR1517773 | SUUR -/- | salivary gland | 129,738,053 |
| SRR1518366 | y ; ry[506] | salivary gland | 128,983,897 |
| SRR1519051 | OreR P2 | ovarian follicle | 141,919,753 |
| SRR1519054 | y ; cn bw sp | whole ovary | 140,138,513 |

Table S14. GenomeScope genome size estimates using *Drosophila melanogaster* embryo, salivary gland, and ovarian cell data set from Yarosh and Spradling (2014).
