## Supplemental File 1 for "Measuring genome sizes using read-depth, k-mers, and flow cytometry: methodological comparisons in beetles (Coleoptera)"

**Oneway Analysis of CovEST Basic By K**
Species=*Bembidion haplogonum*

| Rsquare | 0.916335 |
| --- | --- |
| Adj Rsquare | 0.891236 |
| Root Mean Square Error | 113.4786 |
| Mean of Response | 644.8326 |
| Observations (or Sum Wgts) | 40 |

**Analysis of Variance**

| **Source** | **DF** | **Sum of Squares** | **Mean Square** | **F Ratio** | **Prob > F** |
| --- | --- | --- | --- | --- | --- |
| K | 9 | 4231162.1 | 470129 | 36.5081 | <.0001* |
| Error | 30 | 386321.6 | 12877 |  |  |
| C. Total | 39 | 4617483.7 |  |  |  |

**Means for Oneway Anova**

| **Level** | **Number** | **Mean** | **Std Error** | **Lower 95%** | **Upper 95%** |
| --- | --- | --- | --- | --- | --- |
| 13 | 4 | 33.095 | 56.739 | -82.8 | 149.0 |
| 15 | 4 | 4.900 | 56.739 | -111.0 | 120.8 |
| 17 | 4 | 561.860 | 56.739 | 446.0 | 677.7 |
| 19 | 4 | 742.308 | 56.739 | 626.4 | 858.2 |
| 21 | 4 | 832.864 | 56.739 | 717.0 | 948.7 |
| 23 | 4 | 843.354 | 56.739 | 727.5 | 959.2 |
| 25 | 4 | 836.728 | 56.739 | 720.9 | 952.6 |
| 27 | 4 | 874.741 | 56.739 | 758.9 | 990.6 |
| 29 | 4 | 884.546 | 56.739 | 768.7 | 1000.4 |
| 31 | 4 | 833.929 | 56.739 | 718.1 | 949.8 |

Std Error uses a pooled estimate of error variance

**Means Comparisons**

**Comparisons for all pairs using Tukey-Kramer HSD**

**Confidence Quantile**

| **q*** | **Alpha** |
| --- | --- |
| 3.41119 | 0.05 |

**Connecting Letters Report**

| **Level** |  |  |  | **Mean** |
| --- | --- | --- | --- | --- |
| 29 | A |  |  | 884.54570 |
| 27 | A |  |  | 874.74095 |
| 23 | A |  |  | 843.35402 |
| 25 | A |  |  | 836.72841 |
| 31 | A | B |  | 833.92916 |
| 21 | A | B |  | 832.86432 |
| 19 | A | B |  | 742.30809 |
| 17 |  | B |  | 561.86022 |
| 13 |  |  | C | 33.09547 |
| 15 |  |  | C | 4.89986 |

Levels not connected by same letter are significantly different.

**Oneway Analysis of CovEST Basic By K**

Species=*Chlaenius sericeus*

**Oneway Anova**

**Summary of Fit**

| Rsquare | 0.93213 |
| --- | --- |
| Adj Rsquare | 0.916859 |
| Root Mean Square Error | 42.31397 |
| Mean of Response | 327.2613 |
| Observations (or Sum Wgts) | 50 |

**Analysis of Variance**

| **Source** | **DF** | **Sum of Squares** | **Mean Square** | **F Ratio** | **Prob > F** |
| --- | --- | --- | --- | --- | --- |
| K | 9 | 983619.1 | 109291 | 61.0403 | <.0001* |
| Error | 40 | 71618.9 | 1790 |  |  |
| C. Total | 49 | 1055238.0 |  |  |  |

**Means for Oneway Anova**

| **Level** | **Number** | **Mean** | **Std Error** | **Lower 95%** | **Upper 95%** |
| --- | --- | --- | --- | --- | --- |
| 13 | 5 | 0.479 | 18.923 | -37.8 | 38.72 |
| 15 | 5 | 130.901 | 18.923 | 92.7 | 169.15 |
| 17 | 5 | 282.101 | 18.923 | 243.9 | 320.35 |
| 19 | 5 | 373.827 | 18.923 | 335.6 | 412.07 |
| 21 | 5 | 403.257 | 18.923 | 365.0 | 441.50 |
| 23 | 5 | 412.963 | 18.923 | 374.7 | 451.21 |
| 25 | 5 | 423.580 | 18.923 | 385.3 | 461.83 |
| 27 | 5 | 428.977 | 18.923 | 390.7 | 467.22 |
| 29 | 5 | 433.815 | 18.923 | 395.6 | 472.06 |
| 31 | 5 | 382.714 | 18.923 | 344.5 | 420.96 |

Std Error uses a pooled estimate of error variance

**Means Comparisons**

**Comparisons for all pairs using Tukey-Kramer HSD**

**Confidence Quantile**

| **q*** | **Alpha** |
| --- | --- |
| 3.34782 | 0.05 |

**Connecting Letters Report**

| **Level** |  |  |  |  | **Mean** |
| --- | --- | --- | --- | --- | --- |
| 29 | A |  |  |  | 433.81456 |
| 27 | A |  |  |  | 428.97692 |
| 25 | A |  |  |  | 423.57962 |
| 23 | A |  |  |  | 412.96310 |
| 21 | A |  |  |  | 403.25661 |
| 31 | A |  |  |  | 382.71439 |
| 19 | A |  |  |  | 373.82709 |
| 17 |  | B |  |  | 282.10082 |
| 15 |  |  | C |  | 130.90131 |
| 13 |  |  |  | D | 0.47894 |

Levels not connected by same letter are significantly different.

**Oneway Analysis of CovEST Basic By K**

Species=*Lionepha* "Waterfalls"

**Oneway Anova**

**Summary of Fit**

| Rsquare | 0.914478 |
| --- | --- |
| Adj Rsquare | 0.875993 |
| Root Mean Square Error | 45.05389 |
| Mean of Response | 316.1463 |
| Observations (or Sum Wgts) | 30 |

**Analysis of Variance**

| **Source** | **DF** | **Sum of Squares** | **Mean Square** | **F Ratio** | **Prob > F** |
| --- | --- | --- | --- | --- | --- |
| K | 9 | 434099.22 | 48233.2 | 23.7619 | <.0001* |
| Error | 20 | 40597.07 | 2029.9 |  |  |
| C. Total | 29 | 474696.29 |  |  |  |

**Means for Oneway Anova**

| **Level** | **Number** | **Mean** | **Std Error** | **Lower 95%** | **Upper 95%** |
| --- | --- | --- | --- | --- | --- |
| 13 | 3 | 5.723 | 26.012 | -48.5 | 59.98 |
| 15 | 3 | 185.812 | 26.012 | 131.6 | 240.07 |
| 17 | 3 | 300.134 | 26.012 | 245.9 | 354.39 |
| 19 | 3 | 369.705 | 26.012 | 315.4 | 423.97 |
| 21 | 3 | 376.406 | 26.012 | 322.1 | 430.67 |
| 23 | 3 | 399.519 | 26.012 | 345.3 | 453.78 |
| 25 | 3 | 396.000 | 26.012 | 341.7 | 450.26 |
| 27 | 3 | 370.583 | 26.012 | 316.3 | 424.84 |
| 29 | 3 | 393.859 | 26.012 | 339.6 | 448.12 |
| 31 | 3 | 363.723 | 26.012 | 309.5 | 417.98 |

Std Error uses a pooled estimate of error variance

**Means Comparisons**

**Comparisons for all pairs using Tukey-Kramer HSD**

**Confidence Quantile**

| **q*** | **Alpha** |
| --- | --- |
| 3.54111 | 0.05 |

**Connecting Letters Report**

| **Level** |  |  |  | **Mean** |
| --- | --- | --- | --- | --- |
| 23 | A |  |  | 399.51883 |
| 25 | A |  |  | 396.00013 |
| 29 | A |  |  | 393.85862 |
| 21 | A |  |  | 376.40624 |
| 27 | A |  |  | 370.58297 |
| 19 | A |  |  | 369.70529 |
| 31 | A |  |  | 363.72257 |
| 17 | A | B |  | 300.13385 |
| 15 |  | B |  | 185.81194 |
| 13 |  |  | C | 5.72271 |

Levels not connected by same letter are significantly different.

**Oneway Analysis of CovEST Basic By K**

Species=*Pterostichus melanarius*

**Oneway Anova**

**Summary of Fit**

| Rsquare | 0.906674 |
| --- | --- |
| Adj Rsquare | 0.885676 |
| Root Mean Square Error | 78.73282 |
| Mean of Response | 484.9251 |
| Observations (or Sum Wgts) | 50 |

**Analysis of Variance**

| **Source** | **DF** | **Sum of Squares** | **Mean Square** | **F Ratio** | **Prob > F** |
| --- | --- | --- | --- | --- | --- |
| K | 9 | 2408918.6 | 267658 | 43.1785 | <.0001* |
| Error | 40 | 247954.3 | 6199 |  |  |
| C. Total | 49 | 2656872.8 |  |  |  |

**Means for Oneway Anova**

| **Level** | **Number** | **Mean** | **Std Error** | **Lower 95%** | **Upper 95%** |
| --- | --- | --- | --- | --- | --- |
| 13 | 5 | 3.961 | 35.210 | -67.2 | 75.12 |
| 15 | 5 | 132.038 | 35.210 | 60.9 | 203.20 |
| 17 | 5 | 416.042 | 35.210 | 344.9 | 487.21 |
| 19 | 5 | 558.611 | 35.210 | 487.4 | 629.77 |
| 21 | 5 | 597.579 | 35.210 | 526.4 | 668.74 |
| 23 | 5 | 613.747 | 35.210 | 542.6 | 684.91 |
| 25 | 5 | 622.378 | 35.210 | 551.2 | 693.54 |
| 27 | 5 | 630.261 | 35.210 | 559.1 | 701.42 |
| 29 | 5 | 635.585 | 35.210 | 564.4 | 706.75 |
| 31 | 5 | 639.046 | 35.210 | 567.9 | 710.21 |

Std Error uses a pooled estimate of error variance

**Means Comparisons**

**Comparisons for all pairs using Tukey-Kramer HSD**

**Confidence Quantile**

| **q*** | **Alpha** |
| --- | --- |
| 3.34782 | 0.05 |

**Connecting Letters Report**

| **Level** |  |  |  | **Mean** |
| --- | --- | --- | --- | --- |
| 31 | A |  |  | 639.04645 |
| 29 | A |  |  | 635.58540 |
| 27 | A |  |  | 630.26121 |
| 25 | A |  |  | 622.37847 |
| 23 | A |  |  | 613.74679 |
| 21 | A |  |  | 597.57938 |
| 19 | A | B |  | 558.61096 |
| 17 |  | B |  | 416.04243 |
| 15 |  |  | C | 132.03818 |
| 13 |  |  | C | 3.96131 |

Levels not connected by same letter are significantly different.

**Oneway Analysis of CovEST Repeat By K**

Species=*Bembidion haplogonum*

**Oneway Anova**

**Summary of Fit**

| Rsquare | 0.676426 |
| --- | --- |
| Adj Rsquare | 0.579354 |
| Root Mean Square Error | 452.4897 |
| Mean of Response | 1329.541 |
| Observations (or Sum Wgts) | 40 |

**Analysis of Variance**

| **Source** | **DF** | **Sum of Squares** | **Mean Square** | **F Ratio** | **Prob > F** |
| --- | --- | --- | --- | --- | --- |
| K | 9 | 12840623 | 1426736 | 6.9683 | <.0001* |
| Error | 30 | 6142408 | 204747 |  |  |
| C. Total | 39 | 18983031 |  |  |  |

**Means for Oneway Anova**

| **Level** | **Number** | **Mean** | **Std Error** | **Lower 95%** | **Upper 95%** |
| --- | --- | --- | --- | --- | --- |
| 13 | 4 | 62.08 | 226.24 | -400 | 524.1 |
| 15 | 4 | 514.92 | 226.24 | 53 | 977.0 |
| 17 | 4 | 1360.34 | 226.24 | 898 | 1822.4 |
| 19 | 4 | 1233.61 | 226.24 | 772 | 1695.7 |
| 21 | 4 | 1881.17 | 226.24 | 1419 | 2343.2 |
| 23 | 4 | 1662.90 | 226.24 | 1201 | 2125.0 |
| 25 | 4 | 1711.01 | 226.24 | 1249 | 2173.1 |
| 27 | 4 | 1868.59 | 226.24 | 1407 | 2330.6 |
| 29 | 4 | 1600.47 | 226.24 | 1138 | 2062.5 |
| 31 | 4 | 1400.32 | 226.24 | 938 | 1862.4 |

Std Error uses a pooled estimate of error variance

**Means Comparisons**

**Comparisons for all pairs using Tukey-Kramer HSD**

**Confidence Quantile**

| **q*** | **Alpha** |
| --- | --- |
| 3.41119 | 0.05 |

**Connecting Letters Report**

| **Level** |  |  |  | **Mean** |
| --- | --- | --- | --- | --- |
| 21 | A |  |  | 1881.1733 |
| 27 | A |  |  | 1868.5914 |
| 25 | A |  |  | 1711.0059 |
| 23 | A |  |  | 1662.8997 |
| 29 | A | B |  | 1600.4694 |
| 31 | A | B |  | 1400.3243 |
| 17 | A | B |  | 1360.3423 |
| 19 | A | B |  | 1233.6055 |
| 15 |  | B | C | 514.9156 |
| 13 |  |  | C | 62.0792 |

Levels not connected by same letter are significantly different.

**Oneway Analysis of CovEST Repeat By K**

Species=*Chlaenius sericeus*

**Oneway Anova**

**Summary of Fit**

| Rsquare | 0.831119 |
| --- | --- |
| Adj Rsquare | 0.79312 |
| Root Mean Square Error | 123.0309 |
| Mean of Response | 590.916 |
| Observations (or Sum Wgts) | 50 |

**Analysis of Variance**

| **Source** | **DF** | **Sum of Squares** | **Mean Square** | **F Ratio** | **Prob > F** |
| --- | --- | --- | --- | --- | --- |
| K | 9 | 2979682.0 | 331076 | 21.8725 | <.0001* |
| Error | 40 | 605464.2 | 15137 |  |  |
| C. Total | 49 | 3585146.2 |  |  |  |

**Means for Oneway Anova**

| **Level** | **Number** | **Mean** | **Std Error** | **Lower 95%** | **Upper 95%** |
| --- | --- | --- | --- | --- | --- |
| 13 | 5 | 59.722 | 55.021 | -51.5 | 170.92 |
| 15 | 5 | 176.312 | 55.021 | 65.1 | 287.51 |
| 17 | 5 | 581.408 | 55.021 | 470.2 | 692.61 |
| 19 | 5 | 662.963 | 55.021 | 551.8 | 774.16 |
| 21 | 5 | 697.152 | 55.021 | 586.0 | 808.35 |
| 23 | 5 | 733.210 | 55.021 | 622.0 | 844.41 |
| 25 | 5 | 764.071 | 55.021 | 652.9 | 875.27 |
| 27 | 5 | 757.230 | 55.021 | 646.0 | 868.43 |
| 29 | 5 | 782.285 | 55.021 | 671.1 | 893.49 |
| 31 | 5 | 694.806 | 55.021 | 583.6 | 806.01 |

Std Error uses a pooled estimate of error variance

**Means Comparisons**

**Comparisons for all pairs using Tukey-Kramer HSD**

**Confidence Quantile**

| **q*** | **Alpha** |
| --- | --- |
| 3.34782 | 0.05 |

**Connecting Letters Report**

| **Level** |  |  | **Mean** |
| --- | --- | --- | --- |
| 29 | A |  | 782.28481 |
| 25 | A |  | 764.07140 |
| 27 | A |  | 757.22959 |
| 23 | A |  | 733.20980 |
| 21 | A |  | 697.15219 |
| 31 | A |  | 694.80622 |
| 19 | A |  | 662.96304 |
| 17 | A |  | 581.40837 |
| 15 |  | B | 176.31205 |
| 13 |  | B | 59.72212 |

Levels not connected by same letter are significantly different.

**Oneway Analysis of CovEST Repeat By K**

Species=*Lionepha* "Waterfalls"

**Oneway Anova**

**Summary of Fit**

| Rsquare | 0.810666 |
| --- | --- |
| Adj Rsquare | 0.725466 |
| Root Mean Square Error | 117.5257 |
| Mean of Response | 535.4586 |
| Observations (or Sum Wgts) | 30 |

**Analysis of Variance**

| **Source** | **DF** | **Sum of Squares** | **Mean Square** | **F Ratio** | **Prob > F** |
| --- | --- | --- | --- | --- | --- |
| K | 9 | 1182794.4 | 131422 | 9.5148 | <.0001* |
| Error | 20 | 276246.0 | 13812 |  |  |
| C. Total | 29 | 1459040.4 |  |  |  |

**Means for Oneway Anova**

| **Level** | **Number** | **Mean** | **Std Error** | **Lower 95%** | **Upper 95%** |
| --- | --- | --- | --- | --- | --- |
| 13 | 3 | 88.438 | 67.854 | -53.1 | 229.98 |
| 15 | 3 | 238.161 | 67.854 | 96.6 | 379.70 |
| 17 | 3 | 538.593 | 67.854 | 397.1 | 680.13 |
| 19 | 3 | 561.062 | 67.854 | 419.5 | 702.60 |
| 21 | 3 | 554.198 | 67.854 | 412.7 | 695.74 |
| 23 | 3 | 669.559 | 67.854 | 528.0 | 811.10 |
| 25 | 3 | 645.617 | 67.854 | 504.1 | 787.16 |
| 27 | 3 | 748.635 | 67.854 | 607.1 | 890.17 |
| 29 | 3 | 634.965 | 67.854 | 493.4 | 776.50 |
| 31 | 3 | 675.358 | 67.854 | 533.8 | 816.90 |

Std Error uses a pooled estimate of error variance

**Means Comparisons**

**Comparisons for all pairs using Tukey-Kramer HSD**

**Confidence Quantile**

| **q*** | **Alpha** |
| --- | --- |
| 3.54111 | 0.05 |

**Connecting Letters Report**

| **Level** |  |  |  | **Mean** |
| --- | --- | --- | --- | --- |
| 27 | A |  |  | 748.63499 |
| 31 | A |  |  | 675.35769 |
| 23 | A |  |  | 669.55908 |
| 25 | A |  |  | 645.61669 |
| 29 | A |  |  | 634.96493 |
| 19 | A | B |  | 561.06193 |
| 21 | A | B |  | 554.19803 |
| 17 | A | B |  | 538.59306 |
| 15 |  | B | C | 238.16084 |
| 13 |  |  | C | 88.43830 |

Levels not connected by same letter are significantly different.

**Oneway Analysis of CovEST Repeat By K**

Species=*Pterostichus melanarius*

**Oneway Anova**

**Summary of Fit**

| Rsquare | 0.831823 |
| --- | --- |
| Adj Rsquare | 0.793983 |
| Root Mean Square Error | 195.7973 |
| Mean of Response | 1010.004 |
| Observations (or Sum Wgts) | 50 |

**Analysis of Variance**

| **Source** | **DF** | **Sum of Squares** | **Mean Square** | **F Ratio** | **Prob > F** |
| --- | --- | --- | --- | --- | --- |
| K | 9 | 7584673.6 | 842742 | 21.9827 | <.0001* |
| Error | 40 | 1533463.2 | 38337 |  |  |
| C. Total | 49 | 9118136.9 |  |  |  |

**Means for Oneway Anova**

| **Level** | **Number** | **Mean** | **Std Error** | **Lower 95%** | **Upper 95%** |
| --- | --- | --- | --- | --- | --- |
| 13 | 5 | 69.82 | 87.563 | -107 | 246.8 |
| 15 | 5 | 458.43 | 87.563 | 281 | 635.4 |
| 17 | 5 | 1202.74 | 87.563 | 1026 | 1379.7 |
| 19 | 5 | 1129.69 | 87.563 | 953 | 1306.7 |
| 21 | 5 | 1226.13 | 87.563 | 1049 | 1403.1 |
| 23 | 5 | 1229.29 | 87.563 | 1052 | 1406.3 |
| 25 | 5 | 1025.84 | 87.563 | 849 | 1202.8 |
| 27 | 5 | 1193.92 | 87.563 | 1017 | 1370.9 |
| 29 | 5 | 1268.99 | 87.563 | 1092 | 1446.0 |
| 31 | 5 | 1295.19 | 87.563 | 1118 | 1472.2 |

Std Error uses a pooled estimate of error variance

**Means Comparisons**

**Comparisons for all pairs using Tukey-Kramer HSD**

**Confidence Quantile**

| **q*** | **Alpha** |
| --- | --- |
| 3.34782 | 0.05 |

**Connecting Letters Report**

| **Level** |  |  | **Mean** |
| --- | --- | --- | --- |
| 31 | A |  | 1295.1905 |
| 29 | A |  | 1268.9864 |
| 23 | A |  | 1229.2949 |
| 21 | A |  | 1226.1304 |
| 17 | A |  | 1202.7370 |
| 27 | A |  | 1193.9225 |
| 19 | A |  | 1129.6911 |
| 25 | A |  | 1025.8411 |
| 15 |  | B | 458.4302 |
| 13 |  | B | 69.8174 |

Levels not connected by same letter are significantly different.

**Oneway Analysis of GenomeScope By K**

Species=*Bembidion haplogonum*

**Oneway Anova**

**Summary of Fit**

| Rsquare | 0.034083 |
| --- | --- |
| Adj Rsquare | -0.26312 |
| Root Mean Square Error | 96.98382 |
| Mean of Response | 985.8962 |
| Observations (or Sum Wgts) | 35 |

**Analysis of Variance**

| **Source** | **DF** | **Sum of Squares** | **Mean Square** | **F Ratio** | **Prob > F** |
| --- | --- | --- | --- | --- | --- |
| K | 8 | 8629.26 | 1078.66 | 0.1147 | 0.9982 |
| Error | 26 | 244552.41 | 9405.86 |  |  |
| C. Total | 34 | 253181.67 |  |  |  |

**Means for Oneway Anova**

| **Level** | **Number** | **Mean** | **Std Error** | **Lower 95%** | **Upper 95%** |
| --- | --- | --- | --- | --- | --- |
| 15 | 3 | 939.111 | 55.994 | 824.01 | 1054.2 |
| 17 | 4 | 972.868 | 48.492 | 873.19 | 1072.5 |
| 19 | 4 | 990.701 | 48.492 | 891.02 | 1090.4 |
| 21 | 4 | 994.562 | 48.492 | 894.89 | 1094.2 |
| 23 | 4 | 994.112 | 48.492 | 894.44 | 1093.8 |
| 25 | 4 | 993.331 | 48.492 | 893.65 | 1093.0 |
| 27 | 4 | 993.573 | 48.492 | 893.90 | 1093.2 |
| 29 | 4 | 992.551 | 48.492 | 892.87 | 1092.2 |
| 31 | 4 | 990.561 | 48.492 | 890.88 | 1090.2 |

Std Error uses a pooled estimate of error variance

**Means Comparisons**

**Comparisons for all pairs using Tukey-Kramer HSD**

**Confidence Quantile**

| **q*** | **Alpha** |
| --- | --- |
| 3.37522 | 0.05 |

**Connecting Letters Report**

| **Level** |  | **Mean** |
| --- | --- | --- |
| 21 | A | 994.56166 |
| 23 | A | 994.11203 |
| 27 | A | 993.57276 |
| 25 | A | 993.33086 |
| 29 | A | 992.55129 |
| 19 | A | 990.70146 |
| 31 | A | 990.56058 |
| 17 | A | 972.86785 |
| 15 | A | 939.11094 |

Levels not connected by same letter are significantly different.

**Oneway Analysis of GenomeScope By K**

Species=*Chlaenius sericeus*

**Oneway Anova**

**Summary of Fit**

| Rsquare | 0.055785 |
| --- | --- |
| Adj Rsquare | -0.22398 |
| Root Mean Square Error | 20.3626 |
| Mean of Response | 381.1042 |
| Observations (or Sum Wgts) | 36 |

**Analysis of Variance**

| **Source** | **DF** | **Sum of Squares** | **Mean Square** | **F Ratio** | **Prob > F** |
| --- | --- | --- | --- | --- | --- |
| K | 8 | 661.417 | 82.677 | 0.1994 | 0.9885 |
| Error | 27 | 11195.162 | 414.636 |  |  |
| C. Total | 35 | 11856.579 |  |  |  |

**Means for Oneway Anova**

| **Level** | **Number** | **Mean** | **Std Error** | **Lower 95%** | **Upper 95%** |
| --- | --- | --- | --- | --- | --- |
| 15 | 2 | 368.841 | 14.399 | 339.30 | 398.38 |
| 17 | 5 | 375.400 | 9.106 | 356.72 | 394.09 |
| 19 | 5 | 379.565 | 9.106 | 360.88 | 398.25 |
| 21 | 4 | 385.171 | 10.181 | 364.28 | 406.06 |
| 23 | 4 | 384.852 | 10.181 | 363.96 | 405.74 |
| 25 | 4 | 384.031 | 10.181 | 363.14 | 404.92 |
| 27 | 4 | 383.226 | 10.181 | 362.34 | 404.12 |
| 29 | 4 | 382.558 | 10.181 | 361.67 | 403.45 |
| 31 | 4 | 381.972 | 10.181 | 361.08 | 402.86 |

Std Error uses a pooled estimate of error variance

**Means Comparisons**

**Comparisons for all pairs using Tukey-Kramer HSD**

**Confidence Quantile**

| **q*** | **Alpha** |
| --- | --- |
| 3.36470 | 0.05 |

**Connecting Letters Report**

| **Level** |  | **Mean** |
| --- | --- | --- |
| 21 | A | 385.17138 |
| 23 | A | 384.85221 |
| 25 | A | 384.03080 |
| 27 | A | 383.22614 |
| 29 | A | 382.55813 |
| 31 | A | 381.97237 |
| 19 | A | 379.56509 |
| 17 | A | 375.40019 |
| 15 | A | 368.84055 |

Levels not connected by same letter are significantly different.

**Oneway Analysis of GenomeScope By K**

Species=*Lionepha "Waterfalls"*

**Oneway Anova**

**Summary of Fit**

| Rsquare | 0.411221 |
| --- | --- |
| Adj Rsquare | 0.048895 |
| Root Mean Square Error | 54.51033 |
| Mean of Response | 482.0498 |
| Observations (or Sum Wgts) | 22 |

**Analysis of Variance**

| **Source** | **DF** | **Sum of Squares** | **Mean Square** | **F Ratio** | **Prob > F** |
| --- | --- | --- | --- | --- | --- |
| K | 8 | 26978.828 | 3372.35 | 1.1349 | 0.4027 |
| Error | 13 | 38627.885 | 2971.38 |  |  |
| C. Total | 21 | 65606.714 |  |  |  |

**Means for Oneway Anova**

| **Level** | **Number** | **Mean** | **Std Error** | **Lower 95%** | **Upper 95%** |
| --- | --- | --- | --- | --- | --- |
| 15 | 3 | 398.760 | 31.472 | 330.77 | 466.75 |
| 17 | 3 | 508.487 | 31.472 | 440.50 | 576.48 |
| 19 | 3 | 508.061 | 31.472 | 440.07 | 576.05 |
| 21 | 3 | 507.582 | 31.472 | 439.59 | 575.57 |
| 23 | 2 | 487.200 | 38.545 | 403.93 | 570.47 |
| 25 | 2 | 485.336 | 38.545 | 402.07 | 568.61 |
| 27 | 2 | 483.521 | 38.545 | 400.25 | 566.79 |
| 29 | 2 | 481.853 | 38.545 | 398.58 | 565.12 |
| 31 | 2 | 480.304 | 38.545 | 397.03 | 563.57 |

Std Error uses a pooled estimate of error variance

**Means Comparisons**

**Comparisons for all pairs using Tukey-Kramer HSD**

**Confidence Quantile**

| **q*** | **Alpha** |
| --- | --- |
| 3.67130 | 0.05 |

**Connecting Letters Report**

| **Level** |  | **Mean** |
| --- | --- | --- |
| 17 | A | 508.48673 |
| 19 | A | 508.06074 |
| 21 | A | 507.58162 |
| 23 | A | 487.20017 |
| 25 | A | 485.33644 |
| 27 | A | 483.52060 |
| 29 | A | 481.85294 |
| 31 | A | 480.30428 |
| 15 | A | 398.75992 |

Levels not connected by same\0 letter are significantly different.
