## Supplemental File 6 for "Measuring genome sizes using read-depth, k-mers, and flow cytometry: methodological comparisons in beetles (Coleoptera)": gsec.py

```
#!/usr/bin/env python

import argparse
import os, sys
import numpy as np
import collections

parser = argparse.ArgumentParser(description="Trims and estimates genome sequencing coverage from a base coverage file.", 
    formatter_class=argparse.ArgumentDefaultsHelpFormatter)

parser.add_argument("-basecov", "-i",
                    help="Base coverage file. Should be a tab delimited file with a sequence name, position number, and coverage.",
                    required=True, type=str)

parser.add_argument("-min_trim_size", "-s", default=75,
                    help="Minimum number of bases to trim from both ends.",
                    type=int)

parser.add_argument("-trim_fraction", "-f", default=1, type=float, nargs='+',
                    help="Fraction of bases to retain from center of gene. Set\
                    to 1 to use all bases minus minimal terminal cutoff. Provide\
                    one or more numbers between 0 and 1, separated by spaces.")

parser.add_argument("-basecov_text", "-b", default="basecov",
                    help="Text string used to indicate basecov files if a directory is given as the input for -basecov.",
                    type=str)

parser.add_argument("-exclude_gene", "-e", default="trimmed_18S_MLSG_ trimmed_28S_MLSG_ trimmed_COI_MLSG_",
                    help="Optional list of gene names to ignore.",
                    type=str)

parser.add_argument("-iqr", default=1.5,
                    help="IQR coefficient for excluding outliers. If --exclude-outliers flag not set, this option is ignored",
                    type=float)

parser.add_argument('--print-cov', dest='print_cov', action='store_true', help="Print detailed coverage information")

parser.add_argument('--print-basecov', dest='print_basecov', action='store_true', help="Print long form base coverage")

parser.add_argument('--print-stats', dest='print_stats', action='store_true', help="Suppress stats printout")

parser.add_argument('--inexact-match', dest='inexact_match', action='store_true', help="Allow exclude genes to support inexact matches. Useful for excluding groups of genes containing particular substrings.")

parser.set_defaults(print_stats=False, print_cov=False, print_basecov=False, inexact_match=False)

# def check_fraction(value):
#     fvalue = float(value)
#     if fvalue > 1 or fvalue < 0:
#         raise argparse.ArgumentTypeError("%s is an invalid fraction value. Enter a number between 0 and 1" % fvalue)
#     return fvalue


# parser.add_argument("-trim_fraction", "-f", default=1, type=check_fraction,
#                     help="Fraction of bases to retain from center of gene. Set\
#                     to 1 to use all bases minus minimal terminal cutoff")

if len(sys.argv[1:])==0:
    parser.print_help()
    # parser.print_usage() # for just the usage line
    parser.exit()


in_basecov = args.basecov
trim_size = args.min_trim_size
ignore_gene = args.exclude_gene
trim_fraction = args.trim_fraction

inexact_match = args.inexact_match
print_trim_covs = args.print_cov
print_basecov = args.print_basecov
print_stats = args.print_stats
basecov_text = args.basecov_text
iqr_coef = args.iqr

perbase = True


def exclude_gene():
    '''Takes a list of gene names, separated by whitespace, and returns them as a list'''
    # If ignore_gene list is empty, return None
    if ignore_gene is None:
        exclude_genes = [None]
        return exclude_genes
    ignore_genes = ignore_gene.split()
    exclude_genes = []
    # Populate list of excluded genes
    if ignore_genes is not None:
        for gene in ignore_genes:
            exclude_genes.append(gene)
    return exclude_genes


def find_length_outliers(pos_dict):
    len_list = []
    for key in pos_dict:
        len_list.append(len(pos_dict[key]))
    len_list = np.array(len_list)
    iqr_screen(len_list, "", "Length")
    q1, q3 = np.percentile(len_list, [25,75])
    len_cut = q3
    return len_cut, np.median(len_list)


def populate_and_rename(in_file):
    '''Opens basecov file and populates dict of genes and coverages,
    and renames, if necessary'''

    exclude_genes = exclude_gene()
    pos_dict = collections.OrderedDict()

    with open(in_file, 'r') as f:
        for line in f:
            include = True
            # Ignore header line
            if "#" not in line:
                gene_id, pos, cov = line.strip().split()
                # Rename gene_id if found in tname_dict
                cov = int(cov)
                if gene_id not in pos_dict and gene_id not in exclude_genes:
                    for exc_gene in exclude_genes:
                        if exc_gene.strip() in gene_id.strip():
                            if inexact_match is True:
                                include = False
                    if include is True:
                        pos_dict[gene_id] = [cov]
                        #print gene_id

                elif gene_id in pos_dict and gene_id not in exclude_genes and include is True:
                    for exc_gene in exclude_genes:
                        if exc_gene.strip() in gene_id.strip():
                            print(exc_gene, gene_id)
                        else:
                            pos_dict[gene_id].append(cov)

    return pos_dict


def trim(covs_in, gene_id, trim_size, trim_fract):
    """Main trimming function. Takes trim parameters and returns a trimmed 
    internal window for each gene."""
    

    covs = covs_in[:]
    cov_len = len(covs_in)
    trim_covs = covs_in[trim_size:len(covs)-trim_size]
    trim_covs_len = len(covs_in)
    trim_window = int(round(trim_covs_len*(1-trim_fract)))

    left_clip = []
    right_clip = []
    if trim_window/2 <= trim_size:
        for i in range(0, trim_size):
            left_clip.append(covs[i])
            right_clip.append(covs[-(i+1)])

            covs[i] = "*"
            covs[-(i+1)] = "*"
    else:
        for i in range(0, int(trim_window/2)):
            left_clip.append(covs[i])
            right_clip.append(covs[-(i+1)])
            covs[i] = "*"
            covs[-(i+1)] = "*"

    if print_basecov is True: 
        pos = 0
        for cov in covs_in:
            print("{}\t{}\t{}".format(gene_id, pos, cov))
            pos = pos+1

        for tcov in covs:
            print("{}\t{}\t{}".format(gene_id+"_trimmed", pos, tcov))
            pos = pos+1


    window_covs = [y for y in covs if y is not "*"]
    win_mean = np.mean(window_covs)

    if print_trim_covs is True:
        left_mean = np.mean(left_clip)
        right_mean = np.mean(right_clip)
        print(gene_id)
        print("\tTrimmed {}\tLen: {} Center: {} Fraction: {} Trim Window: {} Rep Count: {}".format(gene_id, cov_len, cov_len/2, trim_fract, trim_window, covs.count("*"))) 
        print("\tMeans Window: {} Left: {} Right: {}".format(win_mean, left_mean, right_mean))


    return window_covs


def print_pos_varpar(in_file, trim_sizes, trim_fracts):
    # trim_size = args.min_trim_size
    # trim_fraction = args.trim_fraction

    entry_list = []
    pos_dict = populate_and_rename(in_file)
    
    header_list = ["#ID", "Length_Init", "Cov_Init", "PerBase_Init"]
    for temp_trim_size in trim_sizes:
        for temp_trim_fract in trim_fracts:
            len_name = "Length_Tr{}Fr{}\tCov_Tr{}Fr{}\tPerBase_Tr{}Fr{}".format(temp_trim_size, temp_trim_fract, temp_trim_size, temp_trim_fract, temp_trim_size, temp_trim_fract)
            len_name = len_name.split("\t")
            header_list.extend(len_name)
    for key in pos_dict:
        covs = pos_dict[key]
        raw_mean_cov = np.mean(covs)
        init_entry = [key, str(len(covs)), str(raw_mean_cov), str(len(covs)*raw_mean_cov)]
        entry_list.append(init_entry)
        trim_sizes_entries = []
        for temp_trim_size in trim_sizes:
            for temp_trim_fract in trim_fracts:
                trim_size = temp_trim_size
                if len(covs) > trim_size*2:
                    trim_cov = trim(covs, key, trim_size, temp_trim_fract)
                else:
                    trim_cov = [0]
                trim_sizes_entries.append( str(len(trim_cov)) )
                trim_sizes_entries.append( str(np.mean(trim_cov)) )
                trim_sizes_entries.append( str(np.mean(trim_cov)*len(trim_cov) ))

        init_entry.extend(trim_sizes_entries)

    seq_name = in_file.split('/')[-1]

    per_file_entry = []
    per_file_entry.append(header_list)

    if print_stats is True:
        print("\n{} Mapping Statistics".format(seq_name))
        print('\t'.join(header_list))

    for entry in entry_list:
        if print_stats is True:
            print('\t'.join(entry))
        per_file_entry.append(entry)

    per_file_array = np.array(per_file_entry)


    return per_file_array


def print_summary(in_array):
    print("Trim Parameters\tMean")
    for x in in_array.T[1:]:
        col = x[1:]
        col = col.astype('f')
        if "Cov" in x[0]:
            print("{}\t{}".format(x[0], np.mean(col)))


def dir_print_summary(in_array, seq_name):
    ret_list = []

    ret_list.append(["Coverage", seq_name])
    for x in in_array.T[1:]:
        col = x[1:]
        col = col.astype('f')

        if "Cov" in x[0]:
            ret_list.append([x[0], str(np.mean(col).item())])

    if perbase is True:
        for i, x in enumerate(in_array.T[1:]):
            col = x[1:]
            col = col.astype('f')
            if "PerBase" in x[0]:
                len_col = in_array[1:,i-1]
                len_col_int = [ int(y) for y in len_col ]
                perbase_cov = sum(col)/sum(len_col_int)
                ret_list.append( [x[0], str(perbase_cov)])


    return ret_list


def slistdir(directory):
    """os.listdir() that ignores files that
    start with a leading period."""
    filelist = os.listdir(directory)
    return [x for x in filelist
            if not (x.startswith('.'))]


def check_for_fraction():
    for fract in trim_fraction:
        if fract <= 1 and fract >= 0:
            pass
        else:
            raise ValueError('trim_fraction must be between 0 and 1')


def main():
    print("")
    check_for_fraction()
    if os.path.isdir(in_basecov) is True and os.path.isfile(in_basecov) is False:
        print("Processing directory: {}".format(in_basecov))
        cov_summaries = []
        for in_file in slistdir(in_basecov):
            if basecov_text in in_file:
                in_file = os.path.join(in_basecov, in_file)
                seq_name = in_file.split('/')[-1].split("_" + basecov_text)[0]
                output = print_pos_varpar(in_file, [trim_size], trim_fraction)

                temp_covs = dir_print_summary(output, seq_name)
                if any(cov_summaries) is False:
                    for cov in temp_covs:
                        cov_summaries.append(cov)
                else:
                    for i, cov in enumerate(temp_covs):
                        cov_summaries[i].append(str(cov[1]))
        print("\n")
        tp_cov_sum = list(map(list, list(zip(*cov_summaries))))
        for line in tp_cov_sum:
            print("\t".join(line))
    else:
        for in_file in in_basecov.split():
            print("Processing file: {}\n".format(in_basecov))
            seq_name = in_file
            output = print_pos_varpar(in_file, [trim_size], trim_fraction)
            print_summary(output)

    print("\nDone  ")


if __name__ == "__main__":
    main()
```
